# VINE: Variational inference for scalable Bayesian reconstruction of species and cell-lineage phylogenies

**DOI:** 10.64898/2025.12.24.696405

**Authors:** Adam Siepel, Rebecca Hassett, Stephen J. Staklinski

## Abstract

Bayesian methods are now widely used in reconstructing both species and cell-lineage phylogenies, but they remain heavily reliant on computationally intensive Markov chain Monte Carlo sampling. Phylogenetic variational inference (VI) circumvents this dependency but so far has been limited in speed and scalability. Here we introduce Variational Inference with Node Embeddings (VINE), a computational method that combines an embedding of taxa in a high-dimensional space and a distance-based “decoder” with several algorithmic innovations to dramatically improve phylogenetic VI. VINE supports both standard DNA substitution models and CRISPR barcode-mutation models for inference of cell-lineage trees and tissue-migration histories. In extensive simulation experiments, we show that VINE approaches the best available Bayesian methods in both model fit and uncertainty quantification, at speeds orders of magnitude faster. We then apply VINE to ∼1,000 complete SARS-CoV-2 genomes and ∼900 lung-cancer cell barcodes, showing reductions in compute time from days to hours or minutes.

## Introduction

Phylogenetic trees are now ubiquitous across the life sciences, with applications ranging from large-scale comparative and population genomics [1, 2] to charting and combating the spread of infectious diseases [3]. A particularly exciting new frontier for phylogenetics is the reconstruction of cell-lineage trees from CRISPR- based lineage-tracing data [4–7], which can shed important light on tumor evolution [8, 9], developmental biology [10–12], and neurobiology [13, 14]. Datasets for species and cell-lineage trees alike are rapidly growing in size and now often include thousands of taxa, leading to steadily increasing demand for fast and accurate phylogenetic methods.

For many applications, Bayesian inference has become the preferred approach for phylogenetic reconstruction. Bayesian methods naturally quantify the uncertainty about the phylogenetic tree given the data, by inferring a full posterior distribution for the tree topology, the branch lengths, and the parameters governing the substitution process. Since their introduction in the mid 1990s [15–17], these methods have steadily improved, and mature implementations such as MrBayes [18, 19], BEAST/BEAST 2 [20–22], and RevBayes [23] are now extraordinarily widely used. Bayesian approaches are particularly valuable in CRISPR-based lineage tracing, where current datasets often contain limited phylogenetic information, leading to considerable uncertainty in tree reconstruction [12, 24–27].

Despite many recent advances [28], however, Bayesian phylogeny inference methods still lag considerably behind the leading maximum-likelihood tools [29–31] in speed and scalability. They are primarily limited by a reliance on Markov chain Monte Carlo (MCMC) sampling, which explores the full space of tree topologies through a series of discrete rearrangement operations, and inevitably stalls as the number of taxa becomes large. In addition, MCMC-based methods often require tuning of proposal distributions and careful monitoring of convergence, sometimes making them challenging to use for non-experts. Bayesian methods for cell-lineage reconstruction also depend on MCMC [24–27] and therefore face the same bottleneck.

Recognizing the limitations of MCMC, investigators have recently explored a variety of alternative methods for characterizing phylogenetic posterior distributions. Most of these efforts have made use of some form of variational inference (VI), which has been widely employed for approximate inference in other fields [32, 33]. The core idea of VI is, instead of sampling from a high-dimensional posterior distribution, to compel a flexible alternative distribution to approximate the true posterior by minimizing the divergence between distributions (reviewed in [34]). In phylogenetics, the first method of this kind—Variational Bayesian Phylogenetic Inference (VBPI)—was proposed seven years ago by Zhang and Matsen [35] (see also [36, 37]) and subsequently extended to make use of normalizing flows and graph neural networks [38–41], among other features. VBPI has now been followed by several other phylogenetic VI methods including VaiPhy [42], VBPI-Mixtures [43], GeoPhy [44], and Dodonaphy [45] (see also [46–48]). In general, however, research in this area is still at the exploratory stage, and the available methods have yet to achieve the scalability and accuracy needed for applied phylogenetics. To our knowledge, VI has yet to be used in any published phylogenetic analysis beyond benchmarking for methods development.

In this article, we revisit the problem of VI for phylogenetics, with an eye toward applications to both DNA alignments and CRISPR-based lineage-tracing data. We begin with a recently proposed idea [44,45] to embed taxa in a continuous space, decode phylogenies by standard distance-based reconstruction methods, and optimize model parameters by stochastic gradient ascent (SGA). We redesign this method from the ground up, introducing numerous simplifications, extensions, and algorithmic innovations that both improve its accuracy and boost its speed by several orders of magnitude. Our methods are implemented in a freely available computer program, called VINE (Variational Inference with Node Embeddings), that supports a variety of nucleotide substitution models, a recently developed model for CRISPR-barcode editing, and an extension to tissue migration-graph inference, among other features. We show that VINE approaches the best available Bayesian phylogenetic methods in model fit and tree accuracy, at considerably faster speeds. Its variational approximation tends to underestimate posterior uncertainty, but an optional second stage substantially mitigates this problem. Overall, we demonstrate for the first time that variational phylogenetic inference can serve as a practical alternative to mature MCMC-based implementations such as BEAST 2 and MrBayes, enabling approximate Bayesian phylogenetic inference for datasets beyond the reach of current methods.

## Results

### Variational inference using continuous embeddings

The central problem in phylogenetic VI is to approximate the Bayesian posterior distribution of phylogenetic trees given a set of observed genotypes **X** using a flexible variational distribution *q*(*τ,* **b**; ***θ***), where *τ* denotes the *topology* of a phylogenetic tree, **b** is a corresponding vector of *branch lengths*, and ***θ*** denotes the free parameters of the variational distribution. Our particular starting point is a method recently introduced by Mimori and Hamada [44] (see also [45]) that has three essential components. First, the tips of the tree (corresponding to the observed data) are represented by an embedding **x** in a *d*-dimensional continuous space, with an induced matrix of pairwise distances **D**. A simple, flexible multivariate Gaussian sampling distribution, *q*(**x**) = MVN(**x**; ***µ***, **Σ**), is assumed for the embedded taxa. Second, classical distance-based methods for tree reconstruction, such as the neighbor-joining method [49], are leveraged to convert the distance matrix **D** to a tree with branch lengths (*τ,* **b**). In this way, sampled embeddings **x** can be deterministically converted to fully defined phylogenies, permitting calculation of the likelihood of the data by standard methods. Third, the free parameters (***µ***, **Σ**) of the variational distribution *q* are optimized by minimizing the Kullback-Leibler (KL) divergence from the true posterior by stochastic gradient ascent (SGA). As the authors noted, the resulting model can be thought of as a variational autoencoder [50, 51], where the decoder takes the form of the deterministic mapping from embedded taxa to trees.

Despite several appealing properties of this general modeling strategy, initial implementations have not been competitive with well-established MCMC-based methods for Bayesian phylogenetics. We therefore sought to devise new algorithms that would substantially improve performance while maintaining the core ideas of a continuous embedding and distance-based phylogeny decoder. Briefly, our new computational method, VINE, encodes tree topologies and branch lengths together, in one unified continuous embedding, rather than separately; operates in substantially higher-dimensional spaces (*d* = 5 or higher) than previous methods (typically *d* = 2 or *d* = 3); makes use of a new algorithm for efficient backpropagation of gradients through distance-based phylogeny reconstruction algorithms; and permits calculation of the objective function for VI using a fast Taylor approximation instead of the original Monte Carlo approach (**Fig. 1**; full details in **Methods**). With these innovations, we obtain large improvements in efficiency without compromising accuracy in variational Bayesian phylogenetic inference, allowing applications to datasets with 1,000 taxa or more. Finally, we extend VINE to support not only a variety of richly parameterized DNA substitution models but also mutation models for CRISPR-based barcode editing, making it the first phylogenetic VI method to apply to both species and cell-lineage phylogeny reconstruction problems. As we will show, an additional extension allows VINE to be used for inference of tissue-migration graphs based on CRISPR-barcode data (see [27]).

**Figure 1:**
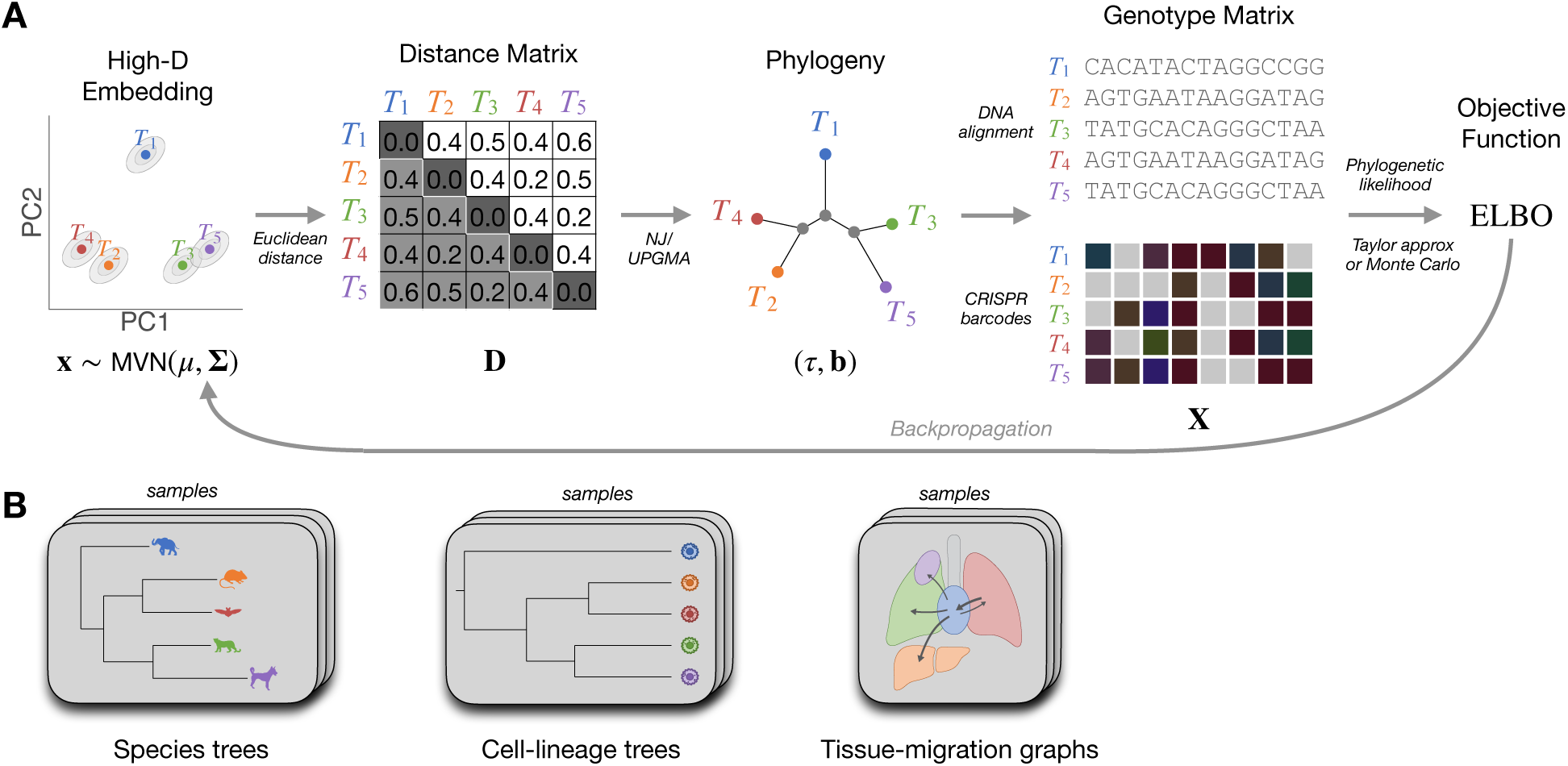
**(A)** Flow of information in VINE from high-dimensional embedding of taxa to Evidence Lower Bound (ELBO) for variational inference, followed by flow in the reverse direction via backpropagation. Each of *n* taxa is represented as a point **x** in a *d*-dimensional space, which is sampled from a multivariate normal distribution, MVN(***µ***, **Σ**). A distance matrix **D** is computed from **x** and then converted by neighbor-joining (NJ) or UPGMA to a tree *τ* with branch lengths **b**, allowing calculation of a phylogenetic likelihood from a genotype matrix **X**, which may be a DNA alignment or a CRISPR-barcode mutation matrix. Finally, the ELBO is computed using either a Taylor approximation or Monte Carlo sampling. An approximate gradient of the ELBO with respect to (***µ***, **Σ**) is computed by backpropagation through the component transformations, allowing for efficient optimization by stochastic gradient ascent (SGA). **(B)** Three applications of interest: inference of species trees (*left*), cell-lineage trees (*middle*), or tissue-migration graphs (*right*). In each case, samples from the approximate posterior distribution are obtained by sampling values of **x** from the optimized MVN distribution and transforming them as in (A).

### VINE is comparable to MCMC in model fit and offers substantially improved speed

We first evaluated VINE’s performance against simulated DNA sequence data, where the true evolutionary history is known and model misspecification can be controlled. We simulated phylogenies under a birth-death model, followed by subsampling and rescaling, to achieve branching patterns and overall tree scales roughly similar to those of real phylogenies for mammals, and then simulated DNA alignments by allowing nucleotides to evolve along these trees under the HKY substitution model [52] (see **Methods**). We generated trees for *n* = 10 to *n* = 1000 taxa, with ten replicates each, and we produced both short alignments of *L* = 300 bp and longer ones of *L* = 10, 000 bp, with the shorter alignments designed to permit faster phylogenetic reconstruction but with more statistical uncertainty.

In our benchmarking experiments, we compared VINE to three recently developed VI methods—VaiPhy [42], GeoPhy [44] and Dodonaphy [45]—as well as to two of the leading MCMC-based Bayesian phylogenetic inference packages, MrBayes [19] and BEAST 2 [21, 22], both of which we tested with and without the use of BEAGLE for acceleration [53]. Notably, GeoPhy and Dodonaphy are the two previous methods to use a similar embedding scheme to the one we adopted, although unlike VINE, both rely heavily on hyperbolic geometries (see **Discussion**). We also tried to include VBPI-GNN [39] in our experiments but found that it was not sufficiently fast for practical consideration. Because the previously published VI methods all assume the simple Jukes-Cantor (JC) substitution model [54], we started by assuming that model for inference with all methods. As a baseline, we also reconstructed trees using a straightforward neighbor-joining implementation with no further optimization.

Across all simulated datasets, we found that VINE was able to obtain reconstructed phylogenetic models that fit the data well. As would be expected for a likelihood-based method, its maximized log likelihoods were reliably higher than the log likelihoods of the true (generating) models at all dataset sizes (**Fig. 2A**). Inspection of individual trees showed that the reconstructions were generally close to the ground truth, with occasional errors that tended to correspond to difficult-to-resolve features of trees (**Fig. 2C&D**). VINE’s performance by these measures was very close to that of MrBayes and BEAST 2 (**Fig. 2A**). The previous VI methods had slightly lower log likelihoods, with GeoPhy and VaiPhy obtaining values close to those of the true models, and Dodonaphy showing somewhat poorer performance (**Fig. 2A**).

**Figure 2:**
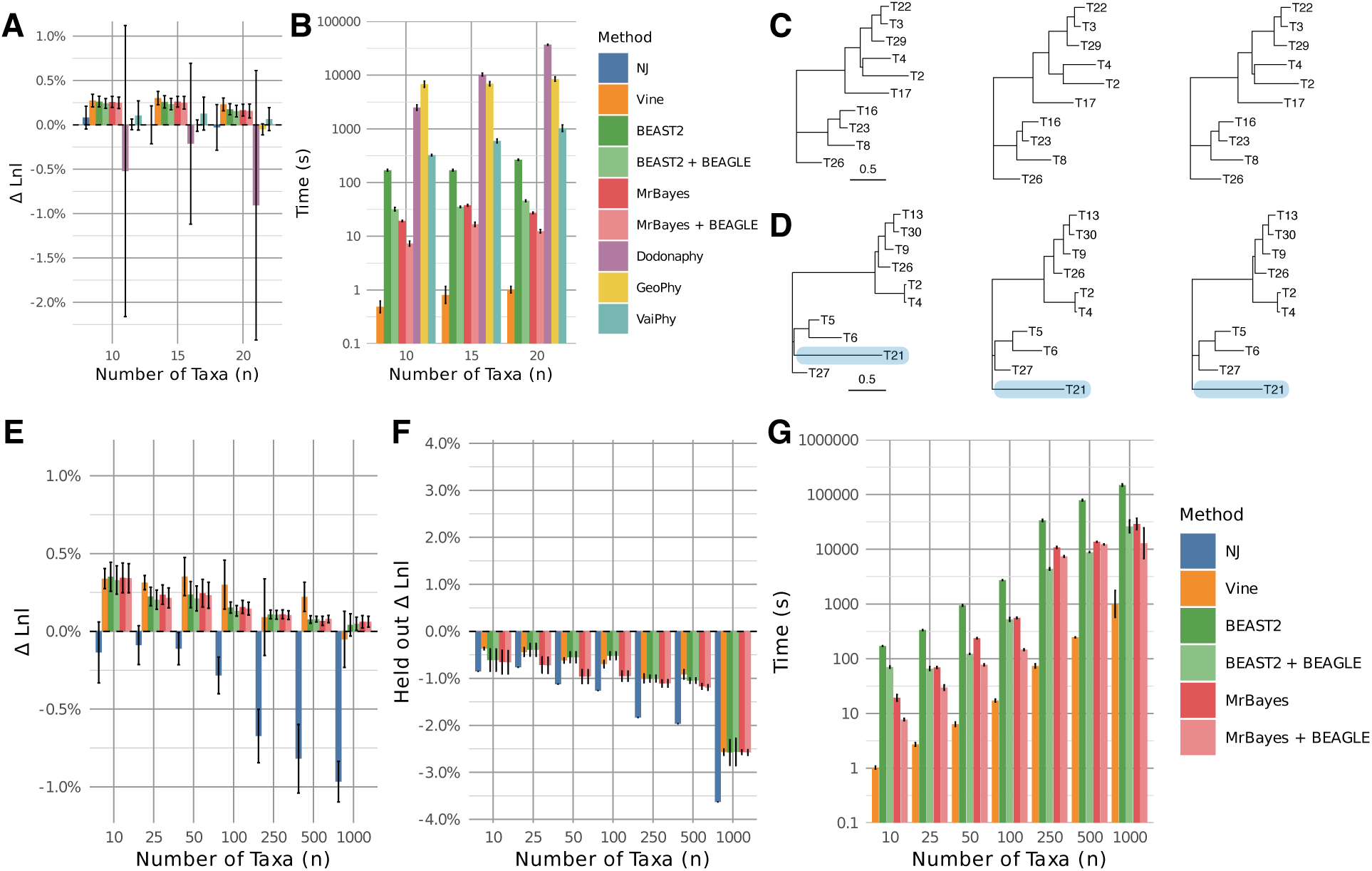
Performance of VI and MCMC-based methods on simulated DNA datasets. **(A)** Maximized log likelihood during model fitting, relative to the log likelihood of the true (generating) model (zero line), for small numbers of taxa (*n* ≤ 20) under the Jukes-Cantor substitution model. **(B)** Compute time required when running without parallelization or GPU acceleration on an HPE ProLiant DL380 Gen10 server (see **Methods**), in seconds per replicate (note log scale). **(C)** Reconstruction of a simulated 10-taxon tree (*left*) by VINE (*center*) and BEAST 2 (*right*). **(D)** A second example with a reconstruction error. Horizontal branch lengths are drawn to scale in substitutions per site. For VINE and BEAST 2, samples from the posterior distributions are summarized by maximum-clade-credibility (MCC) trees (using TreeAnnotator [22]). **(E)** Maximized log likelihood relative to the true model for larger numbers of taxa (up to *n* = 1000) under the HKY substitution model. **(F)** Average log likelihood across posterior samples for held-out data, also relative to the true model. **(G)** Compute time required. Results shown are for 300 bp alignments, with ten replicates per bar. Error bars represent one standard deviation. VINE (*orange*) was applied with default parameters (a Euclidean geometry, the Taylor approximation, and the CONST variance parameterization; see **Methods**). Dimensionality varied from *d* = 5 for *n* = 10 to *d* = 10 for *n* = 1000. See additional results in **Supplementary Figs. S1 & S4**).

Because the main goal of VI is speed, we also kept track of the CPU time required for each experiment. Comparisons of running times are complicated, however, by questions of how long to sample with MCMC-based methods as well as differences in multithreading schemes and GPU utilization. To keep our comparisons as straightforward as possible, we ran all methods using a single CPU core and we used measures based on split chains and effective sample sizes to calibrate the number of MCMC iterations in a dataset-dependent manner (see **Methods**). (In all cases, we reported “wall-clock” times for entire program executions, including posterior sampling as well as any I/O or preprocessing performed by each program.) Nevertheless, we found quite stark differences in running times across methods. The previous VI methods, in particular, were highly compute-intensive, requiring from many minutes to several hours for even small alignments (**Fig. 2B**). The fastest of these methods, VaiPhy, took more than five minutes per replicate for our smallest (10-taxon, 300-site) simulated alignments, and seventeen minutes per replicate for similar alignments with 20-taxa. By contrast, in all cases with *n* ≤ 20 taxa and 300 sites, VINE obtained good-quality phylogenies in roughly one second per replicate, showing speedups of *>*500-fold relative to VaiPhy and as much as ∼30,000-fold relative to Dodonaphy.

The MCMC-based methods were much faster than the experimental VI methods but not as fast as VINE. MrBayes required between about 20 and 40 seconds per replicate on these examples, decreasing to 7–17 seconds with the use of BEAGLE (**Fig. 2B**). In these experiments, BEAST 2 was slower than MrBayes by about an order of magnitude, largely owing to slower convergence by our criteria (see **Methods**), although BEAGLE narrowed this gap. Nevertheless, even relative to MrBayes with BEAGLE—the fastest MCMC method—VINE was able to obtain comparable reconstructions with speeds 12–16 times greater. Because running times began to reach many hours per replicate for some of the VI methods, we did not extend these experiments beyond 20 taxa.

To consider datasets of more realistic size, we ran VINE and the two MCMC-based methods on the simulated alignments with up to 1000 taxa. In this case, we assumed the more realistic HKY model for inference, and we evaluated the quality of all reconstructed phylogenies by both likelihood-based measures of model fit and measures of discordance from the true (simulated) trees. We found, again, that the maximized log likelihoods were similar across methods and generally better than those of the true model (**Fig. 2E**, **Supplementary Fig. S1A**). To control for overfitting in model estimation, we also evaluated the average log likelihoods of sampled trees on held-out sequence alignments generated by the same models, and found—again as expected—that both VINE and the MCMC-based methods performed only slightly worse than the true models, with deviations of ∼1% (**Fig. 2F**, **Supplementary Fig. S1B**). By the Robinson-Foulds (RF) measure of topological discordance [55] and the branch-score distance [56], VINE and the MCMC-based methods also performed fairly similarly, although VINE did show some increased discordance from the true trees at larger values of *n* (**Supplementary Fig. S2**). The difference in RF distances appears to be partly a consequence of VINE’s reduced posterior variance, as discussed further below; VINE tends to converge on a small set of tree topologies and therefore is less effective at “hedging its bets” relative to the truth in measures of topological discordance. Both of these measures may also be influenced by VINE’s tendency to slightly underestimate long branch lengths (see below).

The trend with running times remained similar to what we observed with smaller alignments, with VINE showing speedups of one to two orders of magnitude relative to MrBayes+BEAGLE, which, in turn, improved on BEAST 2+BEAGLE by about another 2–3 times, with some variability across sizes (**Fig. 2G**, **Supplementary Fig. S4**). In this case, BEAGLE typically accelerated MrBayes by factors of 2–3 and BEAST 2 by factors of 3–5. VINE was again able to obtain trees for 10 taxa in about a second per replicate, on average, increasing to about 6 seconds for 50 taxa and 17 seconds for 100 taxa. It required about 4 minutes per replicate for 500 taxa and 17 minutes for 1000 taxa. By comparison, MrBayes+BEAGLE exhibited running times of 8 seconds for 10 taxa up to 3.6 hours for 1000 taxa (**Fig. 2G**). A comparison of convergence trajectories indicated that VINE finds similar maxima to MCMC-based methods but converges considerably more quickly (**Supplementary Fig. S3**). With longer alignments, VINE continued to improve on both MCMC-based methods by an order of magnitude or more (**Supplementary Fig. S4**). We also examined alignments simulated under the more complex general time reversible model with gamma-distributed rate variation (GTR+G) [57, 58], performing inference under the same model, and found that the same trends held, with VINE producing similar maximized log likelihoods to BEAST 2 and MrBayes but with substantially faster run times (**Supplementary Fig. S5**). Overall, these experiments demonstrate that VINE is the first variational phylogenetic inference method to offer both comparable model-fitting performance to MCMC-based methods and significant improvements in speed (see **Discussion**).

### Performance improves with the dimensionality of the embedding space

We noticed that model fit seemed to be quite sensitive to the dimensionality *d* of the embedding space, more than to the use of a hyperbolic geometry rather than a Euclidean one. We therefore carried out a series of experiments where we evaluated the impact of the dimensionality *d* on both goodness of fit and running time, again using simulated data. We focused on datasets with *n* = 25 and *n* = 50 taxa and considered values of *d* between 2 and 8 under both the Euclidean and hyperbolic embedding schemes.

We found, under the Euclidean embedding, that the maximized log likelihood was poor with *d* = 2 and *d* = 3 (e.g., ∼200 units lower that for the true model for *n* = 25 and ∼800 units lower for *n* = 50 taxa; see **Fig. 3A**, **Supplementary Fig. S6**). As *d* increased, however, the log likelihood rapidly improved from *d* = 2 to *d* = 3, tapering off at about *d* = 4. We expected that this improvement in model fit would come with a time penalty, since the number of free parameters in the model is roughly equal to *nd* (see **Methods**). Interestingly, however, we found that the running time slightly *decreased* with *d* rather than increasing, owing to more efficient convergence of the optimization algorithm. For example, with *n* = 25 taxa, the average time to convergence decreased from about 3.8 seconds for *d* = 2 to about 3.2 seconds at *d* = 4 and then reached a minimum of 2.5 seconds by *d* = 5. Similarly, with *n* = 50 taxa, the time to convergence was cut in half from *d* = 2 to *d* = 6.

**Figure 3:**
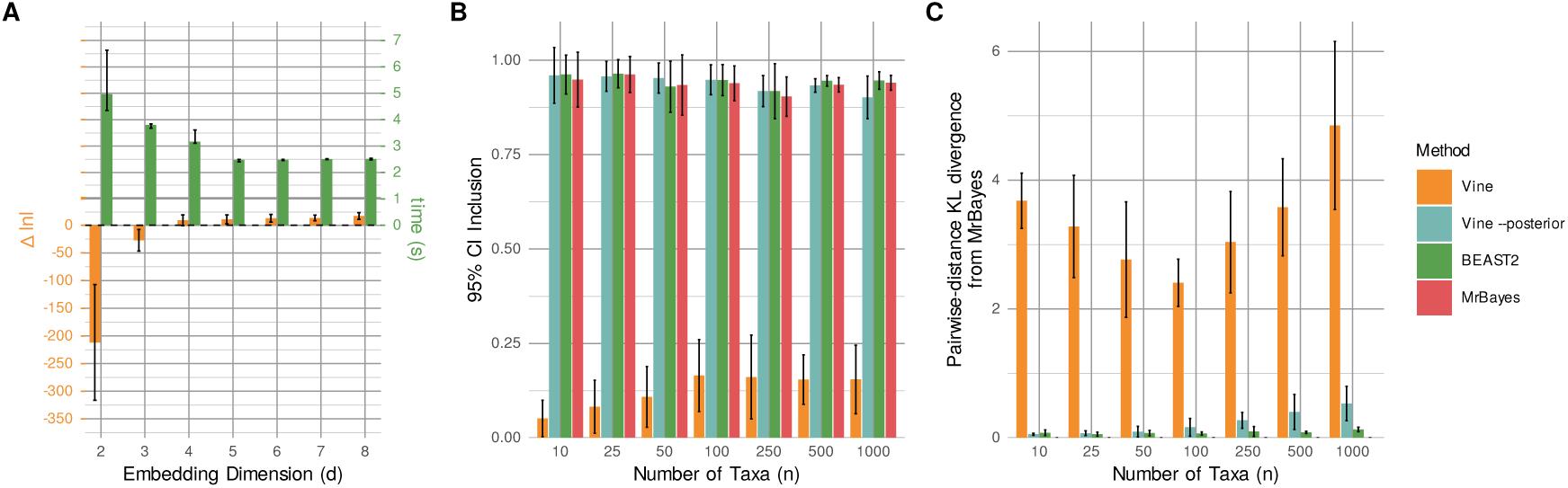
**(A)** Performance improvement of VINE with increasing dimensionality *d* of the embedding space. Shown are running times per replicate (*green*) and deviations of the maximized log likelihood from that of the true model (*orange*). Results are for *n* = 25 taxa, alignments of 300 bp, and estimation under the HKY model with the default variance parameterization and a Euclidean geometry (see also **Supplementary Fig. S6)**. **(B)** Accuracy of posterior distributions, as measured by the fraction of all pairwise distances between taxa that fall within the estimated 95% credible interval (see also **Supplementary Fig. S7**). **(C)** Average marginal distance-KL discrepancy of VINE’s approximation of the posterior distribution from that inferred by MrBayes, computed from the marginal distributions of pairwise distances (**Methods**; see also **Supplementary Fig. S9**). In (B) and (C), results are shown for simulated trees with various numbers of taxa for VINE with default parameters (VINE) or two-stage estimation of posterior uncertainty (VINE `--posterior`; see **Methods**), for BEAST 2, and for MrBayes (not shown in (C)). Bars represent averages over ten replicates.

Under the hyperbolic embedding scheme, the log likelihood was similarly sensitive to the embedding dimension, but the running time behaved in a less predictable manner (**Supplementary Fig. S6**). In addition, the running time was generally greater, and with higher variance across replicates, owing to delays in convergence of the optimizer.

Overall we found that, once SGA is suitably tuned (see **Methods**), it is capable of optimizing thousands of parameters in a highly efficient manner, and there seems to be little cost, and substantial benefit, to operating at values of *d* = 8 or higher. At the same time, we found no reason to prefer the more complex hyperbolic geometry over a simple Euclidean scheme for embedding. At least in our hands, any advantages in flexibility from the hyperbolic geometry are offset by difficulties in fitting the model (see **Discussion**).

### Capturing the full posterior variance remains challenging

We observed in our experiments with simulated data that the variance of VINE’s approximate posterior distribution was difficult to estimate accurately. During optimization, VINE down-weights the log-determinant term in the ELBO, which is equivalent to targeting a tempered posterior distribution (see **Methods**). Without this tempering, we found that the approximate posterior was strongly overdispersed and the expected log likelihood was substantially reduced. With tempering, however, the variance tended to collapse during optimization. For example, under the CONST parameterization, which assumes a simple diagonal covariance structure with a single free parameter (**Σ** = *e^η^***I**), the *η* parameter was driven toward strongly negative values and approached the floor we had set for it (such that *e^η^*= 1 × 10^−3^). In our experiments, we were unable to find a single formulation of the objective function that yielded both high likelihoods and accurate estimates of posterior uncertainty.

We first experimented with various alternative parameterizations of the covariance matrix, with the goal of capturing more of the structure of the true posterior. We introduced a fully parameterized diagonal covariance matrix (DIAG), a covariance matrix proportional to a double-centered version of the initial distance matrix (DIST), and a general low-rank parameterization of the covariance matrix (LOWR) (see **Methods** for details). None of these richer parameterizations, however, enabled accurate estimation of the posterior variance. Difficulties in variance estimation are known to occur in VI when the approximating distribution has insufficient flexibility to capture the structure of the true posterior (see **Discussion**).

We therefore adopted a two-stage strategy for applications in which an improved characterization of posterior uncertainty is desired (`--posterior` option). In the first stage, VINE optimizes the tempered objective, yielding the same variational estimates as usual. But then it carries out a second stage, in which the posterior is approximated locally around the resulting point estimate, using a Laplace approximation for branch lengths and likelihood-weighted nearest-neighbor interchanges for the topology (see **Methods**). This second stage considerably improves the characterization of posterior uncertainty at no net increase in computational cost.

We evaluated these strategies by carrying out experiments with simulated DNA alignments of various sizes, across combinations of our four covariance parameterizations, with and without the second stage of estimation. In each case, we allowed VINE and BEAST 2 to estimate approximate posterior distributions from the same ten simulated alignments, and we considered not only the quality of the model fit, but also the variance of the posterior. In assessing these posterior distributions, we focused on two measures: (1) the fraction of 95% credible intervals (CIs) for all *^n^* pairwise distances between taxa along the reconstructed trees that contained the true value used in simulation; and (2) the average Kullback–Leibler (KL) divergence between marginal summaries of the approximate posterior distribution and those of a high-quality MCMC-based estimate, which we computed either from distributions of pairwise distances or frequencies of splits (see **Methods**). Notably, these KL measures should be regarded as cross-method posterior discrepancies, not KL divergences between full tree posteriors, both because they reflect marginal summaries of the full posterior and because the methods differ in their priors and treatment of substitution parameters.

Overall, we found that, indeed, the posterior variance was poorly characterized in VINE’s default mode (CONST parameterization, Euclidean geometry, and no second stage). In this case, while measures of model fit, tree accuracy, and running time remained favorable (**Supplementary Fig. S7**), the 95% CIs were quite narrow and fewer than 20% of true values were contained within them (**Fig. 3B**). By contrast, BEAST 2 performed exceptionally well by this measure, with 92–97% inclusion of true values across all values of *n*. MrBayes showed similar accuracy to BEAST 2. Notably, the posterior mean estimates of pairwise distances were close to the truth for all methods, although VINE did exhibit a tendency to slightly underestimate the largest distances (**Supplementary Fig. S8**). Consistent with its narrow 95% CIs, VINE’s approximate posteriors showed substantially elevated marginal KL discrepancies from the MCMC-based reference distributions (**Fig. 3C**, **Supplementary Figs. S9&S10**).

As intended, the two-stage procedure did substantially improve VINE’s characterization of the posterior variance. This procedure was effective under CONST and was not enhanced by the other parameterizations. With `--posterior`, the 95% CI inclusion increased to 90–96% across tree sizes, comparable to that of the MCMC-based methods, and the average marginal distance-KL discrepancy was reduced by one to two orders of magnitude (**Fig. 3B&C**). The average marginal split-KL discrepancy, however, improved only modestly (e.g., 0.14 vs. 0.24 at *n* = 100 and 0.35 vs. 0.41 at *n* = 1000) and remained several-fold higher than the discrepancy between BEAST 2 and MrBayes (**Supplementary Fig. S9**). In summary, the two-stage procedure allows VINE to capture most of the posterior uncertainty in branch lengths at no additional computational cost, but, because it explores tree space only locally, it captures topological uncertainty less completely than MCMC-based methods (see **Discussion**). In addition, this procedure does have costs in sample-averaged measures of model fit and tree accuracy (**Supplementary Fig. S7B–D**). We therefore retain the single-stage procedure as the default in VINE and recommend the use of `--posterior` only when improved measures of posterior uncertainty are needed.

### VINE shows top performance in cell-lineage phylogeny inference

As noted, a major area of interest in phylogeny inference today is the reconstruction of cell-lineage trees from CRISPR-based barcoding data. This problem is closely related to standard phylogeny inference, but certain unusual features of the CRISPR-based editing process—which tends to produce insertions and deletions (indels) rather than point mutations, and which can be “silenced” in a site-specific manner if target sites are disrupted—have led to the development of customized mutation models [24, 31, 59–61]. It has been shown that incorporation of these models into likelihood-based phylogenetic inference methods can lead to substantial improvements in reconstructed trees [31].

We therefore extended VINE to support the mutation model recently used in the LAML (Lineage Analysis via Maximum Likelihood) program [31] (see also [24])—a continuous-time Markov model that captures the irreversibility of CRISPR-mediated indels, site-specific mutation rates, barcode silencing, and missing data (available via the `-i CRISPR` option in VINE; see **Methods**). We ensured that VINE produces time-resolved ultrametric trees (with all tips equidistant from the root) similar to those inferred by LAML, by using the UPGMA (Unweighted Pair Group Method with Arithmetic Mean) algorithm [62] in place of neighbor-joining in CRISPR mode (see **Methods**). We then benchmarked VINE against LAML on simulated CRISPR barcode mutation matrices. We also evaluated our recently developed MCMC-based method BEAM (Bayesian Evolutionary Analysis of Metastasis), which is primarily designed for inference of tissue migration graphs in metastasis but can also be used for cell-lineage-tree reconstruction [27]. Several other tree-reconstruction programs are available for this problem, including Cassiopeia [59], TiDeTree [24], and Startle [63], but LAML and BEAM appear to be the best-performing methods at present [27, 31], so we focused on them for our comparisons.

As with our DNA alignments, we began by simulating trees and CRISPR barcode mutation matrices, taking advantage of the barcode-editing simulation tools in the Cassiopeia package and approximately matching mutation patterns observed in real data (see **Methods**). We assumed a barcode cassette with ten arrays of three target sites for a total of 30 editing sites per cell, and we generated 10 simulation replicates for trees with 10, 25, 50, 100, 250, 500, and 1000 taxa. We then applied VINE, LAML, and BEAM to each simulated alignment and recorded measures of model fit and tree discordance. Before comparing the reported likelihoods, we verified that the three programs returned exactly the same values (up to numerical precision) for the same mutation matrices and trees.

We found that VINE was able to obtain trees of similar log likelihood to those reported by LAML and BEAM across all simulated datasets (**Fig. 4A** and **Supplementary Fig. S11**). The maximized log likelihoods differed by less than 1% on average at small *n* and were nearly identical (within ∼0.1%) at *n* = 1000. These average differences were small relative to the standard deviation across replicates and were not statistically significant. The topological distances of the inferred trees from the true ones were also similar under all three methods, although, as noted for the DNA simulations, VINE did show slightly elevated measures of discordance at large *n* (**Supplementary Fig. S12**). We also found, as in the DNA case, that VINE’s default approximation of the posterior distribution was substantially under-dispersed relative to the version inferred by BEAM, but that it was much improved with `--posterior` (**Supplementary Fig. S13**).

**Figure 4:**
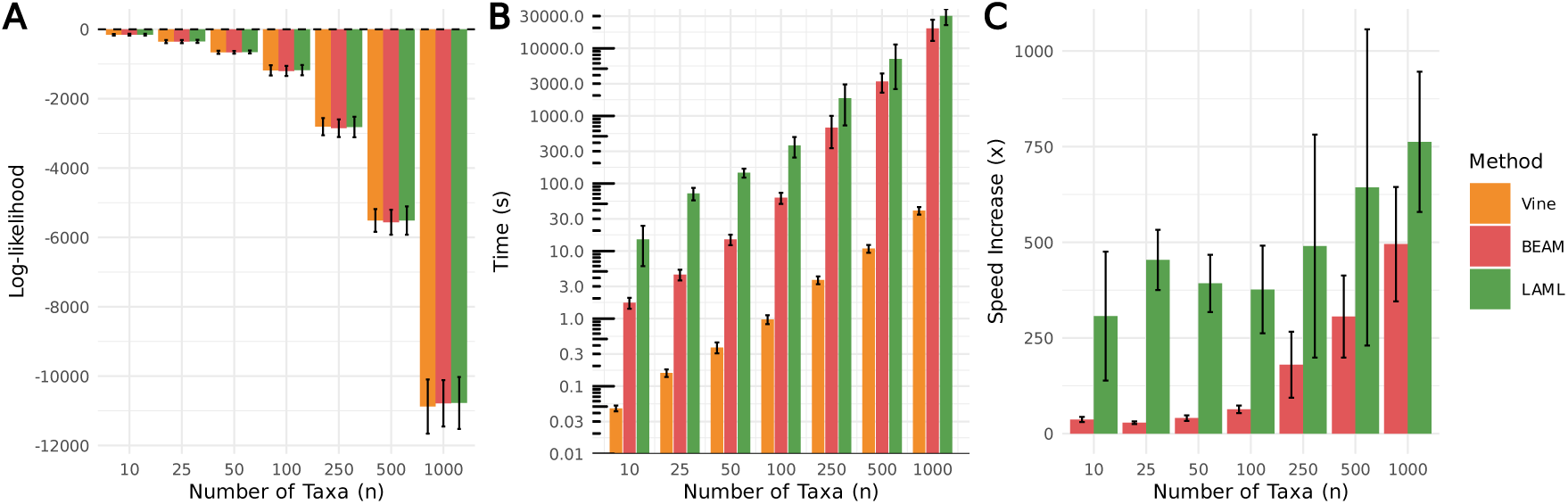
Comparison of VINE with LAML [31] and BEAM [27] on simulated CRISPR-barcoding data, for various numbers of taxa *n*. **(A)** Maximized log likelihood during model fitting. **(B)** Compute time per replicate. **(C)** Speed increase of VINE relative to LAML and BEAM. Results are for 10 simulated datasets for each value of *n* with 30 barcode sites and editing parameters based on real data [67] (see **Methods**). Error bars represent one standard deviation.

As with the DNA models, VINE was able to achieve these generally high levels of accuracy at considerably greater speed than the other methods (**Fig. 4B**). Running times per replicate ranged from a fraction of a second for *n* = 10 to about 40 seconds for *n* = 1000. VINE was roughly 300–750 times faster than LAML, with greater speed advantages for larger *n* (**Fig. 4C**). Interestingly, the MCMC-based BEAM method was also considerably faster than LAML in these benchmarks, particularly for smaller numbers of taxa (e.g., by about 10-fold for *n* ≤ 50). Nevertheless, VINE still improved on BEAM by orders of magnitude across benchmarks. Notably, the other likelihood-based methods available for this problem, such as TiDeTree [24], are considerably slower than LAML, so to our knowledge, VINE is now the fastest such method available by a substantial margin. Overall, VINE is able to fit CRISPR barcoding data at least as well as competing methods, at considerably greater speed, and (like BEAM) it does so by producing an approximate posterior distribution rather than a single point estimate—a feature that can be particularly important in cell-lineage reconstruction, where there is often a great deal of uncertainty in tree inference (see **Discussion**).

### Applications to real DNA data

To demonstrate the applicability of VINE to real DNA data, we first applied it to a collection of eight alignments originally assembled by Lakner et al. [64], which contain various types of nucleic-acid sequence data (DNA, rRNA, rDNA, and mtDNA) for between 27 and 59 taxa (median 42) and range in length from 378 to 2520 (median 1736) sites (**Supplementary Table S1**). These alignments are typical of modest-sized datasets frequently analyzed in applied phylogenetics and have been widely used in recent benchmarking studies (e.g., [44, 45, 65]). We again compared VINE with BEAST 2 and MrBayes using the HKY substitution model, omitting the other VI methods owing to their long running times. As expected from our simulation experiments, all methods were comparable in model fit, with VINE achieving maximized log likelihoods within 0.4% of those of BEAST 2 and MrBayes on average (**Supplementary Fig. S14A**). On visual inspection, the reconstructed trees were generally similar under all reconstruction methods, with some minor differences at difficult-to-resolve branchings (**Supplementary Fig. S15**). As with the simulated data, however, the running times for VINE were substantially reduced, by average factors of 9.2 relative to Mr-Bayes+BEAGLE and 17.8 relative to BEAST 2+BEAGLE (**Supplementary Fig. S14B**). Running times for VINE ranged from 4–20 seconds, in comparison to 46–204 seconds for MrBayes+BEAGLE and 112–293 seconds for BEAST 2+BEAGLE.

We then examined a larger dataset more representative of modern applications in Bayesian phylogenetics. Inspired by recent widespread interest in the use of phylogenetic methods to study the SARS-CoV-2 pandemic [3], we obtained the latest SARS-CoV-2 whole genome sequences from Nextstrain [66] (downloaded February 25, 2026), consisting of ∼74,000 genomes after data-quality filtering, and extracted two subsets by stratified random sampling: a large subset of 1060 genomes and a smaller subset of 364 genomes, each comprising ∼30k nucleotide sites (see **Methods**). We first ran VINE and BEAST 2 on the 364-taxon subset using the general time reversible (GTR) DNA substitution model and the discrete gamma model for rate variation among sites. The two methods produced broadly similar trees (**Fig. 5A**) in which SARS-CoV2 genomes clearly grouped by collection time, reflecting their evolution as the pandemic progressed. These two trees exhibited highly correlated pairwise distances (**Fig. 5B**) and only minor differences in overall branching patterns (**Fig. 5C**). Notably, however, BEAST 2 required over 22 hours for this dataset, whereas VINE finished in about 13 minutes (in this case, both programs were permitted eight threads on our server). We found general agreement between VINE and BEAST 2 in measures of posterior uncertainty, provided VINE was run using the two-stage `--posterior` option (**Supplementary Fig. S16**).

**Figure 5:**
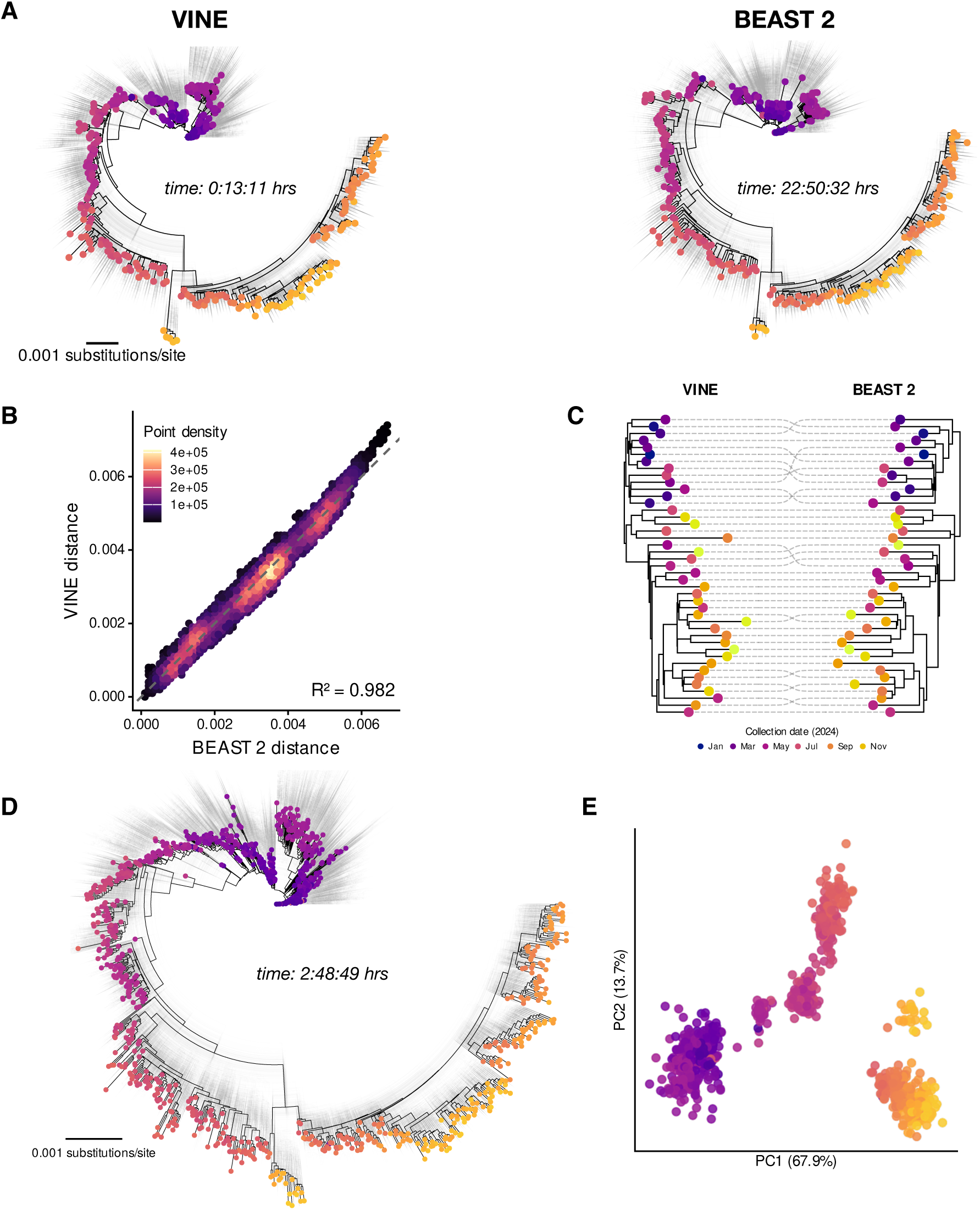
**(A)** Maximum-clade-credibility (MCC) trees inferred by VINE and BEAST 2 for 364 randomly selected SARS-CoV-2 genomes from Nextstrain with tips colored by collection date. Clouds of trees in gray represent posterior samples. **(B)** BEAST 2 vs. VINE posterior mean estimates of pairwise distances in substitutions per site for all pairs of taxa in (A). **(C)** Aligned MCC subtrees for samples from 2024, showing broad topological agreement with minor differences. **(D)** MCC tree and posterior cloud inferred by VINE for a larger 1060-taxon SARS-CoV-2 dataset. **(E)** First two principal components for the embedding learned by VINE for the tree in (D).

Next we applied both methods to the larger 1060-taxon dataset. VINE was able to complete this analysis in under three hours (again with eight threads), producing a tree with a similar overall structure to the smaller one but with about three times as many tips (**Fig. 5D**). We inspected the high-dimensional embedding learned by VINE and found, interestingly, that the overall correlation structure of these SARS-CoV-2 genomes was directly apparent from its first two principal components (**Fig. 5E**). By contrast, BEAST 2 struggled to converge on this larger dataset, and we terminated the run after about three days of processing. Overall, this example demonstrates that VINE is capable of carrying out state-of-the-art variational Bayesian phylogenetic analyses of large modern datasets, with similar point estimates to MCMC-based methods but with considerably better scaling properties, although the two-stage `--posterior` option was required for accurate estimates of posterior uncertainty.

### Applications to real CRISPR data

To demonstrate applicability to real data for cell-lineage tree reconstruction, we focused on a dataset based on a lung-cancer xenograft mouse model that contains 83 clonal populations (CPs) ranging in size from *>*11,000 to ∼30 cells [67]. Larger datasets now exist, but this one includes particularly rich mutational data for a broad range of CPs and it has been widely analyzed. For our comparative purposes, we excluded CPs 1–3, which are prohibitively large (*>*5000 cells) for LAML, leaving 80 CPs that ranged in size (after removing duplicates) from 26 to 899 cells (median 63 cells). As above, we compared VINE with LAML on these data, assuming a uniform prior distribution for mutation rates and allowing for a free silencing-rate parameter (see **Methods**).

On this dataset, VINE obtained somewhat higher log likelihood values than LAML on average (by 20.2%), with some variability across clonal populations (**Supplementary Fig. S17A**). As with the simulated data, VINE and LAML behaved similarly for the smaller trees but VINE often significantly outperformed LAML on the larger ones. Nevertheless, VINE was faster than LAML by orders of magnitude, with an average speed-up of 5663-fold (**Supplementary Fig. S17B**). The average running time for these CPs was 8 seconds for VINE in comparison to 12.7 hours for LAML. The largest clone took over five days with LAML and only 99 seconds with VINE.

Beyond inference of cell-lineage trees, phylogenetic methods have recently been adapted to reconstruct the spread of cancer cells across tissues [68–71] to reveal the rates, routes, and molecular changes associated with metastasis [67, 72–74]. Our recent method BEAM [27] is the first to simultaneously reconstruct both a cell-lineage phylogeny and a tissue-migration graph in a fully Bayesian manner. BEAM models the barcode mutation and tissue migration processes using conditionally independent continuous-time Markov chains, and samples from the joint posterior distribution of lineage trees and tissue labels. The method shows excellent performance in migration-graph reconstruction but requires MCMC for inference (using BEAST 2), and is limited in scalability to a few hundred cells. To address these limits in scalability, we extended VINE to accept tissue labels for cells and support BEAM’s tissue-migration model during inference. This extension required only changes to the likelihood and gradient calculations, as well as support for estimation of migration rates as nuisance parameters in SGA (see **Methods**).

To validate this approach, we benchmarked VINE in migration mode against BEAM, Metient [70], and MACH2 [71], using our recently described simulation framework [27]. We found, indeed, that VINE’s migration model produced reconstructions of simulated migration graphs more accurate than those from Metient and MACH2, and nearly as accurate as those from BEAM (**Fig. 6A**, **Supplementary Fig. S18A**), but with speeds orders of magnitude faster (**Fig. 6B**, **Supplementary Fig. S18B**).

**Figure 6:**
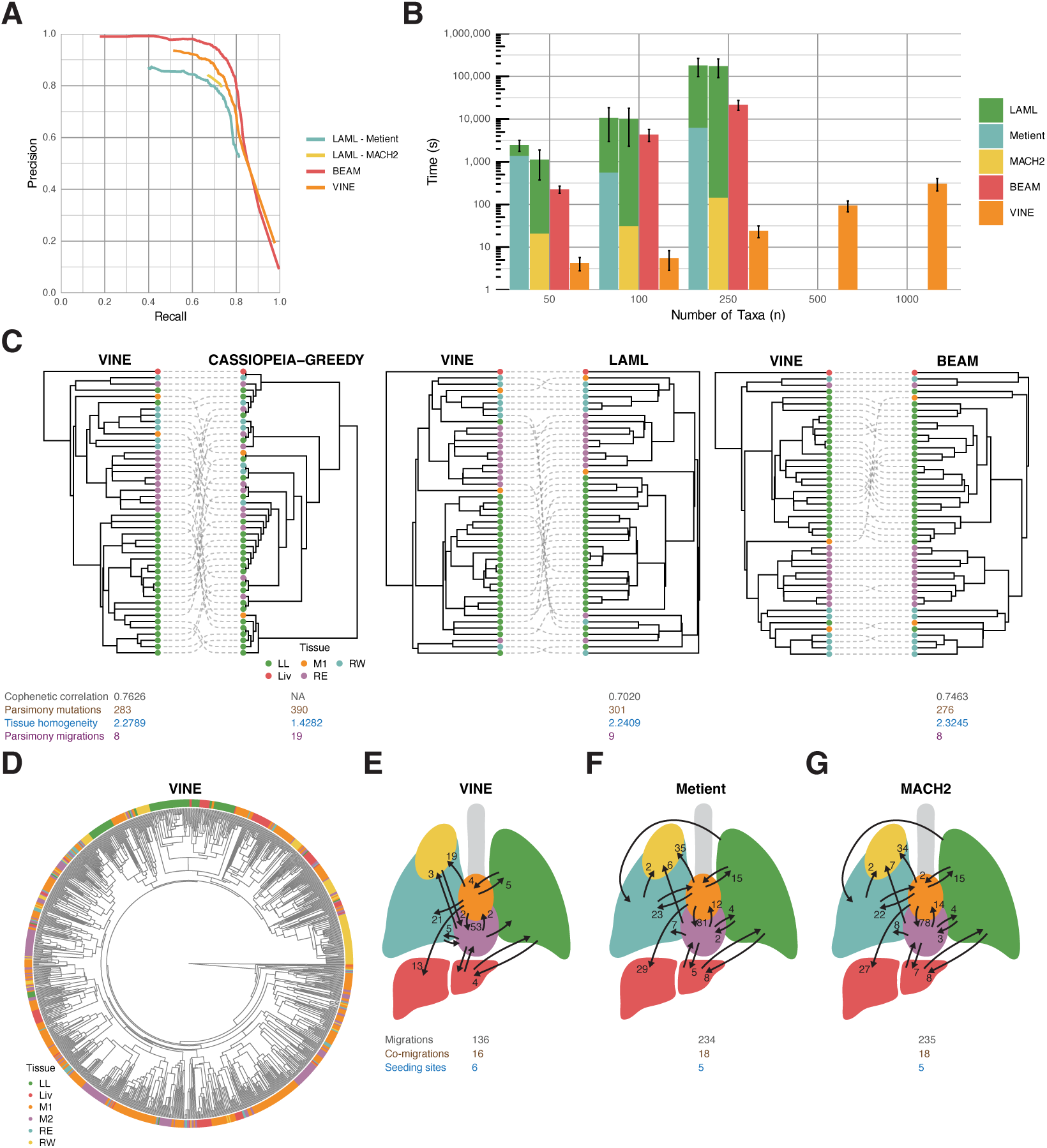
Results from running VINE in migration mode on CRISPR-based lineage-tracing data. **(A)** Precision vs. recall for individual edges of simulated tissue-migration graphs (as detailed in [27]) relative to Metient [70], MACH2 [71], and BEAM [27]. Metient and MACH2 used input trees from LAML. **(B)** Compute time per replicate on simulated data (log scale). Additional time to run LAML [31] is shown (*green*) for Metient and MACH2. Some methods were omitted for *n* ≥ 500 owing to time constraints. **(C)** Comparison of tissue-labeled tree inferred by VINE for CP70 from ref. [67] with trees inferred by Cassiopeia-Greedy [59], LAML, and BEAM. Shown for each tree are the cophenetic correlation (*gray*), number of mutations by parsimony (*brown*), tissue homogeneity (*blue*), and number of migrations by parsimony (*violet*) (see **Methods** and **Supplementary Fig. S19**). **(D)** Tissue-labeled MCC tree inferred by VINE for CP4 from ref. [67], comprising 904 distinct barcode sequences across six tissues. **(E–G)** Tissue-migration graphs for CP4 inferred by **(E)** VINE (edges with *>*0.9 variational support, which should not be read as calibrated posterior probabilities), **(F)** Metient (the best scoring of three solutions), and **(G)** MACH2 (the first of three solutions). Shown for each graph are corresponding numbers of migrations (*gray*), co-migrations (*brown*), and seeding sites (*blue*) (see **Methods**, **Supplementary Figs. S20&S21**). Tissue colors match (D). LL, left lung; Liv, liver; M1, mediastinum 1; M2, mediastinum 2; RE, right lung E; RW, right lung W (see [67]).

We then applied VINE in migration mode to the lung-cancer xenograft data from ref. [67], and found that it was able to reconstruct tissue-labeled trees similar to those from BEAM and generally better than those from other methods. Like BEAM, VINE tends to produce cell-lineage trees that group together cells having the same tissue label, resulting in more parsimonious migration histories than Cassiopeia-Greedy [59] or LAML [31], which do not have access to tissue labels (**Fig. 6C**). These improvements are evident in the numbers of both mutations and migrations required to explain the observed data, as well as the tissue homogeneity index, a measure of clustering by tissue label (**Fig. 6C**, **Supplementary Figs. S19&S20**; see **Methods**). Interestingly, in comparison to BEAM, VINE sometimes appears to trade additional mutations for fewer migrations. Nevertheless, most differences between VINE and BEAM trees appear in the detailed branching patterns within single-tissue clades, for which the signal in the data is weak.

The improved efficiency of VINE allowed us to analyze some of the largest CPs from ref. [67], including ones prohibitively large for BEAM or LAML. For example, VINE took only about three minutes to produce a tissue-labeled tree and migration graph for CP 4, which comprises 904 distinct barcode/tissue combinations (**Fig. 6D&E**), whereas the LAML tree inference step alone for this CP (not including migration inference) required several days. For comparison with VINE, we followed ref. [70] in building a cell-lineage tree for this clone using the heuristic Cassiopeia-Greedy method, and then reconstructing migration graphs using MACH2 and Metient (which took 44 min and ∼9 hrs, respectively, with multithreading). We found that VINE obtained considerably simpler tissue-migration graphs than the other methods, requiring ∼40% fewer migrations to explain the observed data (**Fig. 6E–G**, **Supplementary Fig. S21**). The number of inferred migrations from mediastinum 1 (M1) to mediastinum 2 (M2) was particularly diminished. We expect that these simpler graphs primarily result from simultaneous inference of the cell-lineage tree and the migration history, as shown for BEAM [27], although VINE’s tendency to trade migrations for mutations could also contribute to them. In other respects, the tissue-migration graphs were broadly similar. Together, these analyses indicate that VINE’s variational strategy for migration inference maintains many of the strengths of the fully Bayesian BEAM method, but with considerably improved scalability.

## Discussion

Bayesian methods are now widely used in phylogenetic inference, but as datasets steadily grow in size and complexity, the computational cost of MCMC sampling becomes increasingly burdensome. In this article, we have introduced a new computational method, called VINE, that demonstrates for the first time that variational phylogenetic inference can be competitive with state-of-the-art MCMC methods in terms of model fit while offering substantially shorter running times. In addition to several common DNA substitution models, VINE also supports inference of cell-lineage phylogenies based on CRISPR barcode-editing mutation matrices. On this task, VINE fits the data approximately as well as the recently published LAML [31] and BEAM [27] methods, but is significantly faster across a broad range of simulated and real datasets.

We additionally extended VINE to implement the tissue-migration model in BEAM and found, again, that it offered similar performance at much greater speeds. We did observe some reduction in accuracy on this problem in comparison to BEAM, which appears to be related to limits of VINE’s posterior approximation. Nevertheless, the generally strong performance of VINE on this task suggests that it might be worth extending the model to address other inference tasks associated with phylogeny inference. One possibility would be to replace our model of discrete tissue labels with a more general model for latent cell states (e.g., [75]) to infer differentiation maps or transcriptional dynamics. Other possibilities are to adapt VINE to address the analogous problems of phylogeographic reconstruction of ancestral species [76] or ancestral recombination graph (ARG) inference [2].

Development of VI for Bayesian phylogenetic inference has been an active area of recent methodological development in phylogenetics, with numerous innovative new programs appearing over the last few years [35, 38–45]. In our design of VINE, we drew heavily from this emerging literature, focusing in particular on the use of a continuous embedding of taxa, distance-based phylogenetic reconstruction, and stochastic gradient ascent for optimization of the evidence lower bound (ELBO) [44, 45]. This approach has several important advantages over earlier attempts at phylogenetic VI, by avoiding the need to enumerate discrete topological features [35] and aligning naturally with powerful tools for stochastic optimization. It effectively leverages the long history of distance-based methods for phylogenetic reconstruction (e.g., [49, 62]), but marries them with classical likelihood-based strategies within the framework of variational inference. In our experiments, however, we found that previously published phylogenetic VI methods are not yet viable alternatives to MCMC for applied phylogenetics, and typically require orders of magnitude longer run times on datasets of even modest size. By introducing several new algorithmic innovations and optimizing our code, we were able to improve speeds by factors of hundreds to thousands, making it practical for the first time to use VI in large-scale phylogenetics.

One key difference of VINE from similar VI programs such as GeoPhy [44] and Dodonaphy [45] is that, by default, it makes use of a higher-dimensional embedding space, typically with *d* ≥ 5 rather than *d* = 2 or *d* = 3. To our surprise, we found that stochastic optimization was more, rather than less, effective at these higher dimensions, despite the larger number of free parameters. In this way, our approach is less like conventional phylogenetics and more like modern strategies for training deep neural networks, where investigators typically rely on the remarkable effectiveness of SGA to optimize models that are intentionally overparameterized. Notably, the use of a hyperbolic embedding geometry offered no advantage in our hands, although—as suggested by a number of recent papers [44, 45, 48, 65]—it might be possible, with further work, to derive some benefits from non-Euclidean geometries in the context of VINE.

Another key innovation in VINE is its strategy for differentiation through the neighbor-joining or UP-GMA algorithms. Because gradients are approximate anyway in the setting of SGA, we chose not to work with formally differentiable relaxations of these algorithms (see [45]) and instead derived our own recursive procedure for propagating derivatives through the standard algorithms conditional on a choice of nearest neighbors (see **Methods**). Notably, once the neighbors are fixed, both algorithms can be shown to perform linear transformations on a vector of pairwise distances. This strategy allowed us to keep the procedures for backpropagation lightweight and simple, and avoid the heavy computational machinery of automatic differentiation. Our approach effectively factors out the tree topology from gradient calculations, but as we show empirically, the fundamental stochasticity of the optimization algorithm is nonetheless sufficient to ensure that the space of topologies is explored.

In our benchmarking experiments for DNA alignments, we compared the running times of VINE with those of two of the most widely used packages for MCMC-based Bayesian phylogenetics, BEAST 2 and MrBayes. We found that VINE was able to obtain similar model fits with substantially faster running times than these methods, even when the BEAGLE package for CPU acceleration was enabled. There are many challenges in ensuring that such a comparison is fair, however, including the perennial issue of how to assess convergence of the Markov chain (e.g., [77]), as well as more technical concerns such as whether or not to run multiple coupled chains, how to tune proposal distributions, and how to manage parallelization. We sought to place the MCMC and VI methods on an even field by using rigorous convergence criteria and standard models with default proposals, and running all programs on a single CPU core, but these criteria are inevitably somewhat subjective. Still, even if alternative benchmarking strategies were to diminish the speed advantages of VINE, the VI paradigm has the advantage of requiring less tuning of sampling strategies and monitoring of convergence. In addition, once the variational model has been fitted to the data, any number of independent samples can be drawn from the approximate posterior at low computational cost.

At the same time, we observed a persistent tendency in VINE’s variational approximation to underestimate the variance of the posterior distribution—a known problem in VI when the approximating distribution does not have sufficient flexibility to accommodate the structure of the true posterior [34, 78]. We addressed this problem with a second stage that approximates the posterior locally around VINE’s point estimate. This approach yields credible intervals with near-nominal coverage at no additional computational cost, but because it considers only local rearrangements of the tree, it under-represents topological uncertainty, as reflected in elevated split-KL discrepancies (**Supplementary Figs. S9C&D and S13C**). It also has costs in sample-averaged measures of model fit and tree accuracy (**Supplementary Fig. S7B–D**). The approximation is conditional on VINE’s point estimate and on fixed values of the substitution-model parameters. By contrast, current MCMC-based methods appear to explore the full posterior distribution effectively, at least at the scales we considered. At present, they should be preferred to VI in applications that require a complete representation of posterior uncertainty. More work will be needed to improve the quality of VINE’s approximate posterior, for example, through the use of mixture models (e.g., [43]) or by extending the second stage of the re-estimation procedure to broader topological rearrangements.

While this article was under review, a publication appeared on a promising new method for Bayesian phylogenetics, called Delphy [79], that appears to achieve similar improvements in speed to VINE. Instead of using variational inference, Delphy accelerates MCMC sampling by explicitly annotating candidate trees with mutations, thereby avoiding the need for Felsenstein’s pruning algorithm while maintaining a faithful representation of the posterior. This approach is most effective, however, in a particular evolutionary regime, where sequence diversity is low and sampling of taxa is dense, as with pathogen outbreaks (such as our SARS-CoV-2 example). The method also currently assumes dated sequences, a strict molecular clock, an HKY substitution model, and a coalescent prior. By contrast, VINE has a more general design, with support for greater sequence diversity, various substitution models, relaxed clocks, and undated sequences, as well as extensions to CRISPR-based lineage tracing and joint inference of cell migration. Nevertheless, it will be interesting to compare these methods directly, evaluate their relative strengths and weaknesses, and perhaps experiment with hybrid strategies that combine ideas from both methods.

Our benchmarking experiments compared VINE with available methods under a variety of conditions, including several different DNA substitution models, CRISPR as well as DNA mutation matrices, widely varying numbers of taxa and sequence lengths, both real and simulated data, and different levels of evolutionary divergence. Still, many additional features are known to influence the accuracy of phylogenetic reconstructions, such as the prevalence of indels and missing characters, the presence of recombination, and uneven sampling of taxa. It will be important in future work to subject VINE to careful scrutiny under a variety of such conditions to ensure that its performance truly is competitive with the best available methods. A related issue is that, in this work, we have not extensively compared the behavior of VINE with MCMC-based methods under various choices of phylogenetic priors or strategies for inference of substitution rate parameters or other “nuisance” parameters. More work is needed to understand under what conditions differences in priors and inference strategies can significantly alter reconstructed trees.

In our comparisons of DNA alignments, we chose to focus on Bayesian methods to the exclusion of methods based on maximum likelihood (ML). It bears mentioning, however, that programs such as RAxML [29, 80] and IQ-TREE [30, 81] have recently made it possible to perform ML phylogenetic inference at scale with astonishing speeds. These programs remain substantially faster than both VINE and MCMC-based methods and for the foreseeable future will remain better choices for trees with many thousands of taxa. Still, these ML methods generally report a single tree with branch lengths—a point estimate of the phylogeny—rather than attempting to characterize the full posterior distribution, and in many applications it will be worthwhile to expend additional CPU cycles to approximately characterize the uncertainty of the phylogeny.

The full scaling potential of phylogenetic VI based on continuous embeddings of taxa remains unclear. Our current implementation supports multithreading of likelihood calculations, which considerably accelerates model fitting, and GPU acceleration (perhaps via the latest version of BEAGLE [82]) could potentially also be added. It should also be straightforward to reduce memory usage for large data sets, a feature we have not optimized. At the same time, the use of an explicit distance matrix for taxa and distance-based phylogeny reconstruction on each iteration of the algorithm imposes a fundamental lower bound of *O*(*n*^2^) on VINE’s computational complexity. Nevertheless, we speculate that, with aggressive optimizations to the current design (see, e.g., [83]), VINE could perhaps be scaled up one to two orders of magnitude more, allowing practical application to alignments with *>*10,000 taxa. To scale the method further would likely require more heuristic approximations, such as divide-and-conquer approaches. In any case, the work presented here substantially increases the range of datasets to which approximate Bayesian methods can be practically applied.

## Methods

### The core variational inference problem

The core problem is to approximate the posterior distribution of phylogenetic trees given a set of observed genotypes using an approximate variational distribution *q*(*τ,* **b**), where *τ* is a rooted binary tree with *n* tips and **b** ∈ R^2^*^n^*^−2^ is a corresponding vector of nonnegative branch lengths. We denote the data by an *n* × *L* matrix **X**, which may be either a standard multiple alignment of DNA sequences or a mutation matrix representing CRISPR-edited barcodes (with *L* observed sites in either case).

In the standard manner for variational inference (reviewed in [34]), we fit the model by minimizing the Kullback-Leibler (KL) divergence between *q*(*τ,* **b**) and the true posterior distribution, *p*(*τ,* **b** | **X**). The KL divergence can be expressed as,

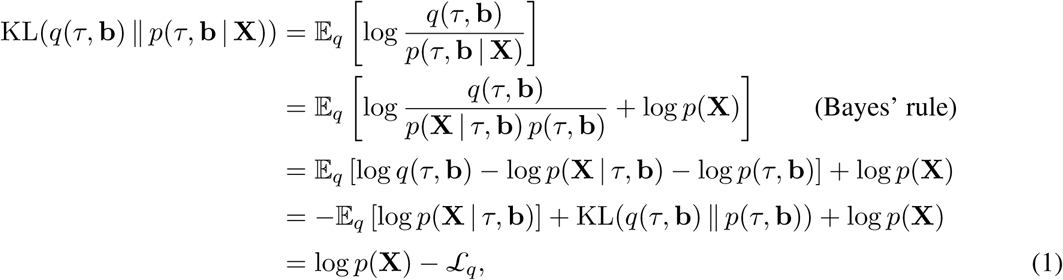

where the expectations are evaluated with respect to *q*(*τ,* **b**) and the evidence lower bound (ELBO) *L_q_*is defined as,

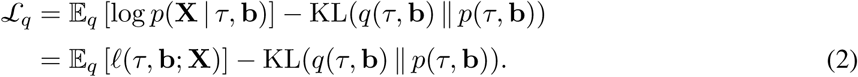

Here we introduce the notation *ℓ*(*τ,* **b**; **X**) = log *p*(**X** | *τ,* **b**) for the standard phylogenetic log likelihood, that is, the log probability of the genotype data given the tree. Notice that, because the KL divergence must be nonnegative, the ELBO *L_q_* is a strict lower bound on the marginal log likelihood, log *p*(**X**). The essential idea of variational inference, therefore, is to choose the free parameters of *q* to maximize *L_q_*, forcing the KL divergence to shrink toward zero and making *q*(*τ,* **b**) approximate the true posterior distribution as closely as possible given its functional form.

### Continuous embeddings and differentiable transformations

Our general strategy (**Fig. 1**) is to sample embeddings **x** ∈ R*^nd^* of *n* taxa in *d*-dimensional space from a multivariate normal (MVN) distribution *q* = MVN(***µ***, **Σ**), with ***µ*** ∈ R*^nd^* and **Σ** ∈ R*^nd^*^×^*^nd^*; to compute an induced distance matrix **D** ∈ R*^n^*^×^*^n^* from **x**; to obtain a rooted phylogenetic tree *τ* with branch lengths **b** ∈ R^2^*^n^*^−2^ from **D** using a deterministic, distance-based tree reconstruction algorithm; to compute the phylogenetic log likelihood *ℓ*(*τ,* **b**; **X**) based on that tree and the specified mutation model; and, finally, to estimate the ELBO using an expectation of the phylogenetic log likelihood. We then compute an approximate gradient of the ELBO, conditional on the tree topologies of the sampled points (see below), with respect to the free parameters (***µ***, **Σ**) by backpropagation through all steps of this transformation, and optimize the parameters by SGA. Notably, all *nd* components of ***µ*** are maintained as free parameters in the optimization problem, but various reduced parameterizations can be assumed for **Σ** (see below).

To allow for pathwise differentiation with respect to ***µ*** and **Σ**, we use the standard technique of reparameterizing the MVN variational distribution (the “reparameterization trick”). We introduce a standard multivariate normal random variate **z** ∈ R*^nd^*, with **z** ∼ MVN(**0**, **I**), and redefine **x** as,

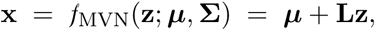

where **L** is a (lower-triangular) Cholesky factor such that **Σ** = **LL**^⊤^. In this way, the expressiveness of the MVN is maintained but the randomness in samples from *q* now derives from a parameter-free density, and all subsequent transformations are deterministic and differentiable.

A second transformation produces a symmetric matrix of pairwise distances **D** ∈ R*^n^*^×*n*^ from **x**:

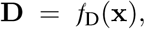

where *f*_D_ is such that *D_ij_* = *d*(**x***_i_,* **x***_j_*), where **x***_i_* ∈ R*^d^* denotes the embedded point for taxon *i* and *d* is a suitable distance function between points in the embedding space (detailed below).

Finally, a third transformation produces a tree with branch lengths, (*τ,* **b**), from **D** using a deterministic distance-based phylogeny reconstruction algorithm:

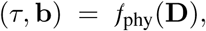

where *τ* is a rooted binary tree on *n* leaves and **b** ∈ R^2^*^n^*^−2^ contains the corresponding branch lengths. In the case of DNA substitution models, we use the neighbor-joining algorithm [49], and in the case of CRISPR barcode editing models, where an ultrametric tree is customary, we use the UPGMA (Unweighted Pair Group Method with Arithmetic Mean) algorithm [62]. In order to allow informative priors to be applied, we impose a rooting on neighbor-joining trees using the midpoint method (root at midpoint of longest span between taxa). In the CRISPR case, we introduce an additional free parameter for a leading branch to the root of the tree, as is typical in this literature.

Taking advantage of the reparameterization trick, let *g* represent the full transformation from **z** to (*τ,* **b**):

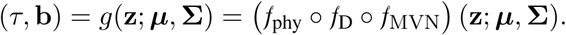

Notice that *g* depends only on ***µ*** and **Σ**, because *f*_phy_ and *f*_D_ are parameter-free.

The ELBO *L_q_*can therefore be re-expressed explicitly in terms of ***µ*** and **Σ** as,

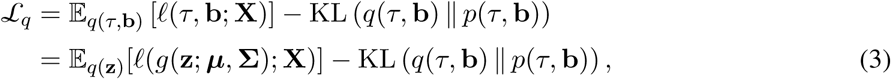

where we take care to specify the distribution used for each expectation. Here, *q*(**z**) denotes the standard MVN for **z** and *q*(*τ,* **b**) denotes the induced distribution over phylogenetic trees.

To proceed further, we must specify the prior distribution, *p*(*τ,* **b**). We consider two cases. In case (1), the default in VINE, we allow the prior to be defined implicitly by assuming that the MVN random variate **x** ∼ MVN(**0**, **I**), as is often done with variational autoencoders. In case (2), discussed below, we allow for a general prior distribution over phylogenetic trees. Details on phylogenetic priors supported by VINE are provided in the **Supplementary Materials**.

In case (1), the variational distribution is defined entirely with respect to **x**; the tree (*τ,* **b**) is simply a deterministic function of this random variable that enables us to evaluate the phylogenetic likelihood. Therefore, we can write,

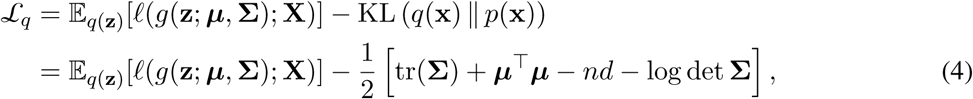

where the last term represents the KLD for *q*(**x**) = MVN(***µ***, **Σ**) with respect to *p*(**x**) = MVN(**0**, **I**).

The first term of equation 4 is simply the expectation of the standard phylogenetic log likelihood under the variational distribution. Because of the complex nonlinear nature of the transformation *g*, there is no straightforward way to calculate this quantity exactly, but it can easily be estimated by Monte Carlo sampling (an alternative estimation method based on a Taylor approximation is described below). As a result, the approximate ELBO becomes,

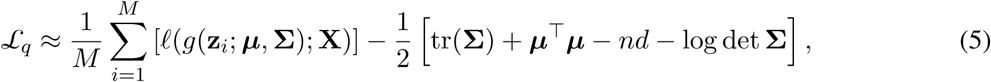

where **z***_i_*is the *i*th sample drawn from *q* and *M* is the number of samples.

In case (2), allowing for a general phylogenetic prior, *p*(*τ,* **b**), the KLD is no longer available in closed form, so we re-express the ELBO in a form more typical for standard VI. Making the transformation *g* explicit, we have,

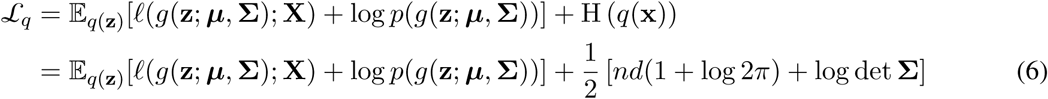

where H(*q*(**x**)) denotes the entropy of *q*(**x**), which is also available in closed form. The expectation in equation 6 now represents the expected complete-data log likelihood, including both the prior *p*(*τ,* **b**) and the conditional likelihood, *p*(**X**; *τ,* **b**). These terms can be evaluated together by Monte Carlo sampling, in an analogous manner to case 1,

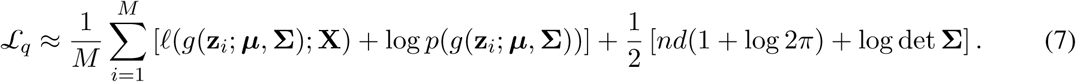

Importantly, whereas in case (1), equation 4 represents a valid ELBO for a generative model defined over the embedding **x**, in which the tree (*τ,* **b**) is a deterministic function of **x**, in case (2), equation 6 does not represent the true tree-space ELBO, because it substitutes the entropy of the embedding distribution, H(*q*(**x**)), for the entropy of the induced tree distribution, H(*q*(*τ,* **b**)). Because *g* is many-to-one and maps a continuous embedding to a mixed discrete-continuous space of trees, these quantities are not equal in general, and the usual change-of-variables argument does not apply. Equation 6 should therefore be regarded as a surrogate objective that replaces an intractable entropy with a tractable one (see [44] for a more rigorous approach). In this work, we assume case (1) for all main results and make use of case (2) only for **Supplementary Fig. S23**. Notably, during optimization by SGA, VINE uses a tempered version of these objective functions in which the log-determinant term in equations 4 and 6 is down-weighted (see **Stochastic gradient ascent**).

Notice in both case (1) and case (2), that for each sample **z***_i_* and induced tree (*τ_i_,* **b***_i_*) = *g*(**z***_i_*), the log likelihood can be evaluated efficiently in the standard manner, using Felsenstein’s pruning algorithm and either an appropriate DNA substitution model or a mutation model for CRISPR barcodes. Thus, the calculation of the objective function reduces to elementary operations and well-known algorithms for phylogenetics (neighbor joining/UPGMA and the pruning algorithm).

### Chain rule for gradients

The gradient of the Monte-Carlo approximated ELBO (equation 5) with respect to the free parameters of ***µ*** and **Σ** can be approximated in terms of gradients of the phylogenetic log likelihoods for the sampled points. In case (1) above (equation 5),

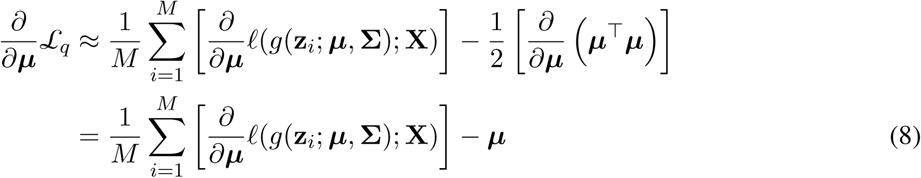

and

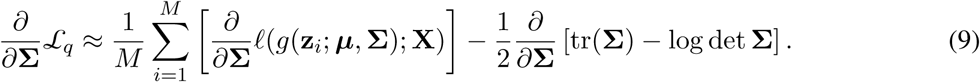

Similarly, in case (2) (equation 7),

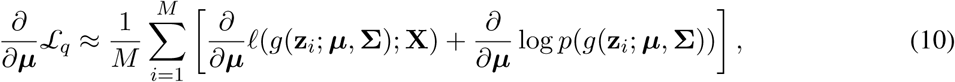

because the entropy of *q*(**x**) does not depend on ***µ***, and,

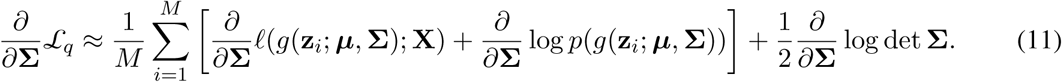

The last terms in equations 9 and 11 can be computed in closed-form in a manner that depends on the choice of parameterization for **Σ**, as shown later. The key challenge is therefore to compute gradients for the phylogenetic log likelihood, *ℓ*(*g*(**z***_i_*; ***µ***, **Σ**); **X**), and optionally, the corresponding log prior term. We will focus on the log likelihood here; the prior follows by analogy.

We start by observing that, in principle, we can propagate derivatives forward along the sequence of component functions *f*_MVN_, *f***_D_**, and *f*_phy_by computing the corresponding Jacobian matrices. In particular, for a vector of free parameters ***ϕ*** ∈ {***µ***, **Σ**} of dimension *k*, we can write,

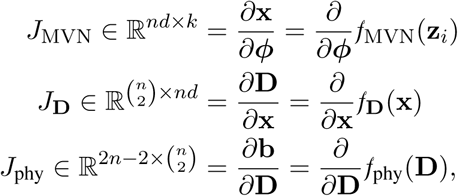

where in the last step we consider the branch lengths **b** only and ignore the tree topology *τ*, as explained in the **Supplementary Material**. To complete the process, we must also consider the gradient of the phylogenetic log likelihood with respect to the 2*n* − 2 branch lengths:

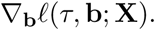

This is a familiar quantity in phylogenetic analysis, which can be efficiently calculated using an inside-outside algorithm on the tree (see **Supplementary Material**).

Assuming for the moment that these objects can all be obtained, the gradient of interest can be computed by a straightforward application of the chain rule:

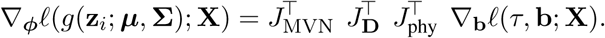

The product of three Jacobian matrices thus converts the standard (2*n* − 2)-dimensional branch-length gradient to a *k*-dimensional parameter gradient, as required. In this way, ∇**_b_***ℓ*(*τ,* **b**; **X**) can be propagated backwards from the branch lengths through the distance matrix and embedded points, to the MVN parameters ***µ*** and **Σ**.

In practice, we avoid instantiating the full Jacobian matrices and instead implicitly propagate gradients through them using an efficient reverse-mode algorithm. Moreover, the first two Jacobians reflect elementary differentiable functions and can be accommodated analytically. The main difficulty lies with the third Jacobian, *J*_phy_, which reflects the entire transformation of a distance matrix **D** to a phylogenetic tree with branch lengths. An algorithm for efficiently computing this Jacobian implicitly and additional details are provided in the **Supplementary Material**.

The per-sample derivatives described above are computed conditional on the topology of each sampled tree, so they do not account for the discontinuities in *g* that occur when a change in the embedding alters the sequence of neighbors selected by NJ or UPGMA. Because the boundaries between topologies shift as ***µ*** and **Σ** change, the gradient of the expected log likelihood generally includes an additional boundary term that these derivatives do not capture [84]. The gradients used by VINE are therefore biased, topology-conditional surrogates for the true gradients of the ELBO.

### Implementations of Neighbor-Joining and UPGMA

VINE uses custom implementations in C of the NJ and UPGMA algorithms that take a distance matrix as input and return a tree with branch lengths. In the case of NJ, the sequence of merged neighbors and associated metadata are recorded for later use in backpropagation (see **Supplementary Material**). Because these tree-building routines are rate-limiting for variational inference, we initially adapted the standard algorithms to use a min-heap for efficient identification of the pair of nodes to join on each step (e.g., the minimum entry of **Q** in NJ), reducing the asymptotic running time from *O*(*n*^3^) to *O*(*n*^2^ log *n*) (see, e.g., [85]). In the end, however, we found the overhead of the heap to be prohibitive and instead adopted a strategy involving on-the-fly calculation of distances, which has an *O*(*n*^3^) bound but is considerably faster in practice.

### Taylor approximation of the ELBO

Estimating the ELBO by Monte Carlo sampling (equations 5 & 7) is effective for optimization but computationally expensive. We found that the efficiency of the algorithm can be substantially improved by making use of a Taylor approximation for the ELBO. In particular, we approximate the expectation of the log likelihood (see equation 4) using a second-order Taylor approximation around the mean (corresponding to **z** = **0**),

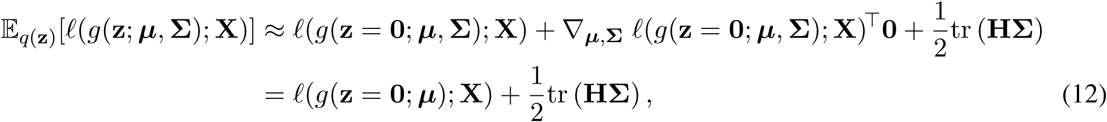

where **H** is the Hessian matrix for the entire transformation, *ℓ*(*g*(**z** = **0**; ***µ***, **Σ**); **X**), as evaluated at **z** = **0**. (Here we focus on case (1) for the prior, **x** ∼ MVN(**0**, **I**); case (2) follows by analogy; see equation 6.)

If we ignore the dependency of **H** on ***µ***, which we expect to be weak, then the gradient of the approximate ELBO with respect to ***µ*** depends only on the first term of equation 12 and the gradient with respect to the covariance **Σ** depends only on the second term. Thus, this formulation allows approximate decomposition of the ELBO into mean- and variance-related components. In practice, we find that mean-related component is strongly dominant when the model is fitted to data.

On its face, the second term in equation 12 still poses a problem, because the *nd* × *nd* Hessian matrix is impractical to instantiate explicitly. It is possible to estimate this term efficiently by way of matrix-vector products, without realizing the Hessian, using Hutchinson’s method [86]. In practice, however, we find that we obtain a better approximation in comparable time by periodically estimating the LHS of equation 12 by our standard Monte Carlo method and then estimating the second term on the RHS by the difference,

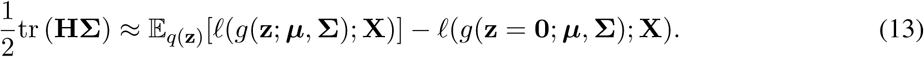

Because this quantity is not highly sensitive to ***µ***, it need not be updated on every iteration of SGA. We delay computing it during a “warmup” period (50 iterations), and then update it only every 30 iterations, using an exponential moving average for both the trace quantity and its gradient. This strategy requires less than one additional evaluation per iteration of the full transformation *g*, decreasing the overall cost of SGA by nearly a factor of *M* in comparison to full Monte Carlo sampling. Overall, we find that this strategy closely approximates the behavior of full Monte Carlo sampling with substantial savings in compute time (see **Supplementary Fig. S22**).

### Parameterizations of the covariance matrix Σ

VINE supports four parameterizations of the MVN covariance matrix **Σ**, which are labeled CONST, DIAG, DIST, and LOWR in order of increasing complexity (accessible via the `--covar` option). The CONST parameterization simply assumes **Σ** = *λ***I** and ensures nonnegativity of *λ* by defining it as *λ* = *e^η^*, with *η* as a free parameter. The DIAG parameterization allows for a general diagonal covariance matrix, with free variance along each of the *nd* dimensions but no covariance between them: **Σ** = diag{*λ*_1_*, . . ., λ_nd_*}, with each *λ_i_* = *e^η^^i^* for *nd* free parameters.

The DIST and LOWR parameterizations attempt to capture more of the covariance structure across taxa while keeping the parameterization as sparse as possible. In these cases, the same covariance structure is assumed for each of the *d* dimensions in the embedding space, with no covariance across these dimensions. The full covariance matrix can therefore be described as **Σ** = **I***_d_* ⊗ **Σ**_0_, where **I***_d_* is a *d* × *d* identity matrix, ⊗ is the Kronecker product, and **Σ**_0_ is the *n* × *n* covariance matrix shared for each embedding dimension. In these cases, VINE gains further efficiency by performing all MVN-related calculations on *d* independent *n*-dimensional MVNs with a shared covariance structure, rather than on one *nd*-dimensional MVN.

The DIST parameterization further assumes **Σ**_0_ = *λ***S** and *λ* = *e^η^*, resulting in a single free variance parameter *η* for all taxa. In this case, **S** is kept fixed at a double-centered version of the initial distance matrix **D**_0_,

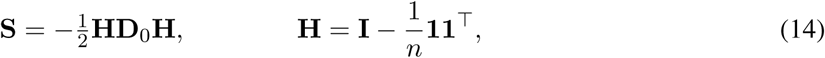

where **1** indicates an *n*-dimensional vector of all 1s and **11**^⊤^ is therefore an *n* × *n* matrix of all 1s. This matrix **S** captures the expected covariance structure of a random variable evolving by Brownian motion along the branches of an unrooted tree, and therefore, the DIST parameterization is a simple way to consider the distances between taxa without restricting the tree topology.

The most flexible parameterization, LOWR, allows for a general low-rank representation of **Σ**_0_, with **Σ**_0_ = **RR**^⊤^ for a general matrix **R** of dimension *n*×*w* (with *nw* free parameters; by default, *w* = 3). In this case, the full covariance of the embedding is Σ = (*RR*^⊤^) ⊗ *I_d_*, which has rank at most *wd*. Consequently, VINE can reparameterize sampling from MVN(***µ***, **Σ**) using only *wd* latent dimensions, instead of the full *nd* dimensions required by an unrestricted covariance matrix. When *w < n*, **Σ**_0_ is singular, so VINE replaces log det **Σ**_0_ in the KLD and entropy terms with the pseudo-determinant log det(**R**^⊤^**R**) (the sum of the logarithms of the nonzero eigenvalues of **RR**^⊤^).

All of these parameterizations permit closed-form expressions for the gradient of the KLD (equation 9) or entropy (equation 11) and for the Jacobian *J*_MVN_ (see **Supplementary Material**).

Note that most results reported in this paper reflect the default CONST parameterization. In addition, most results use the Euclidean geometry, the Taylor approximation, the implicit Gaussian prior, and the default number of dimensions *d*, which is a function of the number of taxa *n*: *d* = ⌊3.25 + 0.92 ln(*n*)⌋ (exceptions are shown in **Fig. 3** and **Supplementary Figs. S6,S22,** and **S23**). These default parameter choices are recommended for general-purpose use. In cases where it is critical to characterize posterior uncertainty, however, we recommend the two-stage procedure described below (`--posterior`), which leaves these choices unchanged in the first stage and approximates the posterior locally in a second stage (**Fig. 3**).

### Euclidean and hyperbolic geometries

By default, VINE embeds taxa in a *d*-dimensional Euclidean space, where *d* can be selected by the user via the `--dimensionality` option. (A general-purpose default is determined as a linear function of log *n*, where *n* is the number of taxa.) Let **x***_i_* ∈ R*^d^* denote the embedded point for taxon *i*. The Euclidean pairwise distance is given by,

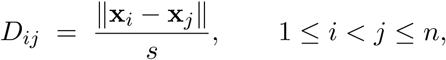

where *s >* 1 is a global scale factor that allows the embedding space to be expanded relative to the distance matrix **D**.

The initial mean ***µ*** of the MVN variational distribution is estimated by classical multidimensional scaling (MDS) applied to the starting distance matrix **D**_0_. Let **D**^2^_0_ be the (symmetric) matrix of squared distances, double-centered to obtain a Gram matrix **G**. An eigendecomposition **G** = **VΛV**^⊤^ is computed, and the initial embedding is given by the top *d* scaled principal components, 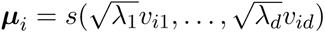 where *s* is the scale factor.

In the hyperbolic setting, VINE follows the general approach outlined by Macaulay et al. [65] and embeds points on the upper sheet of the (*d*+1)-dimensional hyperboloid model with negative curvature −*κ* (set by `--negcurvature`, default −*κ* = 1). Each taxon is represented by a spatial coordinate **x***_i_* ∈ R*^d^*, which is lifted to a point on the hyperboloid via,

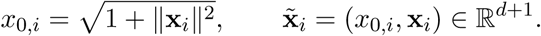

The hyperbolic distance between taxa *i* and *j* is,

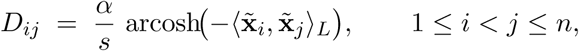

where 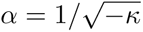 is the curvature radius and ⟨x_i_, x_j_⟩*_L_* is the Lorentz inner product,

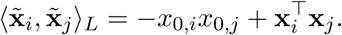

For hyperbolic embeddings, the MVN mean is initialized using a spectral variant of the *hydra* algorithm [87]. Given an initial distance matrix **D**^(0)^, we form a symmetric matrix **A** such that,

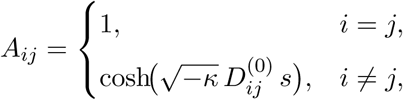

diagonalize **A** = **VΛV**^⊤^, and set the initial spatial coordinates from the leading *d* eigenmodes, followed by the same global scaling by *s*.

The scale factor *s* is chosen so that the median pairwise distance between points is 25 in the Euclidean case and 4 in the hyperbolic case. The Jacobians of these mappings from embedded points to pairwise distances (*J***_D_**) are provided in the **Supplementary Material**.

### Stochastic gradient ascent

For stochastic gradient ascent (SGA), VINE uses a customized implementation in C of the Adam algorithm [88]. In the Monte-Carlo case, the algorithm draws a minibatch of *M* sets of embedded points on each iteration, **x**_1_*, . . .,* **x***_M_* (transformed from **z**_1_*, . . .,* **z***_M_*), and uses them to obtain estimates of the ELBO and its gradient. For each minibatch, it precomputes KL(*q*(**x**) *p*(**x**)) and its gradient, which do not depend on the sampled points. It then updates all parameters based on the gradient in the standard manner. Command-line options allow the user to control the minibatch size (`--batchsize`; default 10) and the learning rate *α* (`--learnrate`; default 0.05), as well as convergence criteria for the algorithm (e.g., minimum number of iterations, `--miniter`, default 200, and number of iterations over which to average when assessing convergence, `--niterconv`, default 50). In the case of the Taylor approximation, the procedure is similar but no minibatch sampling is required; the ELBO is estimated directly from the value of the log likelihood at the current mean ***µ*** and the estimate of the trace term (equation 12). In both cases, the Adam parameters *β*_1_ and *β*_2_ are both fixed at 0.9. A simple scheduler is employed to manage subsampling of sites in the alignment in early iterations of the algorithm, adaptive clipping of unusually large gradients, and gradual increases of the learning rate to its target value. The algorithm also allows for a separate set of “nuisance parameters” (including parameters for substitution rates and tree priors), which are also optimized by SGA but are not encoded in the continuous embedding.

During SGA, VINE multiplies the closed-form terms in equations 4 and 6 by *β* = 1*/s*^2^, where *s* is the scale factor of the embedding, and therefore optimizes

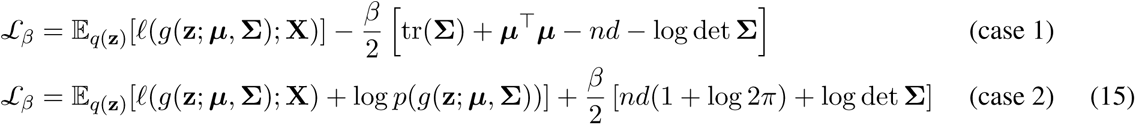

Because ***µ*** and **Σ** are expressed in the units of the embedding, in which the prior is MVN(**0***, s*^2^**I**), the factor of 1*/s*^2^ for tr(**Σ**) and ***µ***^⊤^***µ*** simply reflects this change of scale. The substantive effect of *β* is to down-weight the log-determinant term, which corresponds to the entropy of *q*(**x**). Down-weighting the entropy in this way is equivalent to targeting a tempered posterior distribution [89, 90]. We found that it yielded more accurate point estimates than the untempered ELBO, which led to overdispersion for larger trees. When a calibrated measure of uncertainty is needed, VINE instead approximates the posterior locally in a second stage (see **Two-stage estimation of posterior uncertainty**, below).

The SGA algorithm has no fixed iteration limit and terminates when the objective function reaches a plateau. Once the designated minimum number of iterations (`--miniter`) has been exceeded, convergence is assessed after every block of `--niterconv` iterations, and the algorithm terminates when 1.001 · *S_t_* ≤ 0.999 · *S_t_*_−1_, where *S_t_*is the average ELBO for block *t*. The parameter set corresponding to the single iteration with the highest ELBO encountered is retained.

Because *g* is discontinuous at changes in tree topology, an optimizer with a fixed step size could, in principle, oscillate across a topology boundary. Three features of VINE limit this behavior. First, Adam normalizes each update by a running estimate of the gradient magnitude, so per-coordinate steps are bounded by roughly the learning rate *α* (0.05 by default), which is small relative to the scale of the embedding. Any oscillation is therefore confined to a small neighborhood. Second, sustained oscillation would cause the block-averaged ELBO to plateau, which triggers termination. Third, VINE retains the parameters with the highest ELBO, so oscillation cannot degrade the returned solution. For additional insurance, VINE optionally applies a diminishing step size (`--lrdecay`) to dampen late oscillation, although this option was not used for the results reported here.

The SGA algorithm was carefully tuned to optimize the speed and accuracy of convergence over datasets of various sizes. During this process, we found that it was quite sensitive to the scale of the embedding space, sometimes struggling to converge when distances between points grew close to zero. Therefore, we introduced the scaling factor *s* that allows for a difference in scale between the embedding space and the distances used for tree reconstruction (as detailed above). In particular, VINE rescales the MVN distribution for embedded points so that median distances between points are 25 and 4, respectively, for the Euclidean and hyperbolic geometries.

### Two-stage estimation of posterior uncertainty

When the `--posterior` option is selected, VINE approximates the posterior in a second stage, after SGA has converged. This approximation is local to the tree implied by ***µ*** and nuisance parameters are held fixed. The branch lengths of the starting tree are first optimized by Newton’s method, using the analytical gradient and a diagonal, finite-difference Hessian. Each branch length is then approximated by an independent normal distribution truncated at zero (a Laplace approximation) with mean and variance determined by the local curvature. Topological uncertainty is captured by performing nearest-neighbor interchanges (NNIs) at each internal branch, weighted alternative trees by their likelihoods (as in [91]). Each sampled tree combines independently drawn NNIs and branch lengths. For ultrametric trees (including all CRISPR analyses), the same scheme is applied to node heights, with an additional global scale factor. Poorly supported branches are collapsed into polytomies and resolved at random. This approximation captures uncertainty only near the point estimate, but it is fast and appears to be well calibrated. Indeed, overall running time is slightly reduced (**Supplementary Fig. S7A**), owing to savings in neighbor-joining reconstructions of sampled trees. Two-stage estimation is currently available only for case (1) (the default implicit prior) and the Euclidean geometry.

### Mutation models

VINE includes implementations of the Jukes-Cantor (JC) [54], HKY [52], and general time reversible (GTR) [57] DNA substitution models, as well as the CRISPR-barcode mutation model developed by Seidel et al. [24] and extended by Chu et al. [31]. It also supports the discrete gamma model for rate variation among sites [58] with DNA substitution models. For the DNA models, the substitution rate matrix **Q** is normalized in the standard manner, such that *_i_*_̸=_*_j_ π_i_q_ij_* = 1, where *π_i_* is the equilibrium frequency of nucleotide *i* under the model, so that branch lengths can be interpreted as having units of expected substitutions per site. For the HKY and GTR models, equilibrium frequencies for the four nucleotides are simply estimated using the relative frequencies in the alignment, whereas under the JC model they are assumed to be uniform. As a result, the JC model has no free substitution rate parameters, the HKY model has a single free parameter—the transition/transversion rate ratio *κ*—and the GTR model has five free parameters (after accounting for rescaling). These are considered as “nuisance” parameters that are optimized in SGA but not considered for full variational Bayes characterization. The CRISPR model has a single free parameter, corresponding to the silencing rate for barcodes, which is also treated as a nuisance parameter. VINE supports two parameterizations for the barcode-editing rate matrix: a single rate matrix shared by all sites (as in TiDeTree [24]; `--crispr-modtype GLOBAL`) or a separate rate matrix per site (as in LAML [31]; `--crispr-modtype SITEWISE`). It also allows for either a uniform distribution for all mutation-rate priors (`--crispr-mutprior UNIF`) or a prior that reflects the relative frequencies of mutations in the input matrix (`--crispr-mutprior EMPIRICAL`). The CRISPR mutation matrix is subjected to the same scaling constraint as the DNA substitution matrix, so the estimated branch lengths can be interpreted in expected mutations per site. For comparison, LAML trees must be scaled by the separately estimated mutation rate.

### Simulations

For our simulations of DNA alignments, we used a script based on DendroPy [92] to simulate trees under a birth–death process (birth rate 1.0, death rate 0.5), with three-fold oversampling of tips followed by pruning and rescaling, to produce tree heights of 1 substitution per site and minimum branch lengths of 0.02. Branch-specific rates were drawn independently under an uncorrelated log-normal relaxed clock with mean of 1 and standard deviation of 0.6. Nucleotide alignments were then generated for each tree using `base_evolve` from PHAST under an HKY model with *κ* = 4 and equilibrium frequencies of *π*_A_ = *π*_T_ = 0.3 and *π*_C_ = *π*_G_ = 0.2. As noted in the text, we generated alignments of 300 bp and 10,000 bp, for various numbers of taxa *n* between 10 and 1000. For held-out data, we ran `base_evolve` a second time on the same simulated trees used to generate the training data.

For CRISPR lineage-tracing simulations, we used Cassiopeia’s [59] BirthDeathFitnessSimulator with birth rate 0.075 and death rate 0.005, drawing branch waiting times from the absolute value of a normal distribution (*µ* =1/rate, *σ* = *µ/*5) to obtain approximately balanced ultrametric trees, which were post-scaled to a fixed height of 54 days to match the dataset of ref. [67]. We then applied Cas9LineageTracing-DataSimulator using a cassette of 3 CRISPR target sites repeated 10 times (30 total sites), a mutation rate of 0.01, a heritable silencing rate of 1 × 10^−4^, and no stochastic silencing. Edit outcomes were restricted to a predefined library of 100 possible states with fixed probabilities, ensuring reproducible mutation signatures across simulations. The simulated trees have branch lengths in units of time, whereas VINE and LAML estimate branch lengths in units of expected edits per site. For all comparisons with the true trees that depend on branch lengths (branch-score distances and 95% CI inclusion of pairwise distances), we therefore converted the true trees to expected edits per site by multiplying all branch lengths by the simulated edit rate (0.01). BEAM’s samples, which are in units of time because its origin time was fixed at the true experiment duration (54 days), were converted by the same factor when compared with VINE’s samples (distance-KL discrepancy).

### Convergence criteria for MCMC

For convergence monitoring of MrBayes and BEAST 2, we applied a uniform procedure across all analyses for both simulated and real data. First, for each combination of taxa number, sequence length, substitution model, and program (BEAST 2 or MrBayes), we ran three pilot MCMC replicates with two chains per replicate and intentionally long chain lengths, recording all samples (no thinning). For each replicate, we then retrospectively computed the rank-normalized split *R*^^^ [93] and the effective sample size (ESS) for each of several scalar summaries in each chain in intervals of at least 100,000 iterations. For BEAST 2, the selected statistics were the log posterior, log likelihood, tree height, and tree length, and for MrBayes they were the log likelihood and tree length. For each replicate, we then identified the earliest MCMC iteration calculation at which all parameters met ESS ≥ 400 and *R*^^^ ≤ 1.01. (For the large alignments—with 10,000 columns—we had to relax the *R*^^^ criterion in a few cases because not all pilot replicates converged.) ESS calculations were based on the implementation in Tracer [94], reimplemented in our own scripts. Finally, we averaged these convergence points across the three replicates to determine the target chain length. These target chain lengths were then applied uniformly across all ten replicates for each simulated dataset size, model, and program. Notably, the chain lengths required for our convergence and sampling criteria were generally substantially shorter for MrBayes than for BEAST 2, and this was the dominant factor in the differences in running times we observed. Note also that we experimented with a more stringent requirement of ESS ≥ 625 but found it to require excessively long chains in some cases, so we relaxed the threshold to 400, as recommended in ref. [77].

### Evaluating posterior distributions

We evaluated posterior distributions from VINE and BEAST 2 using two complementary measures: 95% credible-interval (CI) inclusion and average marginal Kullback–Leibler (KL) discrepancies of the posterior approximation with respect to an MCMC-based reference distribution (**Fig. 3**). Both measures were computed from the posterior samples of trees output by each program after running on simulated data. These calculations are supported by the `evalTrees` and `compareTrees` utilities distributed with VINE.

To assess 95% CI inclusion, we extracted the empirical distribution of values induced by the posterior samples for each pairwise distance between taxa, then excluded the top and bottom 2.5% of values to obtain an empirical 95% CI. We then compared this range with the true value used in simulation. The reported values are the fractions of true pairwise distances that fall within the corresponding 95% CI.

To evaluate the discrepancy from a reference distribution, we used two methods, one based on induced pairwise distances and one based on the relative frequencies of “splits,” or bipartitions of taxa, present in the sampled topologies. Because both measures average KL divergences between marginal distributions (of individual distances or splits), we refer to them as the average marginal distance-KL and split-KL discrepancies, respectively. For the distance-based discrepancy, we first approximated the posterior distribution of pairwise distances between each pair of taxa, (*i, j*), as a Gaussian distribution with mean *µ_i,j_* and variance *σ*^2^, where *µ_i,j_* was estimated as the sample mean of all patristic distances (along the tree) between *i* and *j* from sampled trees and *σ_i,j_* was estimated by the corresponding sample variance. Given such estimates for a query and a reference distribution, the directional average marginal distance-KL discrepancy was computed as,

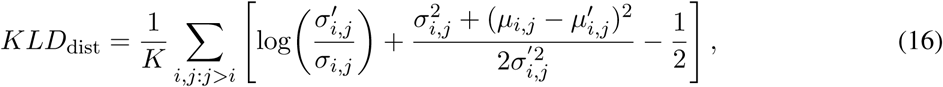

where *µ_i,j_*and *σ*^2^ indicate the mean and variance, respectively, for the query distribution; *µ*^′^ and *σ* indicate the mean and variance for the reference distribution; and *K* is the total number of taxon pairs after excluding any cases with non-finite estimates or zero variance.

For the split-based discrepancy, we start by defining smoothed inclusion probabilities, *p_s_*, for each observed split *s* and each set of sampled trees:

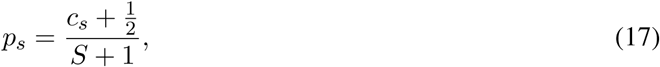

where *S* is the total number of sampled trees, *c_s_* is the number of trees containing split *s*, and a pseudocount of ^1^ is used to avoid estimates of *p_s_* at the boundaries of zero or one. We then compute the average marginal split-KL discrepancy from independent Bernoulli distributions as,

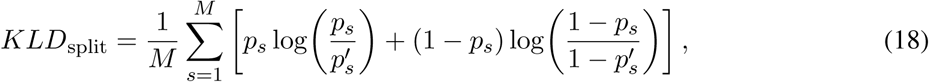

where *p_s_* is the smoothed inclusion probability for the query distribution, *p*^′^ is corresponding value for the reference distribution, and *M* is the total number of splits observed in either distribution. For both measures, natural logarithms were used throughout.

### Models selected for analysis of simulated DNA

We selected models in BEAST 2 (v2.7.7) and MrBayes (3.2.7a) that were as close as possible to the ones in VINE given choices available for each program, generally keeping the prior distributions diffuse so that the log likelihood dominated in inference. In particular, in BEAST 2, we used a Yule prior for the phylogeny with a single free birth-rate parameter (uniform prior) and an uncorrelated log-normal relaxed clock (exponential prior with mean 2 on ucldStdev, the standard deviation in log-space). For the *κ* parameter in the HKY model, we used a log-normal prior with a log mean of 1.386 (corresponding to *κ* = 4, the value used in our simulations) and a standard deviation of 0.1. Similarly in MrBayes, we used a uniform prior over tree topologies and the default Gamma-Dirichlet prior for branch lengths. For the HKY model, we used a Beta(4,1) prior distribution for the rate of transitions relative to the rate of all substitutions. In both programs, we did not allow for variation across sites in rates, we used random starting trees, and we used empirical frequencies for the four nucleotides.

In VINE we used an implicit prior on trees corresponding to a standard multivariate normal distribution for *p*(**x**) (the default, as discussed above). A set of validation experiments with an informative Yule prior and a relaxed local clock suggested that the choice of prior had little effect on the relative performance of the methods (**Supplementary Fig. S23**). The HKY parameter *κ* was optimized as a nuisance parameter in SGA (`--hky85` option). Empirical nucleotide frequencies were used. Analyses that required sampled trees (such as the comparison of Robinson-Foulds distances) were based on 1000 samples from the approximate posterior following convergence of the ELBO (`-s 1000`). In most cases, default parameters were used for the minimum number of iterations before convergence (`--miniter 200`) and the number of iterations over which to average when assessing convergence (`--niteconv 50`), although for some larger trees we used larger values for the minimum number of iterations.

The same choices of models and parameters for BEAST 2, MrBayes, and VINE were used in the benchmarking analysis of real nucleic-acid data from ref. [64], as shown in **Supplementary Figs. S14 & S15**.

### SARS-CoV-2 Analysis

We downloaded raw sequences (`sequences.fasta.xz`) and accompanying metadata (`metadata.tsv.xz`) from https://data.nextstrain.org/ (in `files/ncov/open/100k`), as well as the associated alignment (`aligned. fasta.xz`). All source data were downloaded on February 25, 2026. We used `augur` to exclude sequences less than 29,000 bp in length (`--min-length 29000`) or lacking complete date information (`--exclude-ambiguous-dates-by any`). We then performed stratified subsampling by region and date (`--group-by region year month --sequences-per-group 3 --subsample-seed 1`) to reduce the data set to 1060 aligned sequences. A more stringent stratified sampling step produced the smaller set of 364 sequences (`--group-by region year month --sequences-per-group 1 --subsample-seed 2`). We analyzed the resulting alignments using the same settings as for simulated data except that we used the GTR substitution model (`--gtr` in VINE), the discrete gamma model for rate variation (`--dgamma 4`), and multithreading (`--parallel 8`). For the analysis of posterior uncertainty (**Supplementary Fig. S16**), VINE was additionally run with `--posterior`. Equivalent options were used in BEAST 2, with BEAGLE employed in this case. For this analysis, we simplified our MCMC convergence criteria for BEAST 2 to require only ESS *>* 400, avoiding the use of pilot replicates and the *R*^^^ criterion.

### Migration mode in VINE

VINE was extended to support the joint model for mutation and migration implemented in BEAM [27]. This version of the model is activated when tissue labels for cells are specified using the `--migration` option. In migration mode, the phylogenetic log likelihood is redefined as a sum of the standard mutation-based log likelihood and a log likelihood based on a general time reversible model for migration transitions, both computed for the same tree and branch lengths. The migration rate parameters are treated as nuisance parameters and estimated by SGA, and the calculation of gradients is modified accordingly. The embedding strategy, conversion to trees by UPGMA, and distance-based initialization required no change. An option (`--primary`, used here for both simulated and real data) allows a particular tissue label to be designated as “primary” and enforced at the root of the tree. After convergence, the program produces samples of tissuelabeled trees under the joint model, by first sampling trees from the approximate posterior and then sampling tissue-labels conditional on each tree using an inside-outside algorithm. Output can be tissue-labeled trees in nexus format (`--labeled-trees`) and/or collapsed migration graphs in dot format (`--sample-graphs`).

To compare tissue-labeled trees across inference methods (**Fig. 6C**), we calculated four phylogenetic statistics: the minimum numbers of (1) mutations and (2) migration events required to explain the data at the tips of the tree according to Fitch-Hartigan parsimony [95, 96]; (3) the cophenetic correlation, defined as Pearson’s correlation (*r*) between the Hamming distances between barcodes and patristic distances along the branches of the tree; and (4) a tissue homogeneity index, defined as the mean fraction of same-tissue neighbors within each clade, normalized by the expected frequency under random tip shuffling. The cophenetic correlation is a measure of how well the tree reflects the raw mutation data, approaching a value of 1 for a perfect correlation [97]. The tissue homogeneity index is a measure of clustering by tissue type similar to measures of cluster purity [98] or phylogenetic trait clustering [99].

### Running BEAM

BEAM was run in two modes: tree-only inference using the sitewise model with uniform fixed edit rates, and joint tree-migration inference that additionally incorporated a GTR migration rate matrix. In both modes, the starting tree was a Cassiopeia-Greedy tree uniformly rescaled to match the appropriate origin time. Markov Chain Monte Carlo (MCMC) was used to sample from the posterior distribution of the tree and other phylogenetic parameters using MCMC proposals that included Wilson-Balding subtree prune and regraft [100] and nearest-neighbor interchange tree moves, internal-node and total-tree-height rescaling, and standard continuous scaling operators on all other continuous parameters. Bactrian proposal moves [101–103] were used where possible, and the frequency and scale of different proposals were tuned adaptively as is standard in BEAST 2 adaptive Metropolis-Hastings MCMC. Priors were shared across both modes where applicable. As in ref. [27], the tree prior was a birth-death process. Other priors included exponential priors (mean 0.1) on the strict clock and silencing rates, an exponential prior (mean 1.0) on the birth-death rate difference, and a Uniform(0,1) prior on the relative death rate. For joint tree-migration inference, migration rates additionally had exponential priors (mean 1.0) and the migration clock rate had an exponential prior (mean 0.1). Parameter initializations were based on test runs for birth/death and clock rates to ensure a stable starting state with sufficient prior density, while migration rates were initialized uniformly. MCMC was run until parameter ESS values exceeded 400 and traces were visually stationary.

### Running other VI programs and LAML

Details on how all other programs were run are provided in the **Supplementary Material**.

### Software implementation

VINE is written in C (C99) and available from github (https://github.com/CshlSiepelLab/vine) under a standard BSD 3-Clause License. It requires a recent installation of PHAST (https://github.com/CshlSiepelLab/ phast) [104], from which it borrows DNA substitution models and alignment-handling routines. VINE is easily installable via bioconda (use `conda` `install` `-c conda-forge -c bioconda vine-phylo`) and homebrew (use `brew` `tap` `CshlSiepelLab/tools` and `brew` `install` `vine`). In both cases, PHAST will be installed as a dependency if needed.

Peak memory usage for VINE in representative benchmarking experiments is shown in **Supplementary Table S2**.

### Code availability

VINE is freely available under the BSD 3-Clause License at https://github.com/CshlSiepelLab/vine. Results using the two-stage procedure were obtained with version 0.4.2 and all other results were obtained with version 0.3.5. Both versions are archived at Zenodo [105]. Scripts for data processing, benchmark execution, and figure generation, with the exact versions of all software used for each experiment pinned, are available at https://github.com/CshlSiepelLab/vine-benchmarks and archived at Zenodo [106].

## Acknowledgments

We thank other members of the community at Cold Spring Harbor Laboratory as well as Dawid Nowak and his team at Weill Cornell Medicine for helpful feedback.

