## Supplementary Information for "VINE: Variational inference for scalable Bayesian reconstruction of species and cell-lineage phylogenies"

### Supplementary Text

#### Backpropagation through the neighbor-joining algorithm

In this section, we outline a strategy for efficient backpropagation of gradients through the neighbor-joining (NJ) algorithm, allowing explicit calculation of the critical third Jacobian, $J_{\text{phy}}$. We will show in the next section that a similar (but simpler) strategy can be used for the UPGMA algorithm as well.

We start with the observation that NJ is piecewise smooth and differentiable almost everywhere, meaning that small changes in the distance matrix $\mathbf{D}$, when propagated through the algorithm, mostly produce smooth changes in the branch lengths $\mathbf{b}$ of the resulting tree. Exceptions occur only when changes in $\mathbf{D}$ alter the sequence of neighbors identified by the algorithm and therefore change the reconstructed tree topology. In other words, NJ is differentiable except for intermittent discontinuities that result from the step in the algorithm where the two neighbors of minimum distance are identified.

We can develop this idea further by decomposing the algorithm into two parts: (1) a sequence of selections of neighbors to be joined in the agglomerative clustering procedure; and (2) a mapping from the distance matrix $\mathbf{D}$ to the branch lengths $\mathbf{b}$ conditional on these neighbor selections. Our intuition is that the most of the information for gradient-based optimization is in this second, smooth component of the problem. We will therefore condition on the sequence of neighbors (1) for a particular value of the embedded points $\mathbf{x}$ and efficiently characterize the continuous mapping (2) given that sequence. We will then rely more heuristically on the general stochasticity of SGA to explore different tree topologies as points migrate in the embedding space. Because it conditions on the selected neighbors, this strategy yields an approximate, topology-conditional gradient of the ELBO: it omits the contribution from shifts in the boundaries between topologies as the variational parameters change, and therefore provides a biased surrogate for the full gradient (see **Methods**).

Recall that NJ operates by iteratively updating two distance matrices, typically denoted $\mathbf{Q}$ and $\mathbf{D}$ (e.g., [1]). These matrices have quite different purposes: $\mathbf{Q}$ is used solely to determine the selection of the neighbors to be joined on each step of the algorithm, and $\mathbf{D}$ is used solely to capture the updated distances between nodes. $\mathbf{Q}$ and $\mathbf{D}$, therefore, map exactly to our components (1) and (2) of the algorithm. Since we condition on the neighbor-selection sequence, $\mathbf{Q}$ plays no further role in the derivation of the Jacobian and we can focus solely on $\mathbf{D}$.

As it turns out, the manner in which NJ updates $\mathbf{D}$ conditional on a sequence of neighbors is remarkably simple. Let us recast the problem in the following way. Let $\mathbf{D}^{(t)}$ denote an $m_t \times m_t$ symmetric distance matrix for iteration $t$ of the algorithm, representing pairwise distances between the members of a set $S_t$ of $m_t$ (internal or external) active nodes. During the course of the algorithm, $\mathbf{D}^{(t)}$ will evolve from the original $n \times n$ distance matrix for the $n$ taxa to a matrix of three remaining nodes at the termination step of the algorithm (at which point the unrooted tree is fully resolved). Thus the number of active nodes, $m_t$, on each step $t$ of the algorithm is given by $m_t = n - t$ for $t = 0, \ldots, n-3$.

Let us first consider the recursive step by which NJ derives $\mathbf{D}^{(t+1)}$ from $\mathbf{D}^{(t)}$ conditional on a specified choice of neighbors $(f, g)$. The NJ algorithm obtains $\mathbf{D}^{(t+1)}$ from $\mathbf{D}^{(t)}$ by defining a new node $u$ representing the cluster of $f$ and $g$, and replacing the rows and columns corresponding to $f$ and $g$ with a single row and column for $u$. The new row and column are then populated by the formula,

$$d_{uk}^{(t+1)} = d_{ku}^{(t+1)} = \frac{1}{2}\big(d_{fk}^{(t)} + d_{gk}^{(t)} - d_{fg}^{(t)}\big), \quad k \in S_t \setminus \{f, g, u\}, \tag{S1}$$

with $d_{uu}^{(t+1)} = 0$ as for all diagonal entries. All other entries in $\mathbf{D}^{(t+1)}$ are left unchanged.

Importantly, this operation is not only smooth and differentiable, it is *linear*. Thus, we can postulate that, if $\mathbf{d}^{(t)} \in \mathbb{R}^{\binom{m_t}{2}}$ is a vectorized representation of $\mathbf{D}^{(t)}$, then there exists a matrix $\mathbf{A}^{(t)}$ such that $\mathbf{d}^{(t+1)} = \mathbf{A}^{(t)}\mathbf{d}^{(t)}$ captures the recursive transformation accomplished by the NJ algorithm. Therefore, the effect of the entire algorithm conditional on a choice of neighbors can be described as a product of such matrices,

$$\mathbf{d}^{(T)} = \mathbf{A}^{(T-1)}\mathbf{A}^{(T-2)} \cdots \mathbf{A}^{(0)}\mathbf{d}^{(0)}, \tag{S2}$$

where $T = n - 3$.

In addition to altering the $\mathbf{Q}$ and $\mathbf{D}$ matrices, the NJ algorithm also must update the branch-lengths of the emerging tree. This operation can also be captured as a recursive matrix update. When the algorithm joins neighbors $f$ and $g$ beneath a new node $u$, it sets the branch lengths between $f$ and $u$ and between $g$ and $u$, respectively, to,

$$b_{fu}^{(t)} = \frac{1}{2}\, d_{fg}^{(t)} + \frac{1}{2(m_t - 2)} \sum_{k \in S_t} \big(d_{fk}^{(t)} - d_{gk}^{(t)}\big),$$
$$b_{gu}^{(t)} = d_{fg}^{(t)} - b_{fu}^{(t)},$$

where the $d_{\cdot\cdot}^{(t)}$ terms are all elements of the current vector $\mathbf{d}^{(t)}$. Again, this is clearly a linear operation that can be abstractly described as,

$$\mathbf{b}^{(t)} = \mathbf{C}^{(t)}\mathbf{d}^{(t)},$$

where $\mathbf{b}^{(t)} = (b_{iu}^{(t)}, b_{ju}^{(t)})'$ and $\mathbf{C}^{(t)}$ is a small matrix.

These $\mathbf{b}^{(t)}$ vectors are disjoint, because the algorithm incrementally defines two new branch lengths per iteration. The final vector of branch lengths, $\mathbf{b}$, can be defined by stacking them together,

$$\mathbf{b} = \begin{bmatrix} \mathbf{b}^{(0)} \\ \mathbf{b}^{(1)} \\ \mathbf{b}^{(2)} \\ \vdots \\ \mathbf{b}^{(T)} \end{bmatrix} = \begin{bmatrix} \mathbf{C}^{(0)}\mathbf{d}^{(0)} \\ \mathbf{C}^{(1)}\mathbf{d}^{(1)} \\ \mathbf{C}^{(2)}\mathbf{d}^{(2)} \\ \vdots \\ \mathbf{C}^{(T)}\mathbf{d}^{(T)} \end{bmatrix} = \begin{bmatrix} \mathbf{C}^{(0)} \\ \mathbf{C}^{(1)}\mathbf{A}^{(0)} \\ \mathbf{C}^{(2)}\mathbf{A}^{(1)}\mathbf{A}^{(0)} \\ \vdots \\ \mathbf{C}^{(T)}\mathbf{A}^{(T-1)}\cdots\mathbf{A}^{(0)} \end{bmatrix} \mathbf{d}^{(0)}, \tag{S3}$$

where the last step uses equation S2. The Jacobian of interest, $J_{\text{phy}}$, follows directly from equation S3:

$$J_{\text{phy}} = \frac{\partial \mathbf{b}}{\partial \mathbf{D}} = \frac{\partial \mathbf{b}}{\partial \mathbf{d}^{(0)}} = \begin{bmatrix} \mathbf{C}^{(0)} \\ \mathbf{C}^{(1)}\mathbf{A}^{(0)} \\ \mathbf{C}^{(2)}\mathbf{A}^{(1)}\mathbf{A}^{(0)} \\ \vdots \\ \mathbf{C}^{(T)}\mathbf{A}^{(T-1)}\cdots\mathbf{A}^{(0)} \end{bmatrix}.$$

This derivation makes the structure of $J_{\text{phy}}$ conceptually clear, but it is not practical to compute the matrices in equation S3 explicitly, because the intermediate Jacobians are large and dense. Instead, we compute the gradient $\nabla_{\mathbf{d}^{(0)}}\ell(\tau, \mathbf{b}; \mathbf{X})$ without forming the Jacobian, using a reverse-mode algorithm that mirrors the NJ updates.

To enable this calculation, we record during the forward execution of NJ each pair of neighbors to be joined, the new internal node that replaces them, and the associated distance values. Later, we propagate the branchwise phylogenetic gradient $\nabla_{\mathbf{b}}\ell(\tau, \mathbf{b}; \mathbf{X})$ back to $\mathbf{d}^{(0)}$ using this recorded "tape" of the forward merges as a guide. In particular, we initialize accumulated gradients for the final two active branches, and then update the gradients on each step $t$ using the chain-rule-based recurrence,

$$\lambda_{fg}^{(t)} \mathrel{+}= \tfrac{1}{2}(\lambda_{t_f} + \lambda_{t_g}), \qquad \lambda_{fk}^{(t)} \mathrel{+}= \frac{\lambda_{t_f} - \lambda_{t_g}}{2(m_t - 2)}, \qquad \lambda_{gk}^{(t)} \mathrel{-}= \frac{\lambda_{t_f} - \lambda_{t_g}}{2(m_t - 2)},$$

for joined neighbors $f, g$, and all active $k \neq f, g$, where $\lambda_{t_f} = \frac{\partial \ell(\tau, \mathbf{b}; \mathbf{X})}{\partial b_{fu}}$ and $\lambda_{t_g} = \frac{\partial \ell(\tau, \mathbf{b}; \mathbf{X})}{\partial b_{gu}}$ are the derivatives with respect to the newly created branch lengths, $b_{fu}$ and $b_{gu}$. These updates are combined with the gradient updates implied by equation S1, which propagate accumulated gradients $\lambda_{uk}^{(t+1)}$ from the new node $u$ to $f$, $g$, and $k$. Because all transformations are linear, this reverse sweep ultimately computes all quantities of the form,

$$\lambda_{ij}^{(0)} = \frac{\partial \ell(\tau, \mathbf{b}; \mathbf{X})}{\partial d_{ij}^{(0)}},$$

which together comprise the desired gradient $\nabla_{\mathbf{d}^{(0)}}\ell(\tau, \mathbf{b}; \mathbf{X})$. Even though some intermediate Jacobians are of size $O(n^3)$, this calculation requires only $O(n^2)$ time and memory, making it asymptotically equivalent to the NJ algorithm itself.

### Calculation of the Jacobian for UPGMA

In the case of the UPGMA algorithm for reconstructing an ultrametric tree, which we use with the CRISPR mutation model, the relationship between the branch lengths and the distance matrix is simpler, and the gradient $\nabla_{\mathbf{d}^{(0)}}\ell(\tau, \mathbf{b}; \mathbf{X})$ can be obtained in a single post-processing traversal of the tree. In this case, the height $h_u$ assigned to an internal node $u$ is simply the average of all intercluster distances between the leaves descending from its left child and those descending from its right child. Therefore, for any pair of leaves $i, j$, the partial derivative of $h_u$ with respect to the distance $d_{ij}$ is given by,

$$\frac{\partial h_u}{\partial d_{ij}} = \begin{cases} \dfrac{1}{2\,|L_u|\,|R_u|}, & \text{if } a \in L_u,\, b \in R_u, \\ 0, & \text{otherwise,} \end{cases}$$

where $L_u$ and $R_u$ denote the leaf sets beneath the left and right children of $u$, respectively. The branch lengths can be obtained as height differences between nodes and their parents: $b_u = h_{\pi(u)} - h_u$, where $\pi(u)$ is the parent of node $u$. The component entries of the branch-length Jacobian $J_{\text{phy}}$ are therefore simply equal to the differences in the corresponding node-height derivatives,

$$\frac{\partial b_u}{\partial d_{ij}} = \frac{\partial h_{\pi(u)}}{\partial d_{ij}} - \frac{\partial h_u}{\partial d_{ij}}.$$

As in the NJ case, we avoid instantiating the full Jacobian by working with accumulated gradients. In particular, for each internal node $u$, the only quantity required by backpropagation is the directional derivative $\sum_{i<j}(\partial b_u/\partial d_{ij})\,\lambda_{ij}$, where $\lambda_{ij}$ is the incoming sensitivity with respect to the pairwise distance $d_{ij}$. Because each node height $h_u$ depends only on the distances between leaves in $L_u$ and $R_u$, this directional derivative can be computed in a single pass over those leaf pairs. Thus the branch-length gradients are obtained by directly accumulating the relevant $1/(2|L_u||R_u|)$ contributions, without forming any intermediate matrices, and the overall cost is linear in the size of the tree.

### Gradients for variance parameterizations

As noted in the **Methods** section, closed-form expression are available for the gradients with respect to the variance parameters of the KLD (equation 9) or entropy (equation 11) and the Jacobian $J_{\text{MVN}}$.

These expressions have simple forms under the CONST and DIST parameterizations. In either case, we can write $\mathbf{\Sigma} = e^{\eta}\mathbf{S}$, where $\mathbf{S} = \mathbf{I}$ for CONST. The gradient of the KLD with respect to the single free parameter $\eta$ for CONST and DIST is therefore,

$$\begin{aligned} \frac{\partial}{\partial \eta}\frac{1}{2}\left[\operatorname{tr}(\mathbf{\Sigma}) - \log|\det \mathbf{\Sigma}|\right] &= \frac{1}{2}\frac{\partial}{\partial \eta}\left[\operatorname{tr}(e^{\eta}\mathbf{S}) - \log|\det(e^{\eta}\mathbf{S})|\right] \\ &= \frac{1}{2}\left[e^{\eta}\cdot \operatorname{tr}(\mathbf{S}) - nd\right], \end{aligned} \tag{S4}$$

because $\log|\det(e^{\eta}\mathbf{S})| = \log(e^{\eta})^{nd} + \log\det(\mathbf{S}) = nd\eta + \log\det(\mathbf{S})$. Under the CONST parameterization, we have the additional simplification that $\operatorname{tr}(\mathbf{S}) = \operatorname{tr}(\mathbf{I}) = nd$, so the entire expression reduces to $\frac{nd}{2}\left(e^{\eta} - 1\right)$. In case (2) (equation 11), we can drop the trace term and both expressions reduce to $-\frac{1}{2}nd$.

In the DIAG case, we can write $\mathbf{\Sigma} = \mathbf{\Lambda}$, with $\mathbf{\Lambda} = \text{diag}\{e^{\eta_1}, \dots, e^{\eta_{nd}}\}$, so that $\text{tr}(\mathbf{\Sigma}) = \sum_i e^{\eta_i}$ and $\log|\det \mathbf{\Sigma}| = \sum_i \eta_i$. Therefore, the partial derivative with respect to each $\eta_i$ is simply,

$$\frac{\partial}{\partial \eta_i} \frac{1}{2} \left[\text{tr}(\mathbf{\Sigma}) - \log|\det \mathbf{\Sigma}|\right] = \frac{1}{2}\left(e^{\eta_i} - 1\right),$$

which reduces to $-\frac{1}{2}$ in case 2.

For the LOWR parameterization, $\mathbf{\Sigma}_0 = \mathbf{R}\mathbf{R}^\top$ introduces non-diagonal structure and requires derivatives with respect to the entries of $\mathbf{R}$. Because $\mathbf{R}$ is $n \times w$ with $w < n$, $\mathbf{\Sigma}_0$ is singular, and (as described in the **Methods**) VINE replaces $\log\det \mathbf{\Sigma}_0$ with the pseudo-determinant $\log\det(\mathbf{R}^\top\mathbf{R})$, corresponding to an intrinsic $wd$-dimensional latent Gaussian. Using standard matrix derivatives, $\partial/\partial\mathbf{R}\,\text{tr}(\mathbf{R}\mathbf{R}^\top) = 2\mathbf{R}$ and $\partial/\partial\mathbf{R}\,\log\det(\mathbf{R}^\top\mathbf{R}) = 2\mathbf{R}(\mathbf{R}^\top\mathbf{R})^{-1}$. Therefore, we can express the gradient in matrix form as,

$$\frac{\partial}{\partial \mathbf{R}} \frac{d}{2}\left[\text{tr}(\mathbf{R}\mathbf{R}^\top) - \log\det(\mathbf{R}^\top\mathbf{R})\right] = d\left(\mathbf{R} - \mathbf{R}(\mathbf{R}^\top\mathbf{R})^{-1}\right),$$

or simply $-d\,\mathbf{R}(\mathbf{R}^\top\mathbf{R})^{-1}$ in case 2.

Evaluation of the Jacobian $J_{\text{MVN}}$ leads to similarly simple expressions. Recall that the reparameterization function $\mathbf{x} = f_{\text{MVN}}(\mathbf{z}) = \boldsymbol{\mu} + \mathbf{L}\mathbf{z}$, where $\mathbf{L}$ is a Cholesky factor for $\mathbf{\Sigma}$. Therefore, the portion of the Jacobian $J_{\text{MVN}}$ pertaining to $\mathbf{\Sigma}$ is given by,

$$\frac{\partial}{\partial \mathbf{\Sigma}}\left[\boldsymbol{\mu} + \mathbf{L}\mathbf{z}\right] = \left(\frac{\partial}{\partial \mathbf{\Sigma}}\mathbf{L}\right)\mathbf{z}.$$

Notice in the CONST case, however, that we can simply write $\mathbf{L} = \sqrt{e^\eta}\,\mathbf{I} = e^{\frac{\eta}{2}}\mathbf{I}$, meaning that,

$$\frac{\partial}{\partial \eta}\left[\boldsymbol{\mu} + \mathbf{L}\mathbf{z}\right] = \frac{1}{2}e^{\frac{\eta}{2}}\mathbf{z}.$$

Similarly, in the DIAG case, $\mathbf{L} = \mathbf{\Lambda}^{\frac{1}{2}}$, where $\mathbf{\Lambda} = \text{diag}\{e^{\eta_1}, \dots, e^{\eta_{nd}}\}$, so for each $\eta_i$,

$$\frac{\partial}{\partial \eta_i}\left[\boldsymbol{\mu} + \mathbf{L}\mathbf{z}\right] = \tfrac{1}{2}e^{\frac{\eta_i}{2}}\mathbf{e}_i.$$

where $\mathbf{e}_i$ denotes the $i$th standard basis vector in $\mathbb{R}^{nd}$, i.e. the vector with a $1$ in position $i$ and $0$ in all other positions.

In the DIST case, if we assume $\mathbf{L} = \sqrt{e^\eta}\mathbf{L}'$, where $\mathbf{L}'$ is a Cholesky factor for the double-centered distance matrix $\mathbf{S}$, then,

$$\frac{\partial}{\partial \eta}\left[\boldsymbol{\mu} + \mathbf{L}\mathbf{z}\right] = \frac{1}{2}e^{\frac{\eta}{2}}\mathbf{L}'\mathbf{z}.$$

For the LOWR parameterization, it is convenient to work at the level of $\mathbf{\Sigma}_0 = \mathbf{R}\mathbf{R}^\top$. For a single embedding dimension $k$, we can write

$$\mathbf{x}^{(k)} = \boldsymbol{\mu}^{(k)} + \mathbf{R}\mathbf{z}^{(k)},$$

where $\mathbf{z}^{(k)} \in \mathbb{R}^w$ collects the $k$th block of latent standard normal variables. The Jacobian of $\mathbf{x}^{(k)}$ with respect to $\mathbf{R}$ is linear in $\mathbf{z}^{(k)}$ and can be expressed elementwise as

$$\frac{\partial \mathbf{x}^{(k)}}{\partial R_{ij}} = z_j^{(k)} \, \mathbf{e}_i,$$

with $\mathbf{e}_i$ the $i$th standard basis vector in $\mathbb{R}^n$. Because the full covariance matrix is $\mathbf{\Sigma} = \mathbf{I}_d \otimes \mathbf{\Sigma}_0$, the complete Jacobian with respect to $\mathbf{R}$ is obtained by stacking these $d$ blocks, one for each embedding dimension.

Notice that under all parameterizations, the Jacobian with respect to $\boldsymbol{\mu}$ is trivial.

$$\frac{\partial}{\partial \boldsymbol{\mu}} [\boldsymbol{\mu} + \mathbf{L}\mathbf{z}] = \mathbf{I}_{nd}$$

where $\mathbf{I}_{nd}$ denotes the $nd \times nd$ identity matrix.

### Jacobians for pairwise distances

As noted in the **Methods** section, the Jacobian $J_{\mathbf{D}}$ for the mapping from embedded points to pairwise distances is needed for backpropagation. This Jacobian can be calculated analytically in a manner that depends on the choice of geometry.

In the case of the Euclidean geometry, the Jacobian of the mapping from points $\mathbf{x}$ to pairwise distances $\{D_{ij}\}$ $(\partial \mathbf{D})/\partial \mathbf{x})$ has a simple block structure. For each pair $(i, j)$,

$$\frac{\partial D_{ij}}{\partial \mathbf{x}_i} = \frac{\mathbf{x}_i - \mathbf{x}_j}{s \, \|\mathbf{x}_i - \mathbf{x}_j\|}, \qquad \frac{\partial D_{ij}}{\partial \mathbf{x}_j} = -\frac{\partial D_{ij}}{\partial \mathbf{x}_i},$$

with all other partial derivatives equal to zero.

In the hyperbolic case, the Jacobian blocks of the mapping are,

$$\frac{\partial D_{ij}}{\partial \mathbf{x}_i} = c_{ij} \left( \frac{x_{0,j}}{x_{0,i}} \mathbf{x}_i - \mathbf{x}_j \right), \qquad \frac{\partial D_{ij}}{\partial \mathbf{x}_j} = c_{ij} \left( \frac{x_{0,i}}{x_{0,j}} \mathbf{x}_j - \mathbf{x}_i \right),$$

again with all other partial derivatives equal to zero. Here, $c_{ij}$ is defined as,

$$c_{ij} = \frac{\alpha}{s} \frac{1}{\sqrt{u_{ij}^2 - 1}}, \qquad u_{ij} = x_{0,i} x_{0,j} - \mathbf{x}_i^\top \mathbf{x}_j.$$

### Efficient calculation of branch-length gradients

Backpropagation begins with the gradient of the phylogenetic log likelihood function with respect to the branch lengths, $\nabla_{\mathbf{b}} \ell(\tau, \mathbf{b}; \mathbf{X})$. A naive approach to obtaining this gradient would require $2n - 2$ evaluations of the likelihood function, for $O(n^2)$ total time, but the gradient can be computed in linear time by taking advantage of an "inside/outside" algorithm along the tree (see, e.g., [2]). Briefly, at each site $i$, Felsenstein's pruning algorithm is first performed in a post-order traversal of the tree, producing quantities of the form,

$$P(L_v \mid v = b),$$

at all nodes $v$, where $L_v$ represents the data at the leaves beneath node $v$ and $v = b$ indicates the condition that node $v$ has nucleotide identity (or CRISPR mutation) $b$. This procedure is followed by a similar algorithm that proceeds in pre-order, from root to leaves, and computes the complementary quantities,

$$P(L_{\bar{v}}, v = b),$$

where $L_{\bar{v}}$ indicates the data at all leaves *other than* the ones beneath $v$. The partial derivatives of interest can then be computed in a final traversal of the tree using the property that, for a node $v$, its parent $u$, and its sibling $w$,

$$\frac{\partial}{\partial b_v} \log P(\mathbf{X}_i \mid \tau, \mathbf{b}) = \frac{1}{P(\mathbf{X}_i \mid \tau, \mathbf{b})} \sum_a \sum_b \sum_c p(L_{\bar{u}}, u = a)\, p(L_v \mid v = b)\, p(L_w \mid w = c) \\ \times P(a \to c \mid b_w)\, \frac{\partial}{\partial b_v} P(a \to b \mid b_v),$$

where $\log P(\mathbf{X}_i \mid \tau, \mathbf{b})$ is the contribution to the log likelihood at site $i$, $P(a \to c \mid b_w)$ denotes the probability of an $a \to c$ substitution along a branch of length $b_w$ and $\frac{\partial}{\partial b_v} P(a \to b \mid b_v)$ denotes the partial derivative with respect to the branch length of the probability of an $a \to b$ substitution along a branch of length $b_v$, which can be calculated analytically in a manner dependent on the assumed substitution model. These terms are then summed across all sites $i$ for each branch. Their computation is carried out together with the log likelihood calculation itself for minimal overhead. Note that dynamic rescaling of the inside and outside quantities is needed to avoid numerical underflow on large trees.

### Priors for trees

By default, VINE assumes a prior in the embedding space of $p(\mathbf{x}) = \mathrm{MVN}(0, \mathbf{I})$, which, as discussed in the **Methods** section, is typical for variational autoencoders. The program also allows specification, however, of more informative priors for phylogenies. If these are selected, the KLD can no longer be evaluated in closed form and the prior must be incorporated into Monte Carlo estimation (or Taylor approximation) along with the log likelihood (see equation 7). In particular, the `--treeprior` option allows for either Yule prior over tree topologies (integrating over birth rates) or a Gamma model for total branch length; and the `--relclock` option allows for a relaxed local clock with lognormal clock rates per branch. In these cases, any free parameters of the priors are also considered nuisance parameters during optimization. Evaluation of priors was not a focus of this study, however, and all of our results reflect the default $\mathrm{MVN}$ prior.

### Details on running other programs

We ran VaiPhy with the arguments `--max_iter 200` and `--n_particles 128` as the authors used for their benchmarking experiments [3]. For GeoPhy and VBPI-GNN, we used scripts conveniently provided by Zhou et al. [4]. For GeoPhy we used commands such as,

```
python ./scripts/run_gp.py -v -ip ds-data/ds1/DS1.nex -op results/ds1/ds1_paper
-c ./config/default.yaml -s 0 -ua -es lorentz -edt full -ed 4 -eqs 1e-1 -ast
100_1000 -mcs 3 -ci 1000 -ms 1000_000 -ul
```

which is identical to that reported in ref. [4] except that we reduced the number of steps (`-ms`) from 1,000,000 to 100,000, which appeared to be adequate for convergence in our case. For VBPI-GNN, we used commands such as,

```
python main.py −dataset DS1 −brlen_model gnn −gnn_type edge −hL 2 −hdim 100
−maxIter 400000 −empFreq −psp!
```

We followed Zhou et al. [4] in using IQ-TREE2 [5] for pre-generation of trees, using commands such as:

```
iqtree -s DS1 -bb 10000 -wbt -m JC69 -redo
```

We ran LAML (v1.0.5) with commands such as,

```
run_laml -c quinn.31.csv -t quinn.31.cass.nwk -o quinn.31 −topology_search
−noDropout
```

where `quinn.31.cass.nwk` is a starting tree produced using the Cassiopeia-Greedy algorithm:

```
python cassiopeiaGreedy.py quinn.31.csv quinn.31.cass.nwk
```

All computational experiments were run on a ∼6 yr-old HPE ProLiant DL380 Gen10 server equipped with dual Intel Xeon Gold 5120 processors (28 physical cores total, 56 hardware threads) and 192 GB of RAM, running Rocky Linux 8.10. No GPU acceleration was used.

### Supplementary Tables

| dataset | No. of taxa | No. of sites | Type of data | TreeBase accno. |
|---:|---:|---:|---|---|
| ds1 | 27 | 1949 | rRNA, 18s | M336 |
| ds2 | 29 | 2520 | rDNA, 18s | M501 |
| ds3 | 36 | 1812 | mtDNA, COII (1–678), cytb (679–1812) | M1510 |
| ds4 | 41 | 1137 | rDNA, 18s | M1366 |
| ds5 | 43 | 1660 | rDNA, 18s | M932 |
| ds6 | 50 | 378 | Nuclear protein coding, wingless | M3475 |
| ds7 | 50 | 1133 | rDNA, 18s | M1044 |
| ds8 | 59 | 1824 | mtDNA, COII and cytb | M1809 |

**Supplementary Table S1:** Real data used for benchmarking of DNA substitution models. Reproduced from Table 1 of ref. [6].

| Data type | Length | Source | No. taxa | Peak mem. (MB) |
|---|---|---|---|---|
| DNA | 300 | Simulated | 10 | 18.2 |
| DNA | 300 | Simulated | 25 | 25.6 |
| DNA | 300 | Simulated | 50 | 44.9 |
| DNA | 300 | Simulated | 100 | 85.5 |
| DNA | 300 | Simulated | 250 | 230.8 |
| DNA | 300 | Simulated | 500 | 593.1 |
| DNA | 300 | Simulated | 1000 | 1578.3 |
| DNA | 10000 | Simulated | 10 | 15.4 |
| DNA | 10000 | Simulated | 25 | 27.4 |
| DNA | 10000 | Simulated | 50 | 47.6 |
| DNA | 10000 | Simulated | 100 | 88.9 |
| DNA | variable | Real (Lakner et al.) | variable | 56.1 |
| DNA | 29903 | Real (SARS-CoV-2) | 364 | 396.5 |
| DNA | 29903 | Real (SARS-CoV-2) | 1060 | 2134.8 |
| CRISPR | 30 | Simulated | 10 | 7.7 |
| CRISPR | 30 | Simulated | 25 | 9.1 |
| CRISPR | 30 | Simulated | 50 | 12.7 |
| CRISPR | 30 | Simulated | 100 | 20.5 |
| CRISPR | 30 | Simulated | 250 | 55.9 |
| CRISPR | 30 | Simulated | 500 | 158.3 |
| CRISPR | 30 | Simulated | 1000 | 474.4 |
| CRISPR | variable | Real (Quinn et al.) | variable | 784.0 |

**Supplementary Table S2:** Peak memory usage of VINE for representative benchmarking experiments. For the simulation experiments, average values of peak memory across replicates are shown. For the real data sets, maximum values across datasets of various size are shown. The CRISPR results here reflect tree inference only (not tissue migration), but migration inference does not have a major effect on memory usage.

### Supplementary Figures

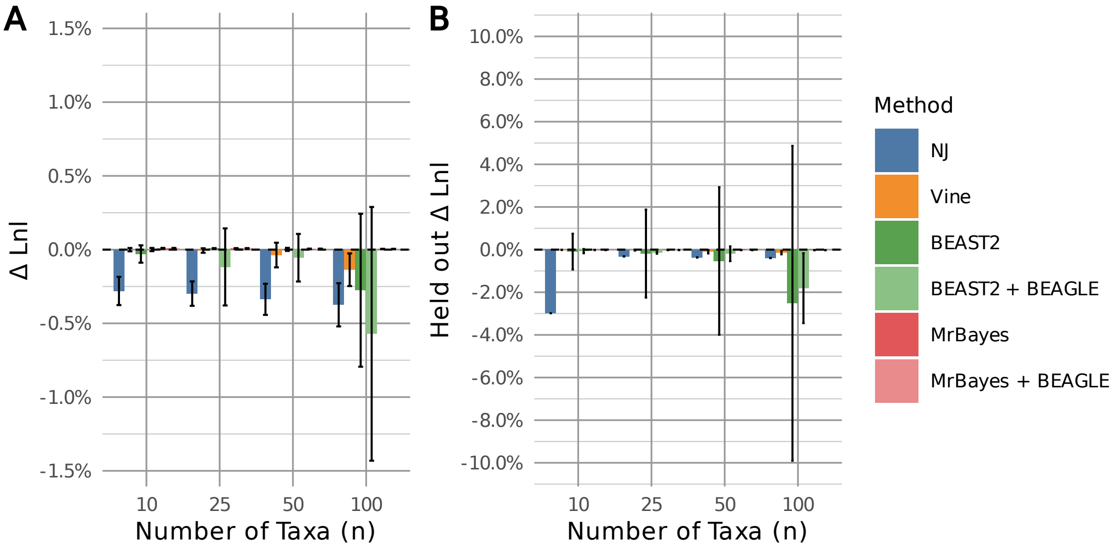


**Supplementary Figure S1:** Measures of model fit for phylogenetic models estimated from simulated data by VINE and other methods, for various numbers of taxa $n$, as in **Fig. 2** but for alignments of 10,000 nucleotides. **(A)** Maximized log likelihood during model fitting, relative to the log likelihood of the true (generating) model (zero line). **(B)** Average log likelihood across posterior samples for held-out data, also relative to the true model. Error bars represent one standard deviation. VINE (*orange*) was applied with default parameters. Results are also shown for neighbor-joining (*blue*), BEAST 2 (*green*), and MrBayes (*red*); variants of BEAST 2 and MrBayes with BEAGLE are also shown (lighter shades).

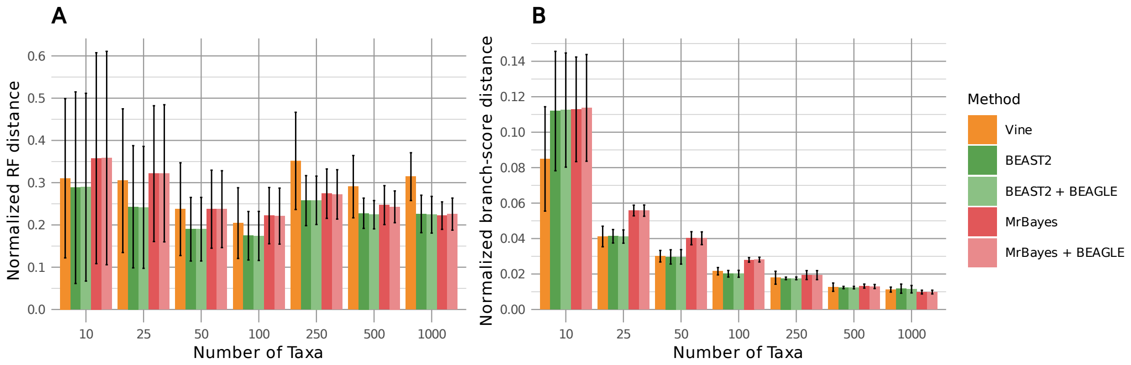


**Supplementary Figure S2:** Accuracy of tree topologies and branch lengths inferred from simulated data by VINE and other methods, for various numbers of taxa $n$. **(A)** Topological accuracy, measured by the normalized Robinson-Foulds distance [7] from the true tree. **(B)** Branch-length accuracy, measured by the branch-score distance [8] from the true tree—the Euclidean distance between the two trees' vectors of branch lengths indexed by splits, including terminal branches—normalized by the total branch length of the true tree. To isolate branch-length accuracy from posterior dispersion, the branch-score distance was computed for each method's posterior-mean tree rather than averaged over the posterior. In both panels, results are shown for 300 bp alignments and estimation under the HKY substitution model, with ten replicates per bar, and error bars representing one standard deviation. VINE (*orange*) was applied with default parameters. Results are also shown for BEAST 2 (*green*) and MrBayes (*red*); variants with BEAGLE are also shown (lighter shades) for completeness.

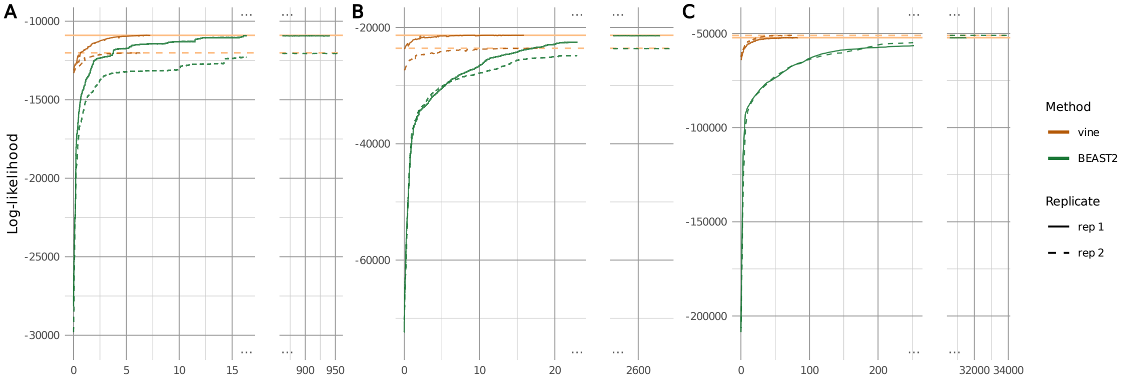


**Supplementary Figure S3:** Convergence trajectories for VINE (*orange*) and BEAST 2 (*green*) on simulated DNA alignments with **(A)** 50, **(B)** 100, and **(C)** 250 taxa. Separate trajectories for the reported log likelihood are shown for replicate 1 (solid lines) and replicate 2 (dashed lines) for the experiments based on the HKY model and 300 bp alignments. Because VINE converges much more quickly, the final points in its trajectories are extended to the right sides of the plots in a lighter shade of *orange*. Notably, BEAST 2 continues to run well after it has appeared to converge in order to ensure that the target effective sample size (ESS) of 400 is achieved. VINE additionally benefits from a better starting point, based on an initial NJ tree. The log likelihood is shown for both methods to allow for a direct comparison, even though the ELBO and log posterior are the quantities of more direct interest for VI- and MCMC-based methods, respectively (values are similar on this scale).

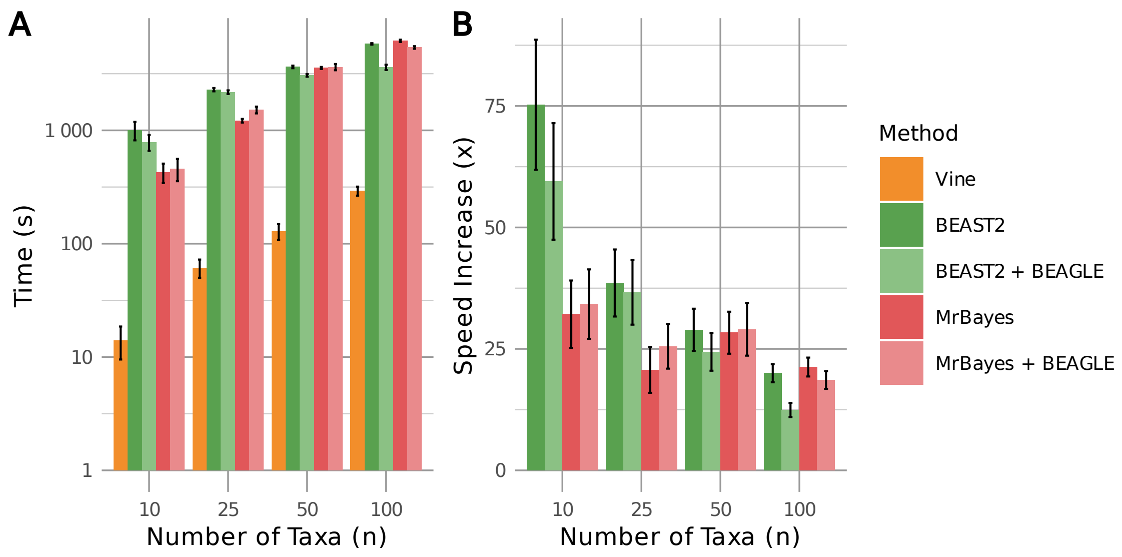


**Supplementary Figure S4:** Compute time required for experiments described in **Supplementary Fig. S1**, in seconds per replicate, as in **Fig. 2** but for alignments of 10,000 nucleotides. **(A)** Results for VINE and MCMC-based methods. **(B)** Speed increase of VINE relative to BEAST 2 and MrBayes (shown with and without BEAGLE). Results are shown for VINE (*orange*), BEAST 2 (*green*), and MrBayes (*red*); variants with BEAGLE are shown in lighter shades.

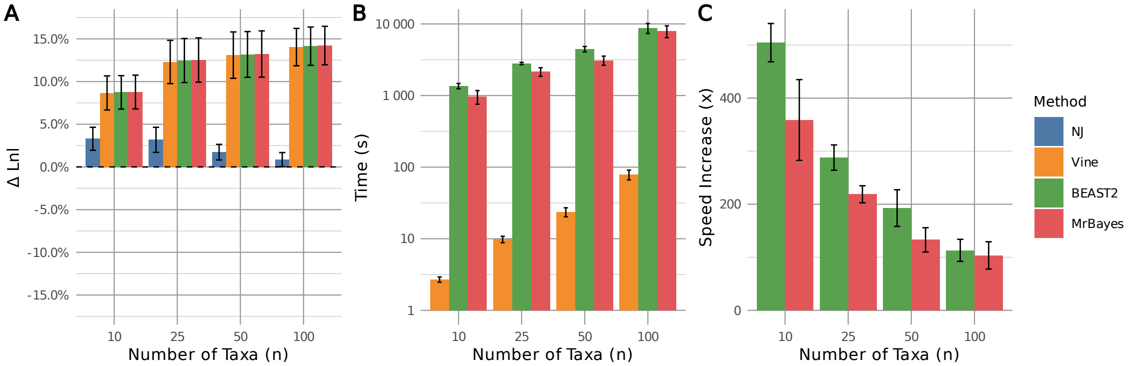


**Supplementary Figure S5:** Performance of VI and MCMC-based methods on DNA data simulated under the general time reversible model with gamma-distributed rate variation (GTR+G), with high variability in rates ($\alpha = 0.5$). **(A)** Maximized log likelihood during model fitting, relative to the log likelihood of the true (generating) model (zero line). Inference was performed under the GTR substitution model and the discrete gamma model for rate variation, with four rate categories. Notice that free estimation of $\alpha$ in this case allows all models to improve substantially on the true model (c.f., **Fig. 2E**). **(B)** Compute time required when running without parallelization or GPU acceleration on an HPE ProLiant DL380 Gen10 server (see **Methods**), in seconds per replicate (note log scale). **(C)** Speed increase of VINE relative to BEAST 2 and MrBayes. Results are shown for neighbor-joining (*blue*), VINE (*orange*), BEAST 2 (*green*), and MrBayes (*red*).

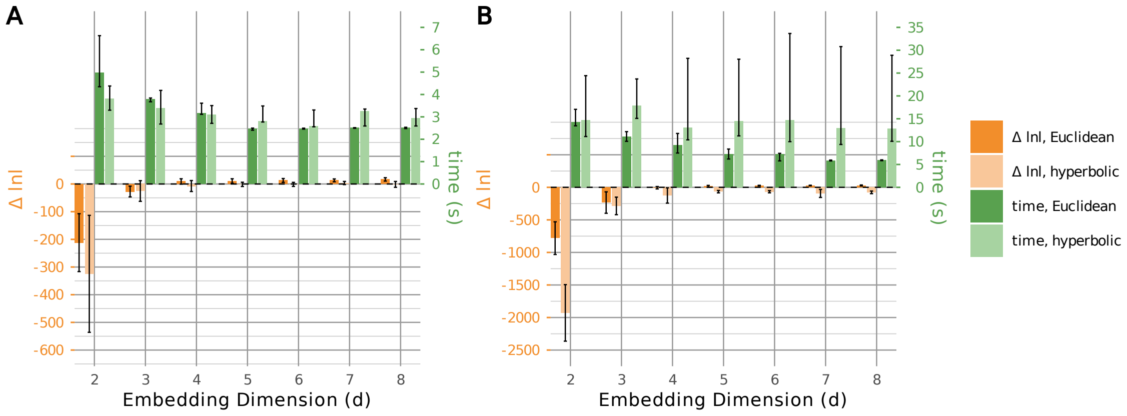


**Supplementary Figure S6:** Performance improvement of VINE with increasing dimensionality $d$ of the embedding space, comparing Euclidean and hyperbolic geometries. Shown are running times per replicate (*green*, right axis) and deviations of the maximized log likelihood from that of the true model (*orange*, left axis) for **(A)** $n = 25$ taxa and **(B)** $n = 50$ taxa. In each panel, darker bars denote the Euclidean geometry and lighter bars denote the hyperbolic geometry. Log-likelihood deviations are summarized by their mean $\pm$ one standard deviation over ten replicates; running times, whose distribution is heavily right-skewed under the hyperbolic geometry, are summarized by their median and interquartile range. Results are for alignments of 300 bp and estimation under the HKY model with other parameters at their default values. The number of free parameters under this parameterization is $nd + 2$, so it ranges from 52–202 in (A) and 102–402 in (B).

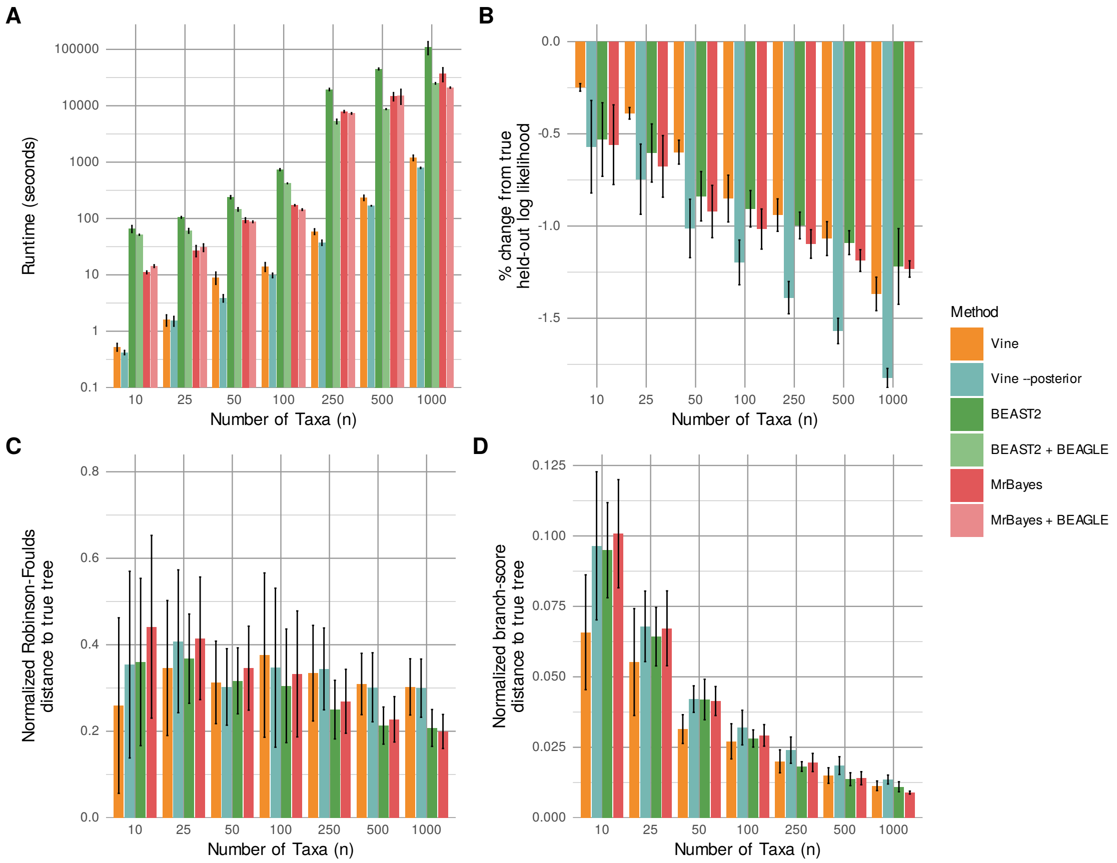


**Supplementary Figure S7:** Additional measures of performance for programs considered in evaluations of posterior accuracy (**Fig. 3B&C**). **(A)** Compute time required in seconds per replicate (note log scale). **(B)** Average log likelihood across posterior samples for held-out data. **(C)** Normalized Robinson-Foulds distance from the true tree. **(D)** Normalized branch-score distance from the true tree. Results are shown for 300-bp HKY-based alignments for VINE with default parameters (VINE; *orange*), VINE with two-stage estimation of posterior uncertainty (VINE `--posterior`; see **Methods**; *turquoise*), BEAST 2 (*green*), and MrBayes (*red*). In panel (A), results are shown for versions of BEAST 2 and MrBayes with and without the use of BEAGLE(lighter shades).

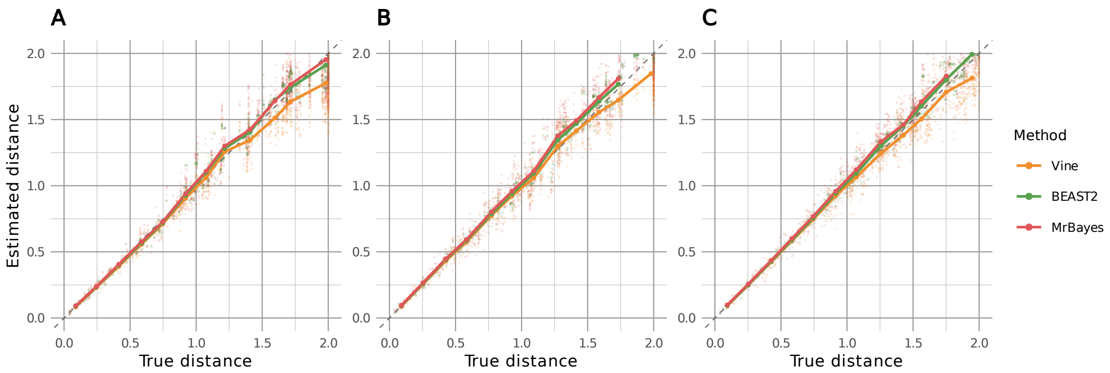


**Supplementary Figure S8:** True vs. estimated pairwise distances based on reconstructions of simulated trees under the HKY model (300 sites). Results are shown for 10 replicates each of trees for **(A)** 25 taxa, **(B)** 50 taxa, and **(C)** 100 taxa. Estimates represent posterior mean values across sampled trees from the approximate posterior and corresponding true distances reflect the ground truth in simulation (both in units of substitutions/site). For legibility, points are shown for 1,500 randomly sampled pairwise distances in each panel (out of $10 \times \binom{n}{2} =$ 3,000, 12,250, and 49,500, respectively). The solid lines represent mean estimates within a sliding window along the $x$ axis. The dashed line is the target of $y = x$. Results are shown for VINE (*orange*), BEAST 2 (*green*), and MrBayes (*red*). Notice the slight underestimation of VINE at large distances.

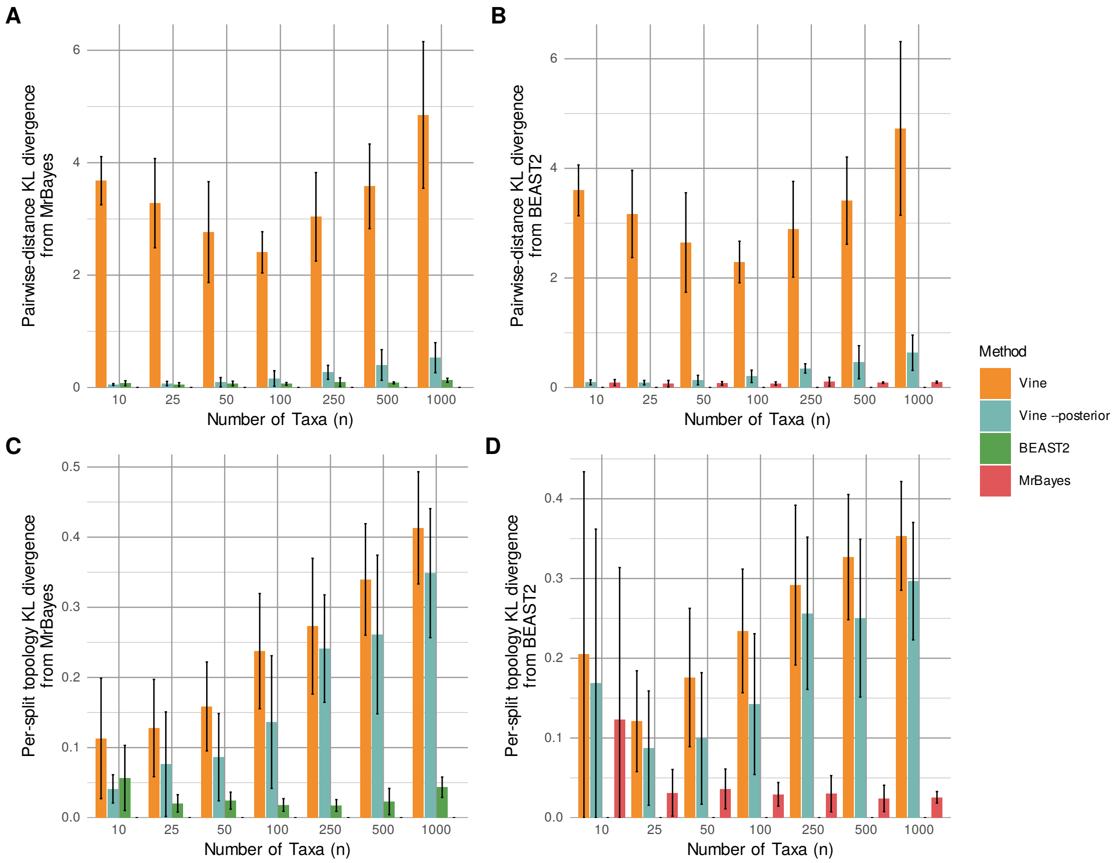


**Supplementary Figure S9:** Average marginal Kullback–Leibler (KL) discrepancies of VINE's posterior approximation from MCMC-based reference distributions. **(A)** and **(B)** show the average marginal distance-KL discrepancy, based on induced distributions of pairwise distances, whereas **(C)** and **(D)** show the average marginal split-KL discrepancy, based on the relative frequencies of all possible topological "splits" (bipartitions of taxa). In **(A)** and **(C)**, samples from MrBayes are used as the reference distribution, and in **(B)** and **(D)**, samples from BEAST 2 are used as the reference distribution. Results are shown for VINE (*orange*), VINE `--posterior` (*turquoise*), BEAST 2 (*green*), and MrBayes (*red*); VINE `--posterior` denotes two-stage estimation of posterior uncertainty (see **Methods**). These measures are averages over marginal distributions, not KL divergences between full tree posteriors; see **Methods** for details.

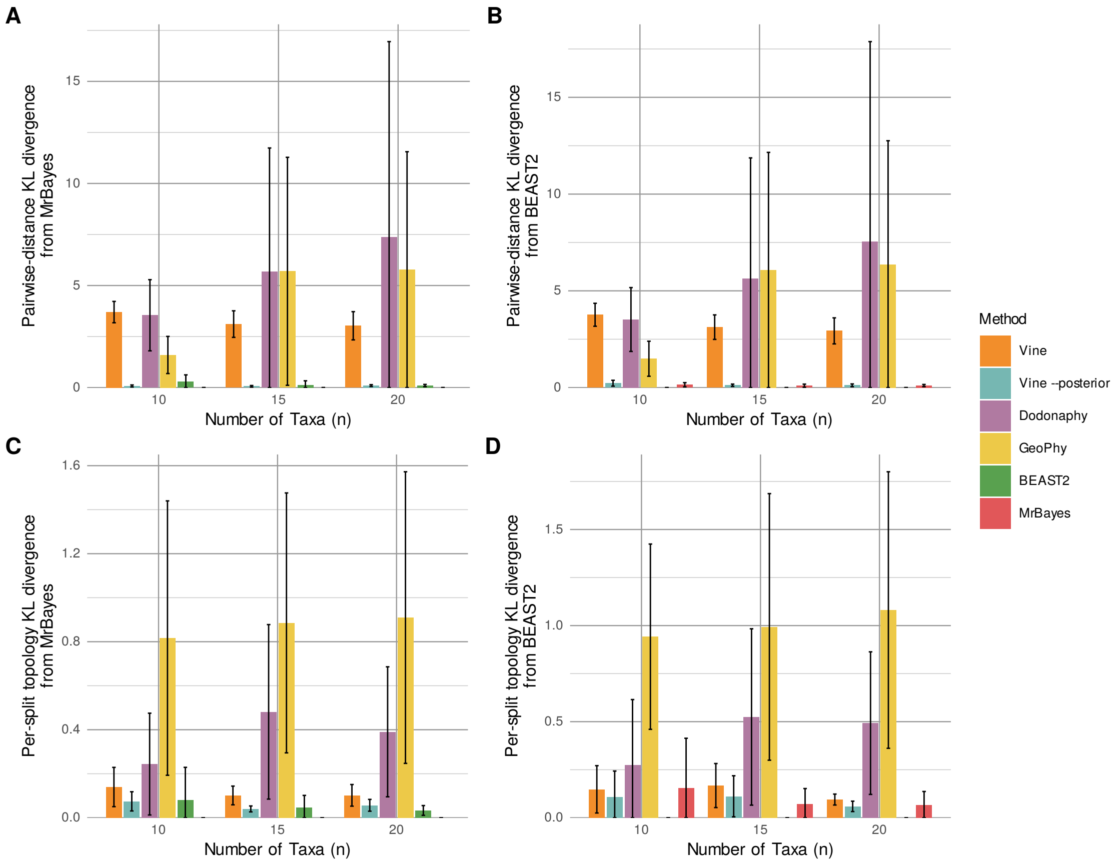


**Supplementary Figure S10:** Average marginal Kullback–Leibler (KL) discrepancies of posterior approximations relative to MCMC-based reference distributions including other VI-based methods. This figure is similar to **Supplementary Fig. S9** but includes the VI-based methods Dodonaphy and GeoPhy. Experiments were performed on 300-bp simulated alignments for $n \in \{10, 15, 20\}$ taxa. For all methods, inference was performed under the Jukes-Cantor (JC) model. Results are shown for VINE (*orange*), VINE `--posterior` (*turquoise*), BEAST 2 (*green*), MrBayes (*red*), Dodonaphy (*purple*), and GeoPhy (*yellow*). VINE `--posterior` denotes two-stage estimation of posterior uncertainty (see **Methods**).

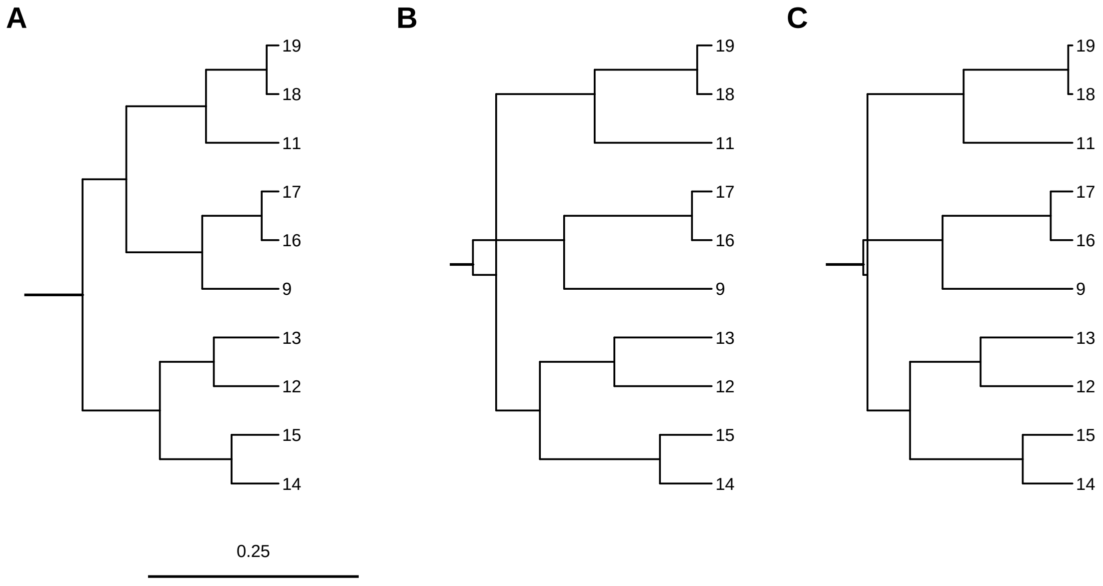


**Supplementary Figure S11:** Reconstruction of **(A)** a simulated 10-taxon tree by **(B)** VINE and **(C)** LAML based on a simulated CRISPR-barcode mutation matrix. Horizontal branch lengths are drawn to scale (see scale bar). Here the VINE tree represents the MCC summary and the LAML tree is the reported ML estimate. Notice that the trees are identical except that both VINE and LAML change the branching order for the clade consisting of taxa 9, 16, and 17 (see crossing branches). VINE was run with default parameter values.

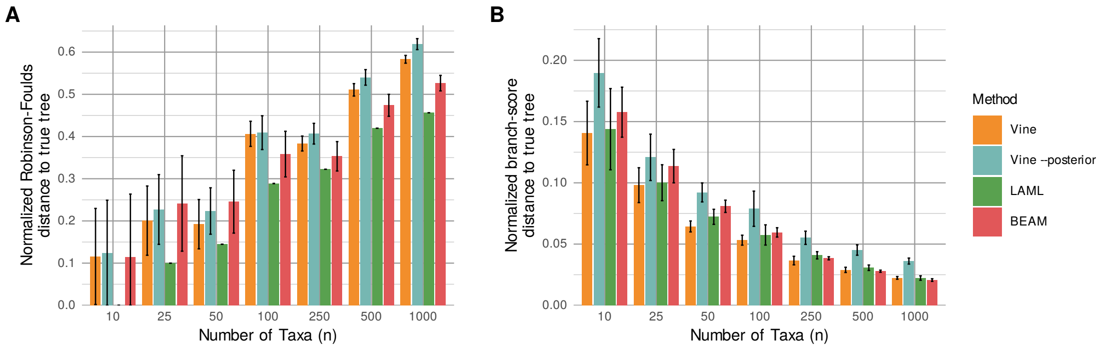


**Supplementary Figure S12:** Accuracy of trees inferred from simulated CRISPR data by VINE (*orange*), VINE `--posterior` (*turquoise*), LAML (*green*), and BEAM (*red*) for various numbers of taxa $n$. **(A)** Topological accuracy, measured by the normalized Robinson-Foulds distance [7] from the true tree. **(B)** Branch-length accuracy, measured by the branch-score distance [8] from the true tree—the Euclidean distance between the two trees' vectors of branch lengths indexed by splits, including terminal branches—normalized by the total branch length of the true tree. True trees, which are simulated in units of time, were converted to expected edits per site by multiplying branch lengths by the simulated edit rate (0.01) before computing branch-score distances (see **Methods**).

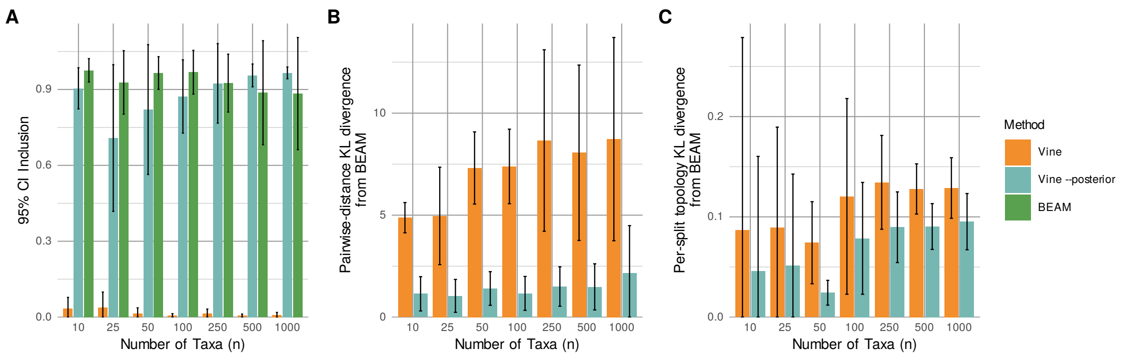


**Supplementary Figure S13:** Accuracy of VINE's approximations of posterior distributions of trees inferred from simulated CRISPR data for various numbers of taxa $n$. **(A)** Fraction of all pairwise distances between taxa that fall within the estimated 95% credible interval (95% CI Inclusion) for VINE with default parameters (*orange*), VINE `--posterior` (two-stage estimation of posterior uncertainty; see **Methods**) (*turquoise*), and the MCMC-based BEAM method (*green*). **(B)** Average marginal distance-KL discrepancy of VINE (with and without `--posterior`) from BEAM's posterior distribution, computed from the marginal distributions of distances for all pairs of taxa. **(C)** Average marginal split-KL discrepancy relative to BEAM, computed from the relative frequencies of each split (bipartition of taxa) in the sampled topologies. True pairwise distances were converted from units of time to expected edits per site using the simulated edit rate (0.01), and BEAM's samples were converted by the same factor for the distance-KL discrepancy (panel B).

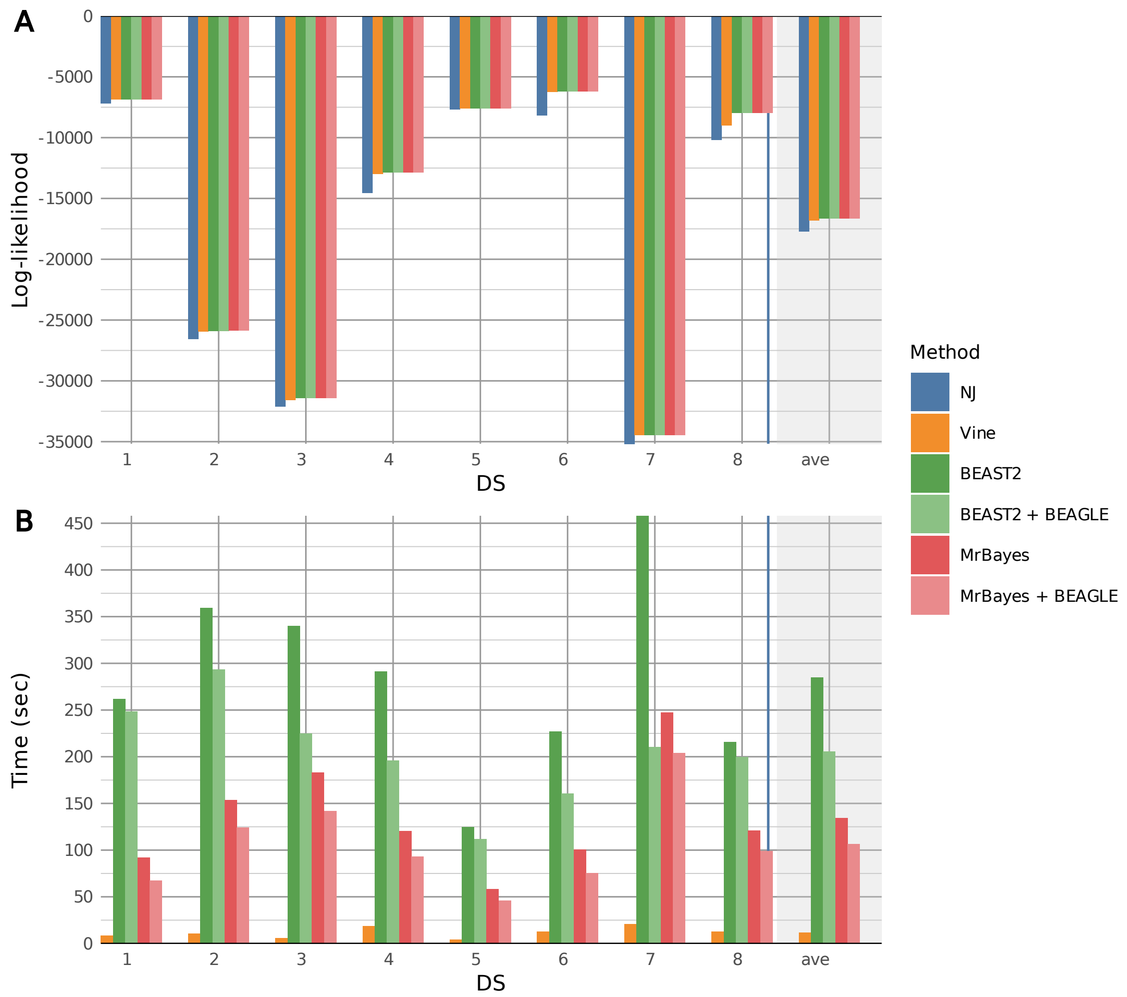


**Supplementary Figure S14:** **(A)** Relative maximized log likelihoods during model fitting for the alignments from ref. [6] (**Supplementary Table S1**) under the HKY substitution model. Results are shown for neighbor-joining (*blue*), VINE (*orange*), BEAST 2 (*green*), and MrBayes (*red*) (with and without BEAGLE). The zero line represents the average of all four methods. **(B)** Compute time required for the same alignments. In both (A) and (B), the last column (*gray* background) shows average values across data sets. VINE was run with default parameter values.

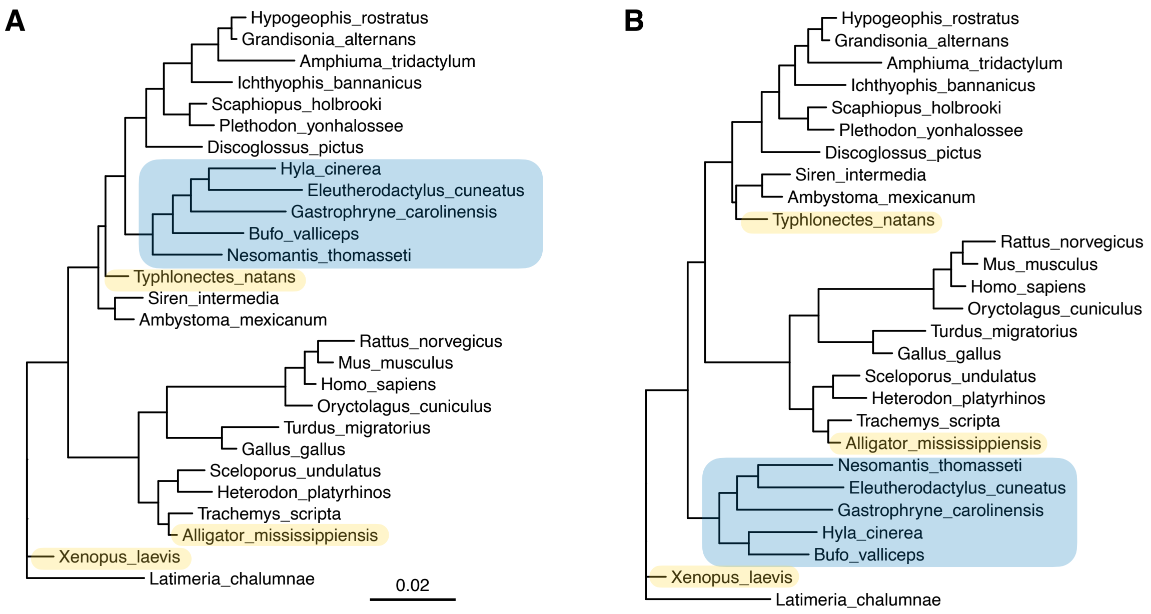


**Supplementary Figure S15:** Examples of reconstructed phylogenies for real alignments of nucleic acid sequences by **(A)** VINE and **(B)** BEAST 2 under the HKY model. Results are for dataset 1 (ds1) from ref. [6] (details in **Supplementary Table S1**). As in **Fig. 2C&D**, samples from the posterior distributions are summarized using maximum-clade-credibility (MCC) trees with horizontal branch lengths drawn to scale (scale bar). Notice that the two trees are generally in agreement, with the exception of the placement and precise arrangement of the neobatrachian frogs (highlighted in *blue*), which are incorrect in both cases (they should group with *Discoglossus pictus* and *Xenopus laevis*). In addition, three other taxa (*yellow*) are placed incorrectly in both trees relative to the accepted phylogeny, not surprisingly for a tree based only on highly constrained 18s rRNA sequences. VINE was run with default parameter values.

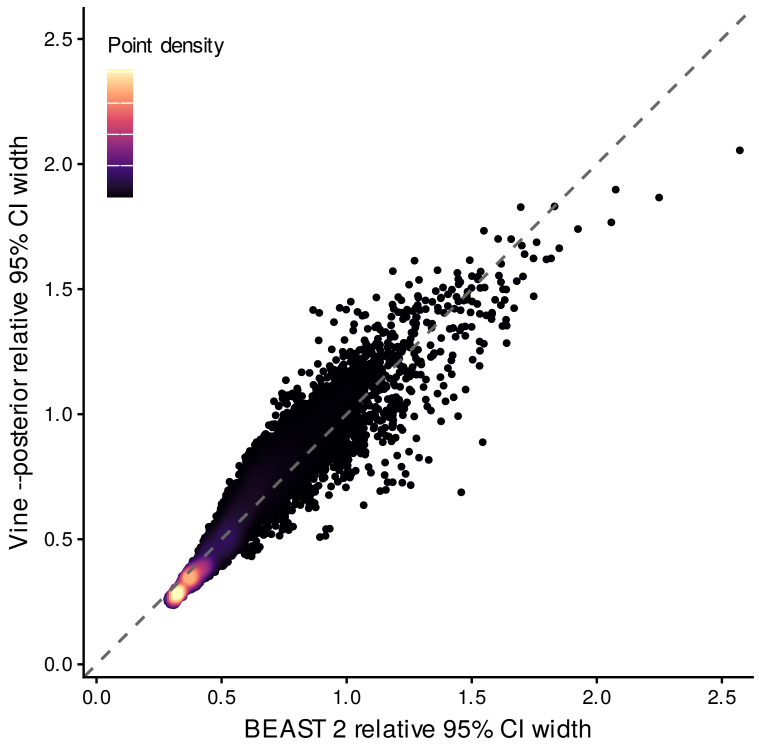


**Supplementary Figure S16:** Posterior uncertainty for the 364-taxon SARS-CoV-2 dataset. Each point represents the width of the 95% CI divided by posterior mean for the patristic distance corresponding to one of the 66,066 taxon pairs under BEAST 2 ($x$) and VINE ($y$). Points are colored by density and the dashed line represents $y = x$. Results are based on 1,000 samples from VINE and 8,000 post-burn-in samples from BEAST 2. VINE was run with `--posterior` (two-stage estimation of posterior uncertainty; see **Methods**).

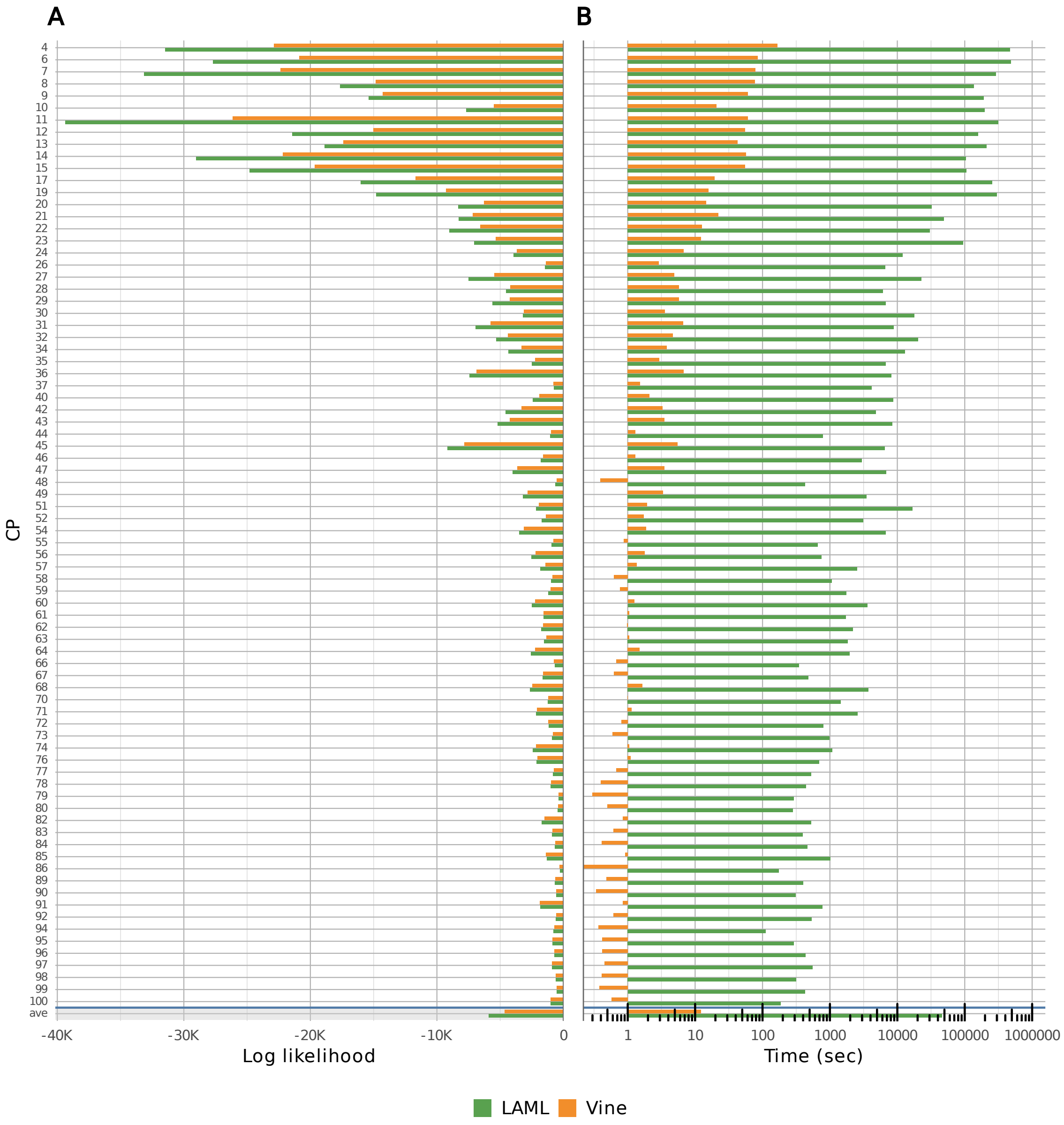


**Supplementary Figure S17:** **(A)** Maximized log likelihoods for the CRISPR-based mutation matrices for 80 clonal populations (CP) from ref. [9]. Results are shown for VINE (*orange*) and LAML (*green*). **(B)** Compute time required for the same mutation matrices (note log scale). In both (A) and (B), the last row (*gray* background) shows average values across datasets. VINE was run with default parameter values. The largest three clonal populations from ref. [9] (CPs 1–3) were excluded because they consisted of >5000 cells, making inference with LAML computationally infeasible.

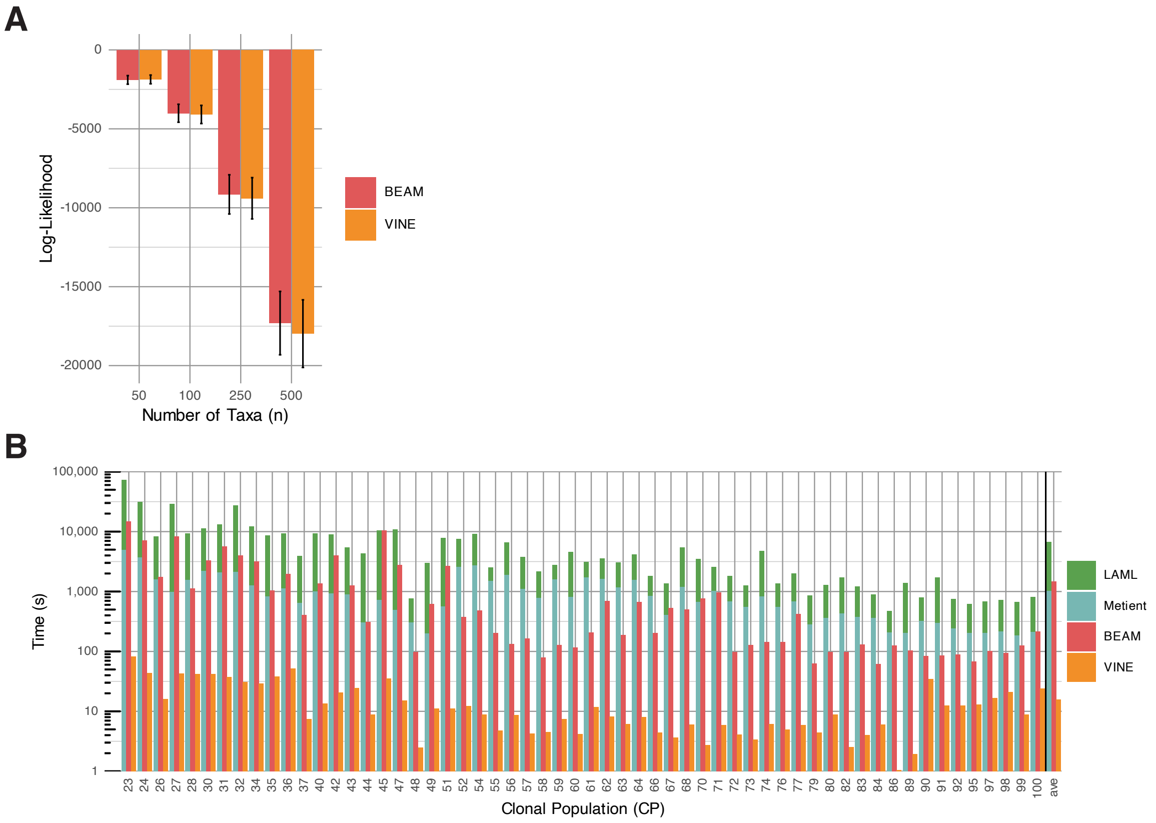


**Supplementary Figure S18:** **(A)** Maximized joint log likelihoods (mutation + migration) for BEAM (*red*) and VINE (*orange*) on simulated tissue-migration datasets of various sizes. **(B)** Compute time required for the same simulated tissue-migration data (note log scale), for BEAM, VINE, and Metient (*turquoise*). The time required to obtain an input tree with LAML (*green*) is shown together with the Metient values.

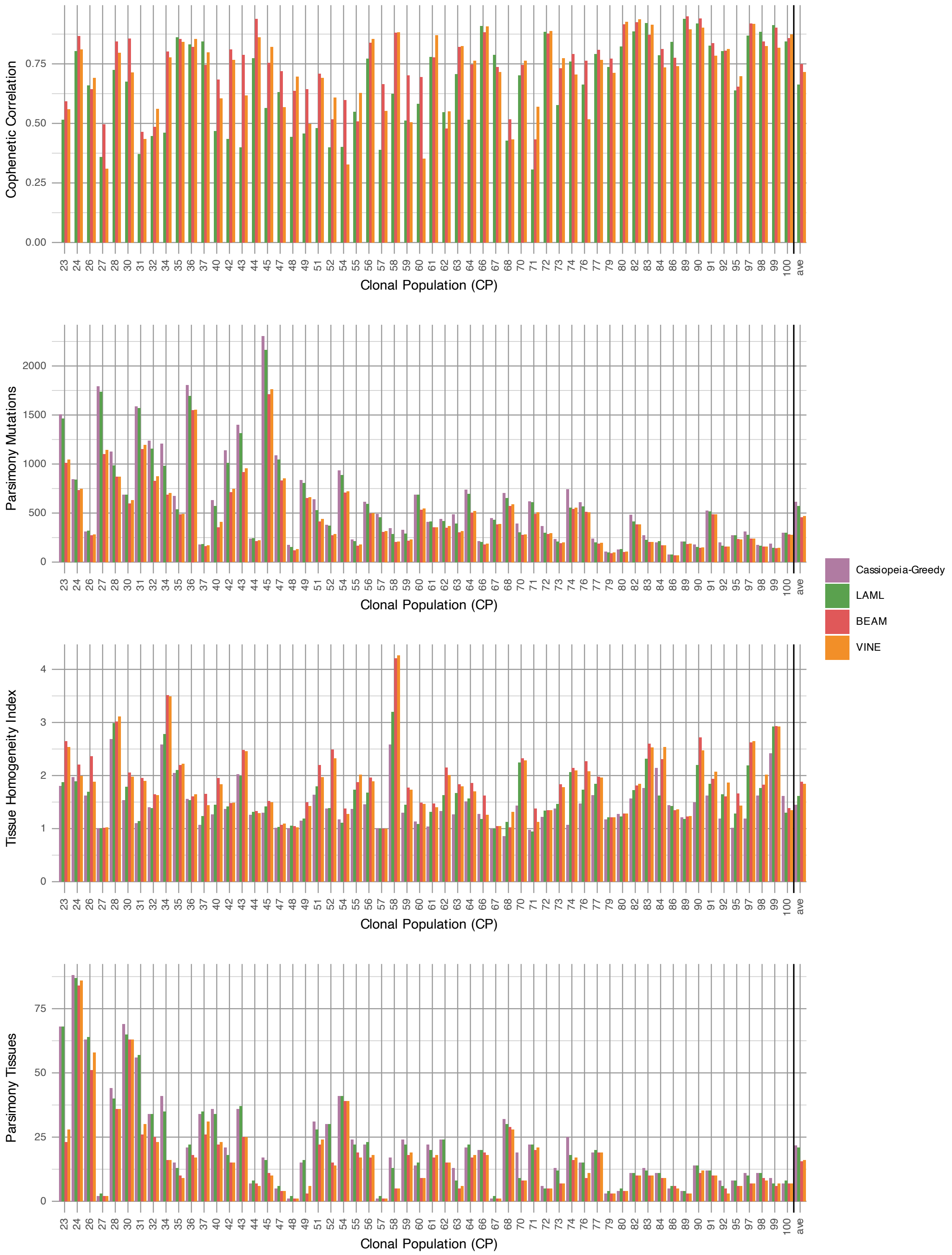


**Supplementary Figure S19:** Cophenetic correlation, parsimony mutation count, tissue homogeneity index, and parsimony migration count (see **Methods** for definitions) for trees inferred by Cassiopeia-Greedy (*mauve*), LAML (*green*), BEAM (*red*), and VINE (*orange*) for Clonal Populations (CPs) from ref. [9]. The largest CPs ($< 23$) were excluded here for reasons of computational efficiency.

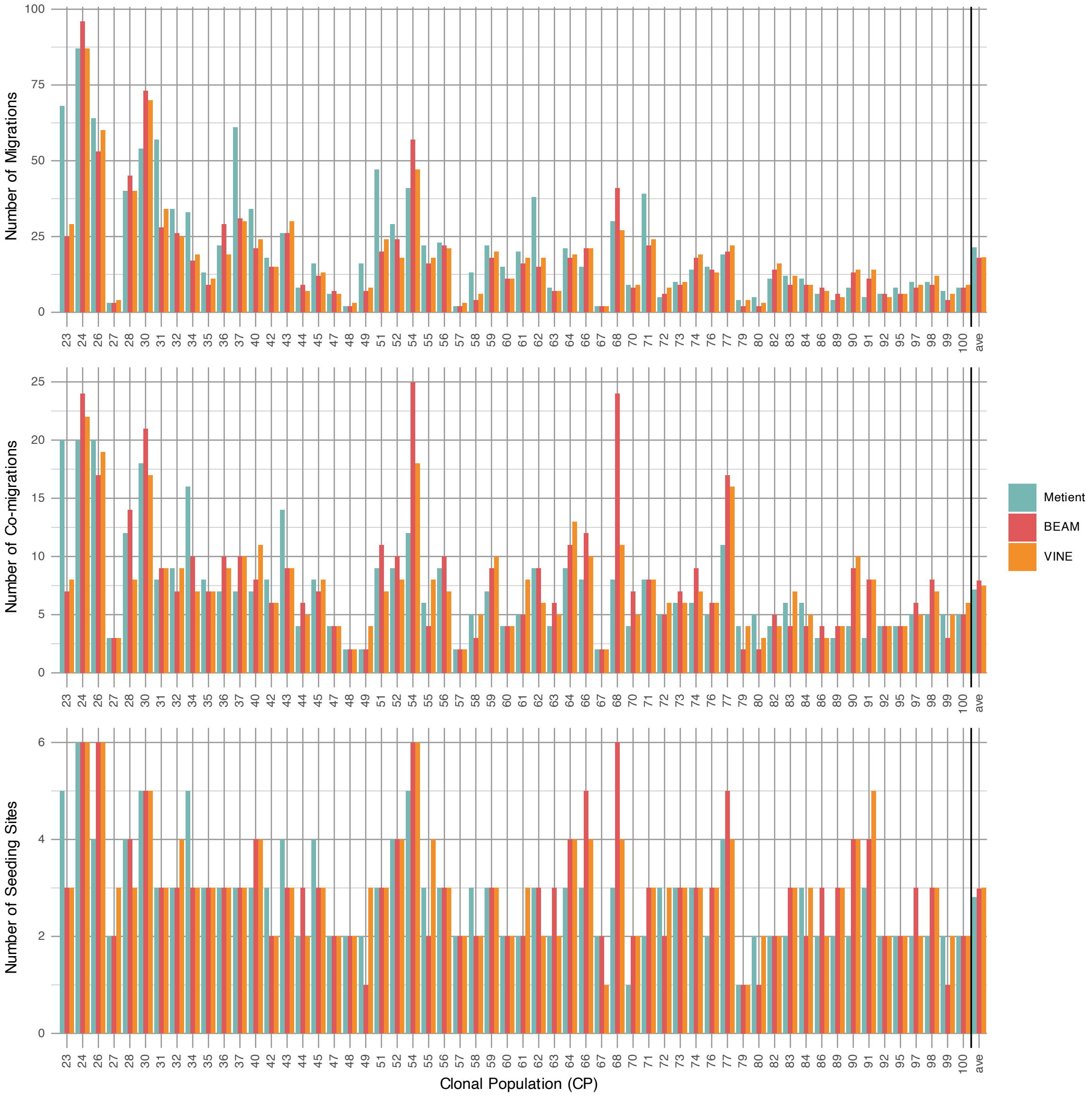


**Supplementary Figure S20:** Number of migrations, co-migration, and seeding sites for migration graphs inferred by Metient (*turquoise*), VINE (*orange*), and BEAM (*red*) for Clonal Populations (CPs) from ref. [9]. The largest CPs ($< 23$) were excluded here for reasons of computational efficiency. Results are shown for the $>0.5$ posterior-probability graph for each method.

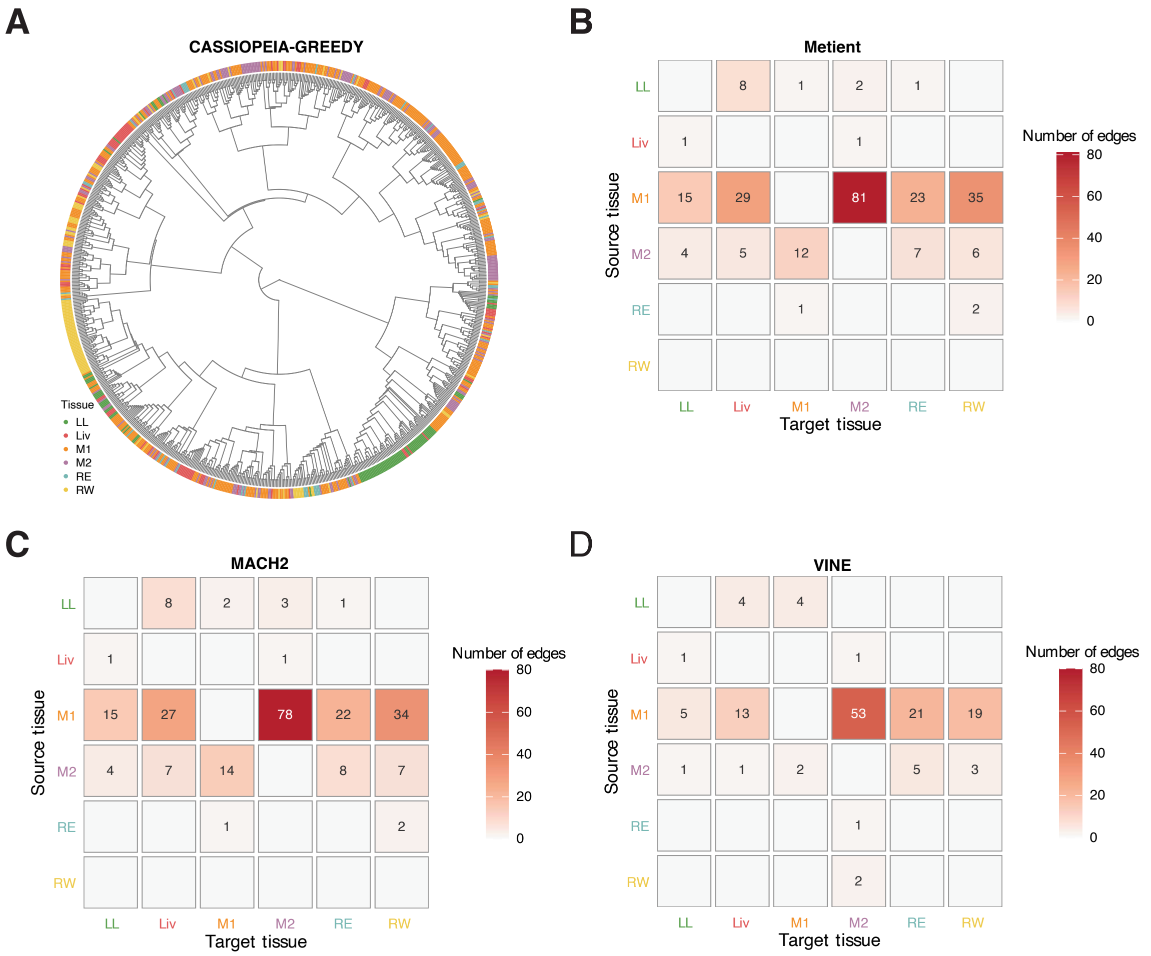


**Supplementary Figure S21:** **(A)** Tree inferred by Cassiopeia-Greedy for Clonal Population (CP) 4 from ref. [9] with colors at tips indicating observed tissues. **(B–D)** Matrix representations of the multi-edge migration graphs inferred for CP4 by (**B**) Metient and Cassiopeia-Greedy, (**C**) MACH2 and Cassiopeia-Greedy, and (**D**) VINE. LL, left lung; Liv, liver; M1, mediastinum 1; M2, mediastinum 2; RE, right lung E; RW, right lung W.

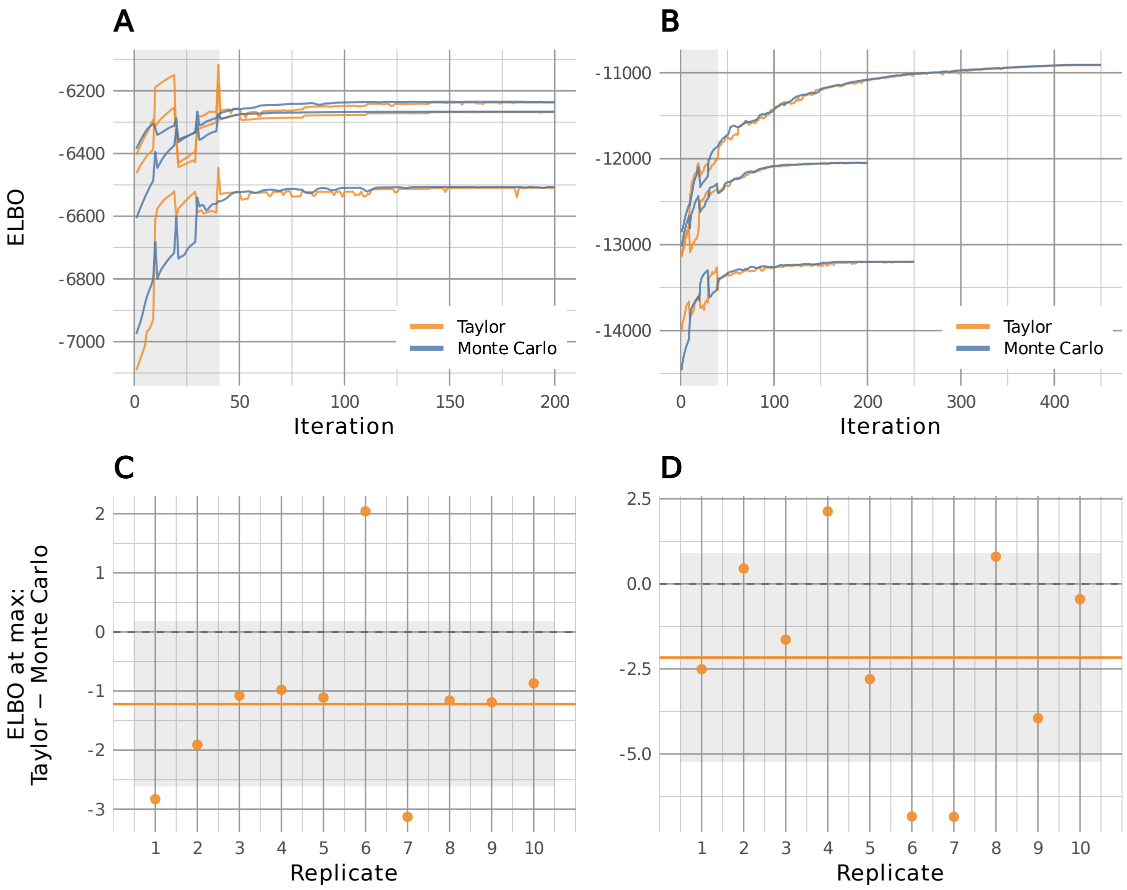


**Supplementary Figure S22:** Comparison of Taylor approximation of ELBO with Monte Carlo sampling. **(A)** Convergence trajectories for VINE during variational inference by SGA for simulated data with 25 taxa (HKY model, 300 sites) under the Taylor approximation (*orange*) or 100-replicate Monte Carlo sampling (*blue*). Pairs of trajectories are shown for the first three of ten replicates. *Gray* indicates warm-up period, where subsampling of sites creates noisier estimates. **(B)** Similar trajectories for 50 taxa. **(C)** Differences in final estimates of ELBO for all ten replicates with 25 taxa. *Gray* background indicates one standard deviation around the mean (*orange* line). **(D)** Differences for 50 taxa. For both datasets, the Taylor approximation underestimated the ELBO by an average of only ∼0.025%. The corresponding parameter estimates were very close to one another.

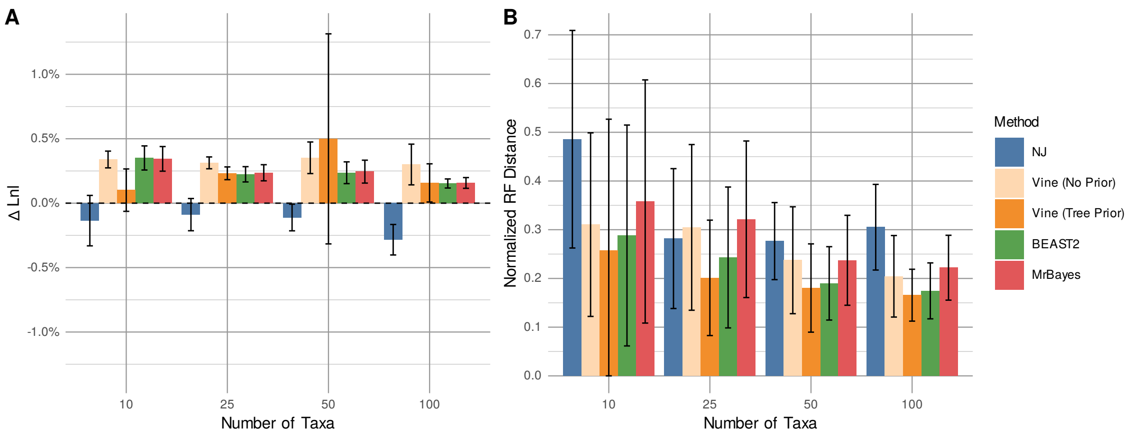


**Supplementary Figure S23:** Impact on VINE's performance of using an informative phylogenetic prior in place of the implicit prior. Results are shown for 300-site DNA alignments with simulation and inference under the HKY model. In the case of the informative prior, VINE was run with the –treeprior YULE and –relclock options. Results are shown for neighbor-joining (*blue*), VINE with the implicit prior (*light orange*) and with the informative prior (*orange*), BEAST 2 (*green*), and MrBayes (*red*). The prior had little effect in this setting. With an explicit prior, VINE optimizes a surrogate objective that substitutes the entropy of the embedding distribution for that of the induced tree distribution (see **Methods**).
